# Synaptic connectome of the *Drosophila* circadian clock

**DOI:** 10.1101/2023.09.11.557222

**Authors:** Nils Reinhard, Ayumi Fukuda, Giulia Manoli, Emilia Derksen, Aika Saito, Gabriel Möller, Manabu Sekiguchi, Dirk Rieger, Charlotte Helfrich-Förster, Taishi Yoshii, Meet Zandawala

## Abstract

The circadian clock and its output pathways play a pivotal role in optimizing daily processes. To obtain novel insights into how diverse rhythmic physiology and behaviors are orchestrated, we have generated the first comprehensive connectivity map of an animal circadian clock using the *Drosophila* FlyWire brain connectome. Intriguingly, we identified additional dorsal clock neurons, thus showing that the *Drosophila* circadian network contains ∼240 instead of 150 neurons. We also revealed extensive contralateral synaptic connectivity within the network and discovered novel indirect light input pathways to the clock neurons. Interestingly, we observed sparse monosynaptic connectivity between clock neurons and down-stream higher-order brain centers and neurosecretory cells known to regulate behavior and physiology. Therefore, we integrated single-cell transcriptomics and receptor mapping to decipher putative paracrine peptidergic signaling by clock neurons. Our analyses identified additional novel neuropeptides expressed in clock neurons and suggest that peptidergic signaling significantly enriches interconnectivity within the clock network.

## Introduction

Almost all living organisms from humans to bacteria possess a circadian clock (Dunlap, 1999). This internal timekeeping system enables organisms to anticipate and adapt to the rhythmic environmental changes that occur over a 24-hour cycle. At their molecular core, these clocks are comprised of cell-autonomous transcription-translation negative feed-back loops (Takahashi, 2017). In most animals, the master circadian clock in the brain receives light cues via the eyes which enable synchronization (or entrainment) with the external 24-hour light-dark cycles. The master clock sits at the top of the hierarchy and in turn modulates the activity of downstream neurons, as well as the peripheral clocks located in tissues throughout the body via endocrine and systemic signaling. In vertebrates, the master clock is located in the suprachiasmatic nucleus (SCN) of the hypothalamus and is comprised of approximately 20,000 neurons (Mohawk *et al*., 2012). Extensive intercellular coupling between these neurons likely via neurotransmitters, neuropeptides, and gap junctions forms a neuronal network that is resilient to internal and environmental perturbations (Maywood *et al*., 2006, Liu *et al*., 2007). Systematic characterization of these diverse coupling mechanisms between clock neurons is thus crucial to understanding circadian clock entrainment and the mutual coupling of the clock neurons. In addition, unraveling the clock output pathways which generate rhythmic behaviors and hormonal signaling can provide mechanistic insights into circadian regulation of organismal physiology and homeostasis.

These aims are particularly challenging to accomplish in vertebrates due to the large neuronal network size and resultant increase in complexity. However, the molecular clock architecture as well as the neuronal network motifs are highly conserved from humans to insects (Panda *et al*., 2002, Helfrich-Förster, 2004). Hence, *Drosophila melanogaster* with its powerful genetic toolkit and a complete brain connectome represents an ideal system for deciphering the clock network and its input and output pathways (Zheng *et al*., 2018, Dorkenwald *et al*., 2023, Schlegel *et al*., 2023). Not surprisingly, the fly circadian clock, generally regarded to be comprised of approximately 150 neurons, is extremely well characterized (Dubowy and Sehgal, 2017). These neurons have historically been classified into different groups of Lateral and Dorsal Neurons (LN and DN, respectively) based on their size, anatomy, location in the brain, and differences in gene expression (Abruzzi *et al*., 2017). Further, these neuronal subgroups are active at different times during the day and are consequently distinct functionally (Liang *et al*., 2016). Single-cell transcriptome sequencing analyses of clock neurons recently revealed additional heterogeneity within some of these clusters, which can largely be explained by neuronal signaling molecules that they express (Ma *et al*., 2021, Ma *et al*., 2023). Comprehensive mapping of the synaptic partners of clock neurons and comparing this connectivity can determine if the molecular heterogeneity based on gene expression also translates into heterogenous synaptic connectivity. Nonetheless, given the rich array of neuropeptides expressed in clock neurons, both synaptic and paracrine signaling appear crucial in mediating the connectivity between clock neurons as well as their output pathways. While recent work has begun to uncover some of this connectivity (Nagy *et al*., 2019, Barber *et al*., 2021, Reinhard *et al*., 2022a, Reinhard *et al*., 2022b, Shafer *et al*., 2022, Hidalgo *et al*., 2023), global analyses encompassing entire neuronal networks across both brain hemispheres are lacking.

Here, we harnessed the power of connectomics to generate the first comprehensive connectivity map of the circadian clock. Intriguingly, we identified additional DN in the network, thus showing that the *Drosophila* circadian clock is in fact comprised of at least 240 neurons instead of 150 neurons. In addition, our analyses revealed that light input from extrinsic photoreceptors to the clock neurons is largely indirect. Furthermore, we discovered extensive ipsilateral synaptic connectivity between the clock neurons and identified a subset of DN as an important hub that link the clock network across the two brain hemispheres via contralateral projections. We also elucidated the output pathways from the clock network that could affect general behavioral activity levels and organismal physiology. In particular, we characterized clock inputs to subsets of neurosecretory cells (NSC) in the brain which collectively represent a major neuroendocrine center (Nässel and Zandawala, 2020). We observed sparse monosynaptic connectivity between clock neurons and NSC, suggesting that multi-synaptic connections and peptidergic signaling account for most of the connectivity, as is characteristic of the vertebrate clock output pathways (Wen *et al*., 2020). Hence, as a complementary approach, we also deciphered putative paracrine signaling pathways within the clock network by extensively mapping the expression of clock neuropeptides and their receptors, and filtering them based on the topographical constraints determined by the connectome. This peptidergic signaling greatly enriches the connectivity within the clock network and is the major output pathway to the neuroendocrine center which regulates systemic physiology.

## Results

### Identification of the circadian neuronal network in the FlyWire and hemibrain connectomes

The master clock in the *Drosophila* brain is widely regarded to be comprised of approximately 150 neurons. This number is derived from neuronal expression of different clock genes (Ma *et al*., 2021). *Clk856-Gal4*, based on the *Clk* promoter, faithfully recapitulates expression in most of these clock neurons (Figure 1A). These neurons can be broadly classified into four classes each of LN and DN (Figure 1B and Table 1). The LN comprise Lateral Posterior Neurons (LPN), dorsoLateral Neurons (LN_d_), Ion Transport Peptide (ITP)-expressing LN (LN^ITP^), and Pigment-Dispersing Factor (PDF)-expressing ventroLateral Neurons (LN_v_^PDF^). Conversely, the DN include anterior Dorsal Neurons 1 (DN_1a_), posterior DN_1_ (DN_1p_), DN_2_ and DN_3_. These clock neuron classes can be further subdivided into different cell types (Figure 1C and Table 1). Morphological characterization of some DN_1p_ and DN_3_ subtypes is currently lacking (Reinhard *et al*., 2022b). Moreover, the estimated number of clock neurons in *Drosophila* is likely an underestimation as the precise number of DN_3_ has not been determined thus far (Shafer *et al*., 2006). This is partly because drivers like *Clk856-Gal4* only include a small proportion of DN_3_ (Ma *et al*., 2021). In addition, most DN_3_ have small somata that are densely packed together, making it difficult to count. Majority of these small DN_3_ project to the central brain, and hence they are aptly called small Central Projecting DN_3_ (s-CPDN_3_) (Table 1) (Sun *et al*., 2022). Similarly, a pair of DN_3_ with large somata project to the central brain (large Central Projecting DN_3_; l-CPDN_3_), whereas about 6 neurons per brain hemisphere project to the anterior brain (Anterior Projecting DN_3_; APDN_3_) (Sun *et al*., 2022).

**Table 1:**
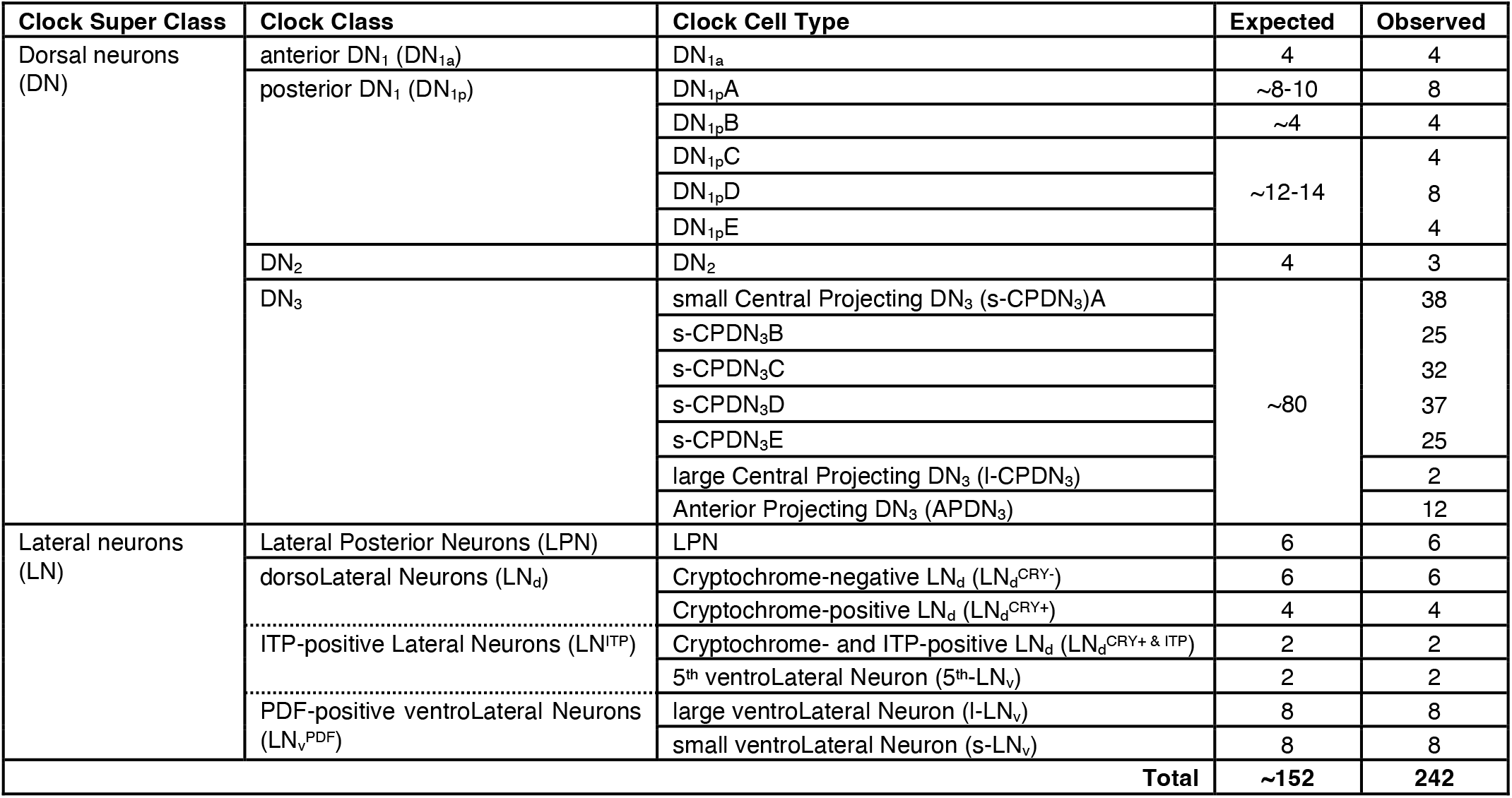
Identification and classification of *Drosophila* clock neurons in the FlyWire brain connectome.

**Figure 1:**
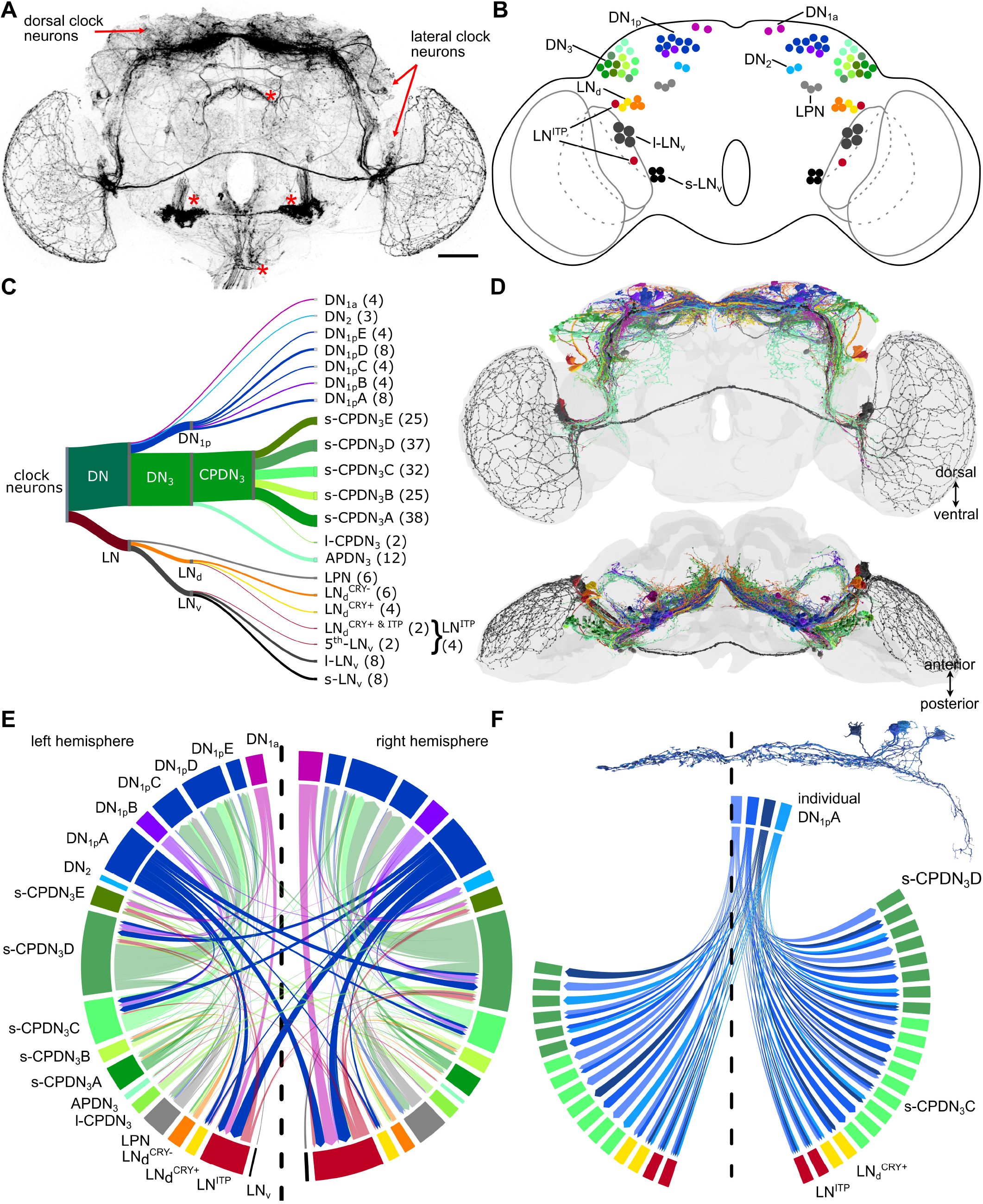
*Drosophila melanogaster* circadian clock network. **(A)** *Clk856-Gal4* drives GFP expression in most of the circadian clock neurons and some non-clock neurons (asterisk). Only a few DN_3_ are included in this Gal4-line. Scale bar = 50μm. **(B)** *Drosophila* clock neurons can be divided into four classes each of Dorsal neurons (DN) and Lateral neurons (LN) based on their cell body location. These can be further subdivided into different cell types based on their morphology and gene expression. **(C)** Classification and numbers of all clock neurons identified in the FlyWire connectome. Refer to Table 1 for details. **(D)** The morphology of identified clock neurons in the connectome largely resembles the morphology of clock neurons marked by *Clk856-Gal4*. **(E)** Broad synaptic interconnectivity between different cell types within the clock network. The direction of the arrow indicates the flow of information. Strong connectivity is observed from DN_1a_ to LN^ITP^, from s-CPDN_3_C,D to DN_1p_C-E, as well as from DN_1p_A to LN^ITP^, LN_d_^CRY+^, s-CPDN_3_C and s-CPDN_3_D. Note extensive contralateral connectivity between different cell types, most of which can be accounted for by the DN_1p_A. Dashed line indicates the brain midline. **(F)** Individual DN_1p_A in the right hemisphere form both ipsilateral and contralateral connections with s-CPDN_3_C, s-CPDN_3_D, LN_d_^CRY+^ and LN^ITP^.

These latter cells also have larger somata than the s-CPDN_3_ (Reinhard *et al*., 2022b). Despite the large number of DN_3_, previous studies suggest that they are less important for behavioral rhythmicity compared to some of the LN, which are regarded as the master pacemaker neurons (Rieger *et al*., 2006, Yoshii *et al*., 2012, Li *et al*., 2018).

As a first step in determining the synaptic connectivity within the clock network, we used a systematic approach (Figure S1) to identify most, if not all, of the clock neurons in the FlyWire connectome generated in our companion papers (Dorkenwald *et al*., 2023, Schlegel *et al*., 2023). In total, we successfully identified 242 clock neurons based on a combination of morphology, previously determined connectivity, and location of their cell soma (Figure 1C-D, Supplementary Videos 1-2) (Schubert *et al*., 2018, Reinhard *et al*., 2022a, Reinhard *et al*., 2022b, Shafer *et al*., 2022, Sun *et al*., 2022). The number of clock neurons that we identified in the connectome was considerably higher than the expected number of clock neurons (∼152) in the adult brain (Table 1). This was mainly due to the presence of more s-CPDN_3_ in the connectome than estimated, which prompted us to accurately quantify the total number of DN_3_ in adult brains. To comprehensively quantify DN_3_, we used antibodies against clock proteins Period (PER) and Vrille (VRI) in *tim-Gal4* > GFP expressing flies. Since PER and VRI exhibit peak expression at different Zeitgeber times (Figure 2A), the inclusion of three clock markers (PER, VRI, and *tim*) can provide a reliable estimate of the total number of DN_3_ (Figure 2B and C, Supplementary Video 3). Indeed, our analyses revealed that there are more than 70 DN_3_ per hemisphere in both males and females (Figure 2D). Thus, the number of DN_3_ identified in the connectome (171 neurons) is in line with our DN_3_ estimate (∼166 neurons) based on anatomical analysis. The majority of these are s-CPDN_3_ (157 neurons) which we now classified into five subtypes (s-CPDN_3_A-E) based on morphological similarity (Figure 2E-I). These five DN_3_ subtypes are comprised of 26 distinct cell types (Schlegel *et al*., 2023) and could thus be subdivided further in the future.

**Figure 2:**
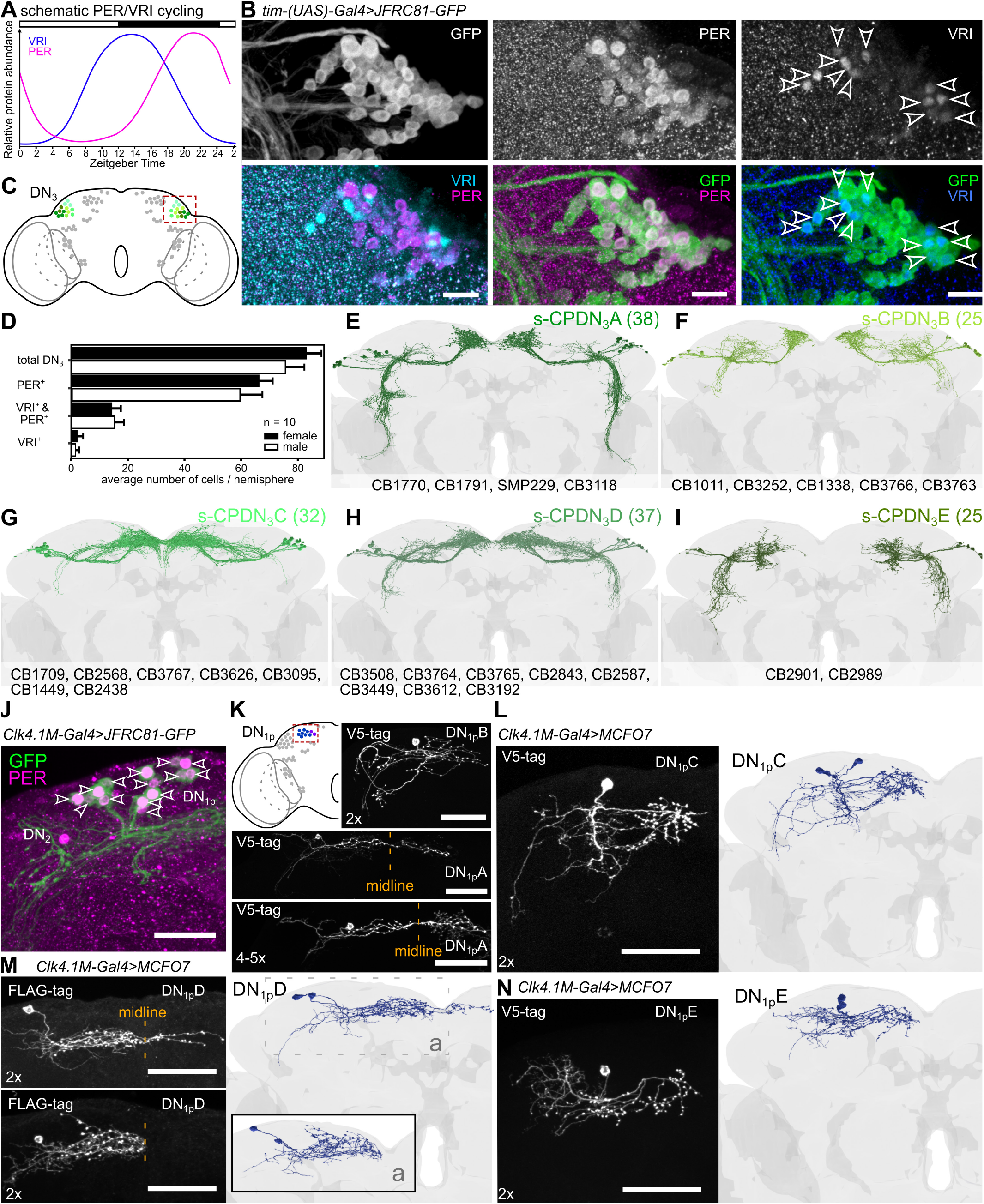
Morphology of newly identified s-CPDN_3_ and DN_1p_ subtypes. **(A)** Schematic cycling of Period (PER) and Vrille (VRI) protein abundance in clock neurons (Glossop *et al*., 2003). **(B)** *tim-(UAS)-Gal4* drives GFP expression in all DN_3_, many of which coexpress PER and about 12-19 coexpress VRI. When VRI staining is strong, PER staining is weak or not detectable. Image based on a female brain fixed at ZT1. Scale bar = 10μm **(C)** DN_3_ are closely associated with the lateral horn. **(D)** The number of DN_3_ per hemisphere accounts for s-CPDN_3_, l-CPDN_3_, and APDN_3_ identified in the connectome. On average, females have slightly more DN_3_ (83±5 standard deviation) compared to males (76±6 standard deviation), which could be attributed to DN_3_ being more densely packed (and thus difficult to quantify) in males. n = 10 hemispheres from 5 brains. Error bars depict standard deviation. **(E-I)** s-CPDN_3_ identified in the FlyWire dataset. Numbers in brackets represent numbers of neurons in the total brain. s-CPDN_3_ could be subdivided into five subtypes. Each subtype comprises of two to eight different cell types that have unique morphological characteristics. **(J)** *Clk4*.*1M-Gal4* reliably drives GFP expression in 14-15 DN_1p_ per hemisphere. **(K)** Multi-color flip-out (MCFO) analysis reveals previously characterized DN_1p_A (4-5 neurons/hemisphere) and DN_1p_B (2 neurons) subtypes. **(L-N)** MCFO analysis additionally reveals the morphology of uncharacterized DN_1p_ subtypes. DN_1p_C and DN_1p_E comprise two cells each per hemisphere (L, N). DN_1p_D comprises four cells per hemisphere two of which project over the midline while the other two do not, suggesting that the DN_1p_D is comprised of two subtypes (M). Scale bar = 50μm.

Additionally, we also identified several candidate DN_1p_ which could not be classified into previously characterized DN_1p_ cell types (Table 1). To verify if these neurons identified in the connectome are indeed DN_1p_, we characterized them anatomically. Specifically, we performed multi-color flip out (MCFO) analysis of DN_1p_ using *Clk4*.*1M-Gal4* to elucidate the morphology of this densely packed neuronal cluster. In total, about 14 DN_1p_ per hemisphere are labeled with the *Clk4*.*1M-Gal4* (Figure 2J). MCFO analysis revealed DN_1p_A (4-5 neurons) and DN_1p_B (2 neurons) subtypes characterized previously (Table 1 and Figure 2K). Our analysis additionally revealed the morphology of previously uncharacterized DN_1p_ subtypes (Figure 2L-N). DN_1p_C and DN_1p_E are each comprised of two neurons per hemisphere (Figure 2L and N). DN_1p_D is comprised of four neurons per hemisphere (Figure 2M). Two of these project over the midline while the other two remain ipsilateral, suggesting that the DN_1p_D is comprised of two morphologically distinct subtypes. Notably, our analysis confirmed that the candidate DN_1p_ identified from the connectome are in fact morphologically similar to DN_1p_C-E subtypes labeled by *Clk4*.*1M-Gal4*. Taken together, our identification and morphological characterization of novel DN_3_ and DN_1p_ subtypes provide a solid framework to comprehensively examine the connectivity of clock neurons.

The FlyWire connectome combines automatically detected chemical synapses with proofread neurons. These synapses represent an additional anatomical feature that could potentially distinguish neuronal groups. Consequently, we asked whether the classification of clock neurons based on differences in their synaptic connectivity aligns with the traditional anatomical and recent gene expression-based classification. To address this, we clustered clock neurons based on cosine similarity between their total synaptic inputs and outputs. Our clustering analysis shows that neurons of a given clock cell type (Table 1) usually cluster together, suggesting that neurons from the same group are more similar (in terms of synaptic connectivity) to each other than to other clock neurons (Figure S2A-B). For example, all three LN_d_^CRY-^ from one hemisphere are part of the same clade. Similarly, the two DN_2_ are part of a clade. Exceptions to this are the clades containing DN_1p_ and DN_3_. For DN_1p_, this can be explained by our findings which show that this group comprises five morphologically distinct subtypes. In the case of DN_3_, while some of them form their own cluster, other DN_3_ cluster together with different clock neuron subtypes. On one hand, this is not unexpected since it is unlikely for such a large group of neurons to have similar connectivity patterns. On the other hand, this is quite interesting since it provides insights into their possible function. For instance, heterogenous clusters comprising clock neurons of different classes have similar synaptic inputs and outputs, and may thus play similar roles in the clock network and beyond. Moreover, the connectivity-based clustering does not resolve the five subtypes of the s-CPDN_3_. This suggests that these five s-CPDN_3_ subtypes comprise synaptically heterogeneous cell populations. Nonetheless, synaptic connectivity-based classification of clock neurons largely aligns with the ones determined based on anatomical and gene expression differences.

Having identified all the clock neurons, we next sought to determine their synaptic interconnectivity which could facilitate intercellular coupling within the network. Generally, we regarded >4 common synapses per neuron as significant connections and >9 synapses as strong connections. We first wanted to validate our analysis by comparing it with previously reported connections. In agreement with previous reports (Reinhard *et al*., 2022b, Shafer *et al*., 2022), we observed strong synaptic connectivity from DN_1a_ to LN^ITP^, from DN_1p_A to LN^ITP^ and LN_d_, and from DN_1p_B to DN_2_ clusters (Figure 1E and S3), highlighting the robustness of our approach. Importantly, our analysis also uncovered novel connections between the different subgroups. Specifically, we observed strong contralateral and ipsilateral connectivity from DN_1p_A to LN^ITP^, as well as additional significant connections with s-CPDN_3_C, s-CPDN_3_D, and LN_d_^CRY+^ across both hemispheres. Similarly, LN^ITP^ also provide synaptic inputs to s-CPDN_3_ clusters in both hemispheres (Figure 1E and S3). This raised the question of whether DN_1p_A represents a heterogeneous population where one subgroup forms ipsilateral connections and the other contralateral. To address this, we examined the connectivity at cellular resolution (Figure S4-S5) which revealed that individual DN_1p_A indeed form both ipsilateral and contralateral connections (Figure 1F). Our analysis thus identified DN_1p_A as an important center which links the clock network across the two brain hemispheres. These results are in line with previous reports of contralateral projections of DN_1p_ (Lamaze *et al*., 2018). In contrast, there are virtually no synaptic connections between the s-LN_v_ and the LN_d_ neurons (Figure 1E and S3-S5), which control morning and evening activities, respectively. This is consistent with previous analyses using the hemibrain connectome (Shafer *et al*., 2022). Interestingly, the connectivity of clock neurons across both hemispheres is not symmetrical, owing to the differences in the number of synapses (Figure 1E and S3).

To assess the extent of inter-individual differences in the numbers, neuronal projections, and synaptic connectivity of clock neurons, we next performed comparisons with the partial hemibrain connectome (Scheffer *et al*., 2020). Several groups of clock neurons were previously identified in the hemibrain connectome including all s-LN_v_, l-LN_v_, LN_d_, LN^ITP^, LPN, DN_1a_, and some DN_2_, DN_1p_, and DN_3_ (Reinhard *et al*., 2022a, Reinhard *et al*., 2022b, Shafer *et al*., 2022). Here, we identified additional DN_1p_ and DN_3_ (Figure S6A-D). In total, 64 clock neurons can be identified in the hemibrain connectome, with the majority of missing neurons belonging to the DN_3_ subgroups (Figure S6A-D, Supplementary Table 5). Comparison of different subgroups revealed stable neuronal numbers across the two connectomes (Table 1 and Figure S6D). Similarly, there is a high degree of stereotypy in the connectivity between the clock clusters (Figure S6E-F). For instance, l-LN_v_ and DN_2_ form the least synaptic contacts with other clock clusters. At the opposite end of the spectrum, s-CPDN_3_A are connected to all the clock clusters except for s-LN_v_, l-LN_v_, DN_1a_, and DN_1p_E. Given its partial nature, the hemibrain connectome lacks information about all contralateral connections, reiterating the significance of characterizing information flow across entire networks. Taken together, our analyses revealed hitherto unknown connectivity between the clock neurons which could contribute to the robustness of the master clock. Moreover, the identification of the complete circadian neuronal network in the FlyWire connectome underscores the power of the fruit fly in pushing forward the frontier of our understanding of chronobiology.

### Validating clock neuron connectivity using trans-synaptic tracing

While the FlyWire and hemibrain connectomes exhibit a high degree of stereotypy, we further used an independent approach to validate our connectivity analyses. We performed light microscopy-based trans-synaptic circuit tracing by expressing *trans*-Tango (Talay *et al*., 2017) using specific driver lines for different populations of clock neurons (Figure S7-S8). Upon driving *trans*-Tango with *Clk4*.*1M-Gal4* which labels most DN_1p_ (Figure S7B), we observed post-synaptic signals in DN_1p_, DN_2_, DN_3_, LPN, LN_d_, 5^th^-LN_v_, and s-LN_v_ (Figure S9A). However, l-LN_v_ and DN_1a_ were not post-synaptic to DN_1p_. Thus, our *trans*-Tango analysis of DN_1p_ agrees with the connectivity of DN_1p_ based on the connectomes (Figure S3 and S6F). Similarly, a split-Gal4 line targeting DN_3_ drives post-synaptic signals in DN_1a_, DN_1p_, DN_2_, DN_3_, LPN, LN_d_, and l-LN_v_ (Figure S9B), which mirrors the connectivity seen in the connectomes. While post-synaptic signal was not detected in most clock neurons of control flies, occasionally, a false post-synaptic signal was detected in two clock neurons (LN_d_) from the entire network (Figure S10A). Hence, any potential synaptic output to LN_d_ should be treated with caution. Overall, we observed similar congruency between the two approaches with other Gal4-lines including those targeting DN_2_ (Figure S10B), LPN (Figure S10C), and LN^ITP^ (Figure S10D).

Differences compared to the connectomes were observed when driving *trans*-Tango with *Pdf-Gal4* (for s-LN_v_ and l-LN_v_) (Figure S10E) and with the DN_1a_-specific split-Gal4 line (Figure S10F). In both cases, *trans-*Tango generated post-synaptic signals in more clock neurons than anticipated based on the connectomes. This discrepancy could be explained by: 1) the presence of additional neurons in the Gal4 (e.g. PDF tritocerebrum neurons (Selcho *et al*., 2018)) and/or 2) daily remodeling of neural circuits, as shown previously for s-LN_v_ and DN_1a_ (Fernandez *et al*., 2008, Song *et al*., 2021). In summary, our *trans-*Tango analysis is largely in agreement with the clock network generated using the connectomes.

### Deciphering light input pathways via in-silico retrograde tracing of clock neurons

Following the successful validation of our connectivity data, we next identified all the major classes of neurons providing inputs to the clock network. To this end, we utilized the annotation scheme of our companion paper (Schlegel *et al*., 2023), which provides a hierarchical classification of all neurons in the connectome (Figure 3A). We found that neurons intrinsic to the brain provide the majority of the inputs to the clock network (Figure 3A-C). This includes visual centrifugal neurons projecting from the central brain to the optic lobes, visual projection neurons projecting from the optic lobes to the central brain, as well neurons intrinsic to the optic lobes and central brain (Figure 3C). Examining inputs to specific clock clusters, we observed differential inputs across all the subgroups (Figure 3B). As expected, s-LN_v_ and l-LN_v_ receive most of their input from optic lobe and visual centrifugal neurons as they have a large number of input sites in the optic lobes and the accessory medulla (AME) (Figure S11). In contrast, APDN_3_, l-CPDN_3_, and LN^ITP^ populations receive major inputs from visual projection neurons. The remaining clock clusters receive most of their inputs from central brain neurons (Figure 3B, S11, and S12). In some cases, such as DN_1p_C-E, DN_2_, and s-CPDN_3_A-E, a significant portion of these central neurons are clock neurons themselves, confirming prominent intercellular synaptic connectivity between some clock clusters. Interestingly, only 4 sensory neurons provide direct inputs to the clock network. These are anterior cells (aDT4) (Dr. Gregory Jefferis, personal communication) (Figure 3A and D) which provide temperature inputs to LPN, DN_1p_C, and DN_1p_E (Jin *et al*., 2021, Alpert *et al*., 2022).

**Figure 3:**
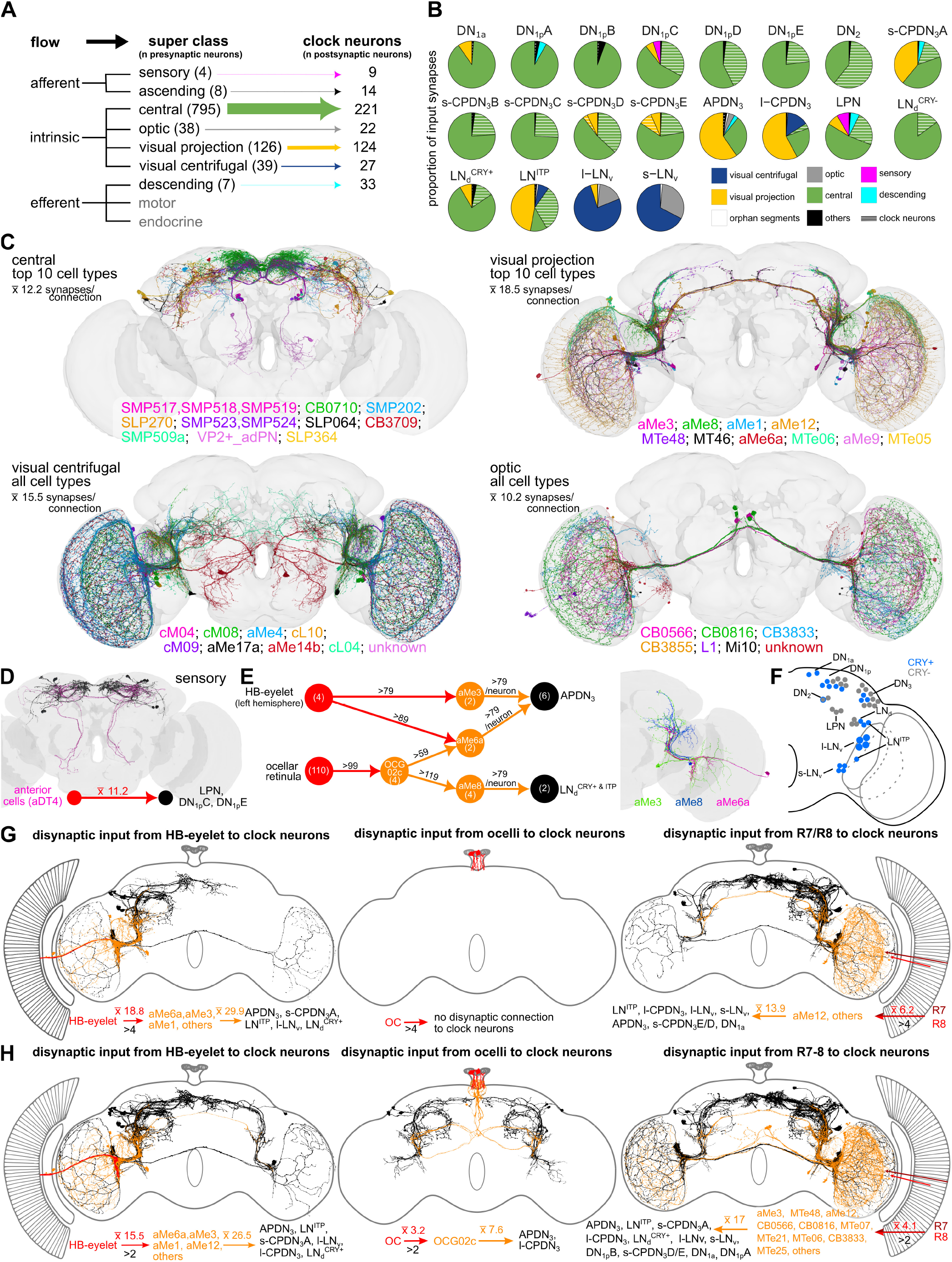
Light and other inputs to the clock network. **(A)** Input to the clock neurons grouped by the nine neuronal super classes annotated in the FlyWire connectome. Intrinsic neurons, specifically the central and visual projection neurons, are the largest groups providing inputs to the clock. **(B)** Breakdown of inputs to different types of clock neurons. **(C)** Neurons from the four different super classes (central, visual projection, visual centrifugal and optic) which provide major inputs (based on number of neurons) to the clock. **(D)** Anterior cells (AC, magenta) providing temperature inputs to LPN, DN_1p_C, and DN_1p_E. (black). **(E)** aMe neurons provide the strongest inputs to clock neurons. Upstream partners of aMe neurons with strong connectivity include HB-eyelet and OCG02c. Numbers above the arrows indicate the number of synapses and numbers in circles indicate the number of neurons. Neuronal reconstructions of aMe3, aMe6a and aMe8 are shown in one hemisphere. **(F)** Cryptochrome (CRY)-positive clock neurons are shown in blue. **(G)** Disynaptic inputs to the clock from the three types of extrinsic photoreceptors (HB-eyelet, ocelli, photoreceptor cells of the compound eye) using a threshold of >4 synapses. **(H)** Disynaptic inputs to the clock from the three types of extrinsic photoreceptors using a threshold of >2 synapses. For C, D, G and H, numbers represent the average number of synapses. All cell types are listed according to their input strength (from high to low) to clock neurons. Note: HB eyelet was only identified in the left hemisphere. Abbreviations: HB, Hofbauer-Buchner; OC, ocelli.

Having broadly classified the inputs from different neuronal super classes to the clock network, we probed further and identified individual cells providing the strongest synaptic inputs to clock neurons. For this purpose, we used a stringent threshold of 80 synapses to obtain a narrow list of candidate inputs. Our analysis discovered 13 neurons, including 7 aMe neurons (a subset of visual projection and visual centrifugal neurons) that are strongly connected to specific downstream clock neurons (Figure 3E). For example, individual aMe3 and aMe6a neurons can form more than 79 synapses with APDN_3_, while aMe8 are similarly connected to LN_d_^CRY+ & ITP^ clock neurons (Figure 3E). The unifying feature of these aMe neurons is their dense arborization in the AME and posterior lateral protocerebrum (Figure 3E), where they anatomically interact with clock neuron dendrites (Figure S11 and S12) (Reinhard *et al*., 2022b, Tang *et al*., 2022). Interestingly, the aMe neurons themselves receive strong inputs from the extraretinal photoreceptors. Specifically, aMe3 and aMe6a neurons receive strong inputs directly from the Hofbauer-Buchner (HB) eyelets (Figure 3E). Conversely, aMe8 receive indirect inputs from ocellar retinula cells via the ocellar ganglion neurons (OCG) type 2c (OCG02c) (Figure 3E). This suggests that the clock receives strong light inputs from extrinsic photoreceptor cells, albeit indirectly. This is not surprising since light is the most important Zeitgeber for circadian clocks (Helfrich-Förster, 2020). Flies synchronize their circadian clocks with the light-dark cycles using these extrinsic photoreceptor cells as well as via the blue-light photoreceptor Cryptochrome (CRY), which is expressed in about half of the clock neurons (Figure 3F) (Helfrich-Förster, 2020). While CRY interacts with the core clock protein Timeless and can quickly reset the clock, the different photoreceptor cells are important for sensing dawn, dusk, high light intensities, and day length, and for adapting morning and evening activities to the appropriate time of day. Regardless, we found little direct inputs from the photoreceptor cells and other sensory cells to the clock neurons (Figure 3A). This is consistent with previous findings which revealed that most of the light input to the clock appears to be indirect (Alejevski *et al*., 2019). In light of this and our *in-silico* circuit tracing analysis described above, we comprehensively characterized indirect connectivity between photoreceptor cells and clock neurons. Therefore, we traced all the disynaptic connections between them. Using the normal threshold of >4 synapses, we again recovered the strong connections from the H-B eyelets via the aMe3/aMe6a to the APDN_3_, and additional weaker connections to the s-CPDN_3_A, LN^ITP^, l-LN_v_, and LN_d_^CRY+^ (Figure 3G). Furthermore, we revealed connections from R7/R8 compound eye photoreceptors to several clock neurons via aMe12 (Kind *et al*., 2021) and other interneurons. While we did not observe any disynaptic connections from the ocellar retinula cells to clock neurons using the normal threshold (Figure 3G), reducing the threshold to >2 synapses revealed connections from the ocelli to APDN_3_ and l-CPDN_3_ via OCG02c (Figure 3H). The synaptic connections from the ocelli to OCG and beyond are extensively characterized in our companion paper and demonstrate interesting details that may also be valid for the other photoreceptor inputs to clock neurons (Dorkenwald *et al*., 2023). The majority of ocellar photoreceptors are synaptically connected to ocellar ganglion neurons with thick axons (OCG01a-f) or directly to descending neurons (DNp28). These connections likely enable fast behavioral responses. In contrast, axons of OCG02c that connect to the clock neurons are rather thin and not suited for fast neurotransmission. Instead, these neurons appear suited for collecting light information over time – a property needed for entraining the circadian clock. Further, collecting light information over larger time intervals may not require a high synapse density. Thus, 3 to 4 synapses between retinula cells and the relevant downstream OCG observed here could be sufficient for this purpose (Figure 3H). The same is also true for the photoreceptor cells of the compound eyes. Reducing the threshold of significant connections from 5 to 3 synapses revealed indirect clock input from additional photoreceptor cells, including those that project from the dorsal rim area of the eye (Figure 3H). These photoreceptor cells are involved in polarized vision and might contribute to time-compensated sun compass orientation (Homberg, 2004). Whether the connectivity observed with a lower threshold of >2 synapses is functional *in vivo* remains to be seen; however, this is very likely since there are usually many photoreceptor cells that synapse onto only a few aMe neurons. For example, theoretically, the ∼300 pale R8 cells project to only 3-4 aMe12 neurons (Kind *et al*., 2021), resulting in ∼100 connections on average per aMe neuron. Even if each of these connections were mediated via only 3 synapses, each aMe neuron could potentially receive inputs from R8 cells via 300 synapses, which is quite substantial (Figure 3H).

### In-silico anterograde tracing of clock neurons

Delineating the output pathways that translate daily 24-hour oscillations of the molecular clock into physiological and behavioral rhythms remains a major focus in chronobiology. Using the same strategy as above to identify the inputs, we systematically classified all the neurons downstream of the clock network. Most synaptic output from the clock network is directed to intrinsic brain neurons, and in particular, the central brain neurons (Figure 4A-C). Except for l-LN_v_, all clock clusters have a majority of their output onto central brain neurons. l-LN_v_ mostly provide inputs to Medullary intrinsic neurons in the optic lobe (Mi 1, Figure 4B-C), consistent with their role in adapting the sensitivity of the visual system to the time of day (Chen *et al*., 1992, Pyza and Meinertzhagen, 1997). Further, the majority of the output from DN_1p_A is onto visual projecting and central brain neurons that are part of the clock network (LN^ITP^, s-CPDN_3_C/D, Figure 4B). After broadly classifying clock outputs, we next focused on specific cell types which receive the strongest synaptic inputs (>49 synapses) from clock neurons using an approach similar to the one used earlier for clock inputs. Our analysis identified the enigmatic Clamp neurons (Figure 4D), which receive strong synaptic inputs from APDN_3_. While the functions of most of these clamp neurons are still unknown, some of them output onto descending neurons, while others promote sleep (Sun *et al*., 2022). Moreover, DN_1p_B provide strong inputs to Tubercle-innervating neurons (Figure 4E), which are part of the anterior visual pathway (Hulse *et al*., 2021). Lastly, several clock neurons are strongly connected to diverse neurons from different neuropil regions (Figure 4F).

**Figure 4:**
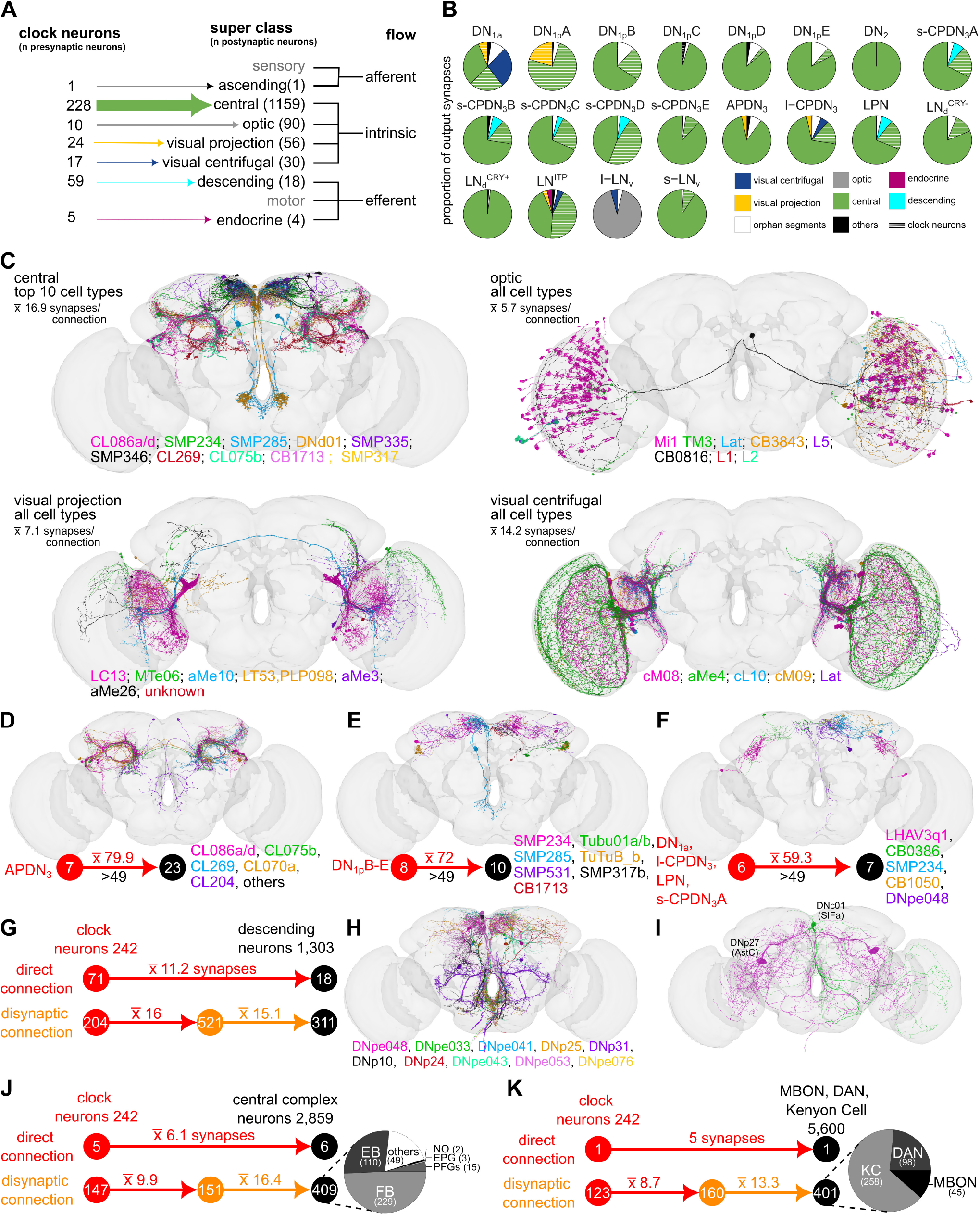
Direct and indirect clock output pathways. **(A)** Clock output grouped by the nine neuronal super classes annotated in the FlyWire connectome. Intrinsic neurons, specifically the central neurons, are the largest group of neurons downstream of clock neurons. **(B)** Breakdown of outputs from different types of clock neurons. All clock neurons, except for l-LN_v_, provide a majority of their output to central neurons. l-LN_v_ mostly provide inputs to optic neurons. **(C)** Neurons from the four different super classes (central, optic, visual projection and visual centrifugal) which receive inputs from the clock. **(D)** APDN_3_ form the most output synapses in the clock network and are highly connected to several types of Clamp neurons (CL). **(E)** DN_1p_B-E provide strong outputs to Tubercle-innervating neurons. **(F)** Several clock neurons provide strong inputs to neurons from diverse neuropils. **(G)** Mono- and di-synaptic clock inputs to descending neurons. **(H)** Novel and **(I)** peptidergic descending neurons receiving direct clock input. Mono- and di-synaptic clock inputs to neurons associated with the **(J)** central complex and **(K)** mushroom bodies. Note that there are only a few direct connections from the clock to downstream higher brain centers. Numbers above the arrows indicate the number of synapses and numbers in circles indicate the number of neurons. Note: all numbers refer to neurons across both hemispheres. For C-I, cell types are listed according to the strength of the clock input (from high to low). Abbreviations: EB, ellipsoid body; FB, fan-shaped body; NO, noduli; EPG, ellipsoid body-protocerebral bridge-gall neuron; PFGs, protocerebral bridge glomerulus-fan-shaped body-ventral gall surround; KC, Kenyon cell; DAN, dopaminergic neuron; MBON, mushroom body output neuron.

Next, we examined clock inputs to descending neurons which could influence locomotor and other behaviors regulated by neurons in the ventral nerve cord. Interestingly, clock neurons provide direct inputs to 18 descending neurons (Figure 4A and G). These descending neurons include those which have not yet been classified (Figure 4H), as well as Allatostatin-C (AstC) and SIFamine (SIFa) peptidergic neurons (Figure 4I), the latter of which modulate feeding, mating, and sleep (Nässel and Zandawala, 2022). Direct synaptic inputs to downstream descending neurons predominantly derive from s-CPDN_3_A-D, and LPN (Figure 4B). When considering disynaptic connections, the connectivity between the clock network and descending neurons increased drastically, with approximately 24% of all descending neurons receiving indirect inputs from most of the clock neurons (Figure 4G). In summary, our connectivity analysis indicates that the clock can have a major influence on diverse behaviors, including locomotion, via outputs to descending neurons.

In addition, the circadian clock is also known to modulate behaviors such as activity/sleep, spatial orientation, and learning and memory (Beer and Helfrich-Förster, 2020, Flyer-Adams *et al*., 2020). These behaviors are regulated by higher brain centers such as the central complex and mushroom bodies. Consistent with previous results (Reinhard *et al*., 2022a, Reinhard *et al*., 2022b), we found few direct connections from the clock to neurons associated with the central complex (Figure 4J) and mushroom bodies (Figure 4K). Consequently, we predicted that the clock output to these higher coordination centers is either indirect or paracrine via neuropeptides. In line with this prediction, we found prominent disynaptic connectivity between clock neurons and central complex neurons (mainly fan-shaped body (FB) and ellipsoid body neurons (EB)), as well as between clock neurons and Kenyon cells (KC), dopaminergic neurons (DANs) and mushroom body output neurons (MBONs) (Figure 4J-K). In support of paracrine signaling, receptors for several clock neuropeptides are also enriched in higher brain centers (Nässel and Zandawala, 2019). Taken together, circadian modulation of neurons regulating diverse behaviors is largely indirect or paracrine. Similarly, we observed very few direct connections between clock neurons and endocrine cells which influence organismal physiology and systemic homeostasis (Figure 4A-B). We address this connectivity in more detail below.

### Identification and characterization of the neuroendocrine center in the FlyWire connectome

While recent work has unraveled some clock output pathways to endocrine cells, our collective understanding of the circadian regulation of endocrine rhythms is poor (Nagy *et al*., 2019, Barber *et al*., 2021, Hidalgo *et al*., 2023). To address this knowledge gap, we first identified and classified all endocrine or NSC in the brain which are a major source of circulating hormones. These endocrine cells can be broadly classified into lateral, medial, and subesophageal zone NSC (l-NSC, m-NSC, and SEZ-NSC, respectively) based on their location in the brain. Their axons exit the brain via a pair of nerves (nervii corpora cardiaca, NCC), and depending on the cell type, innervate the corpora cardiaca, corpora allata, hypocerebral ganglion, crop, aorta, or the anterior midgut (Figure 5A). Their axon terminals form neurohemal sites through which hormones are released into the circulation or locally on peripheral targets such as the crop. Collectively, the NSC form a major, yet distributed, neuroendocrine center that is functionally analogous to the hypothalamus (Nässel and Zandawala, 2020). We identified all brain NSC in the FlyWire connectome by isolating the nerve bundle containing their axons (Figure 5B-C). In total, we independently identified 80 brain NSC (Supplementary Video 4), in agreement with our companion studies (Dorkenwald *et al*., 2023, Schlegel *et al*., 2023). We propose and utilize a systematic nomenclature for all brain NSC based on their location and neuropeptide identity (Table 2).

**Table 2:**
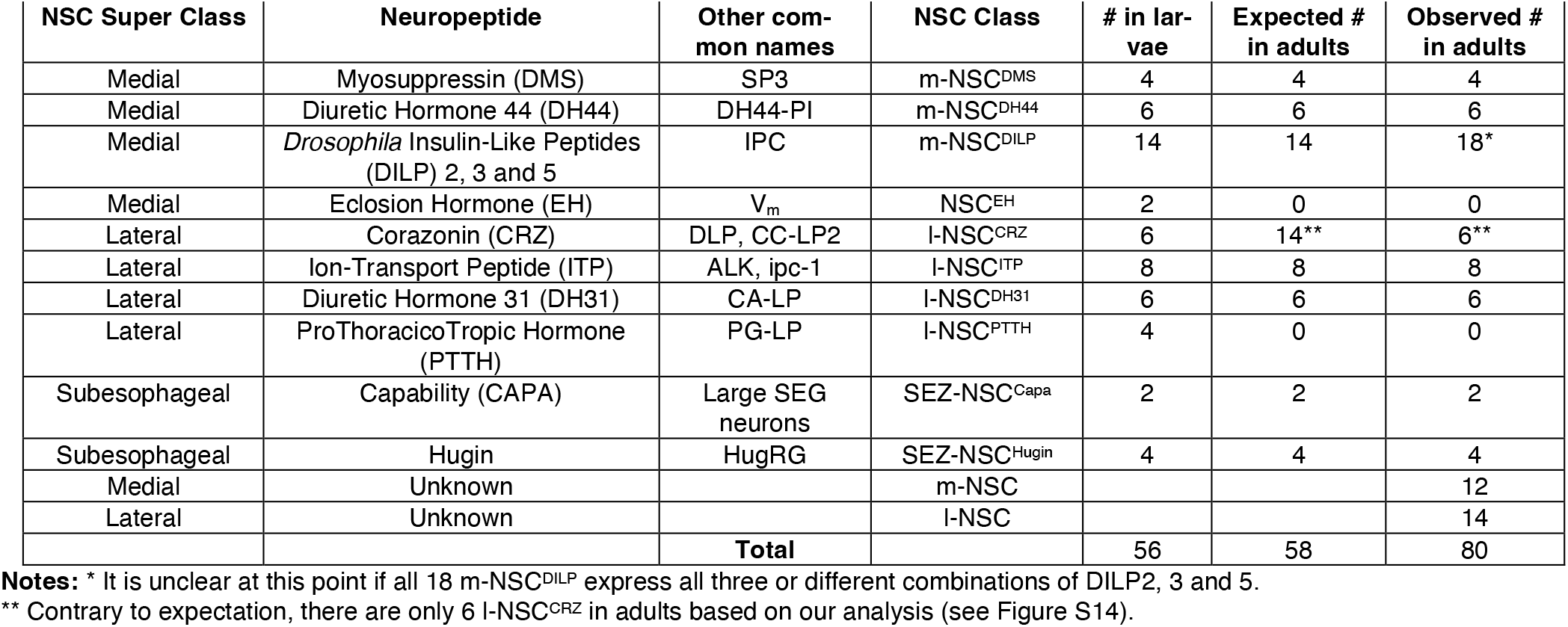
Classification of *Drosophila* neurosecretory cells (NSC) based on their cell body position in the central brain.

**Figure 5:**
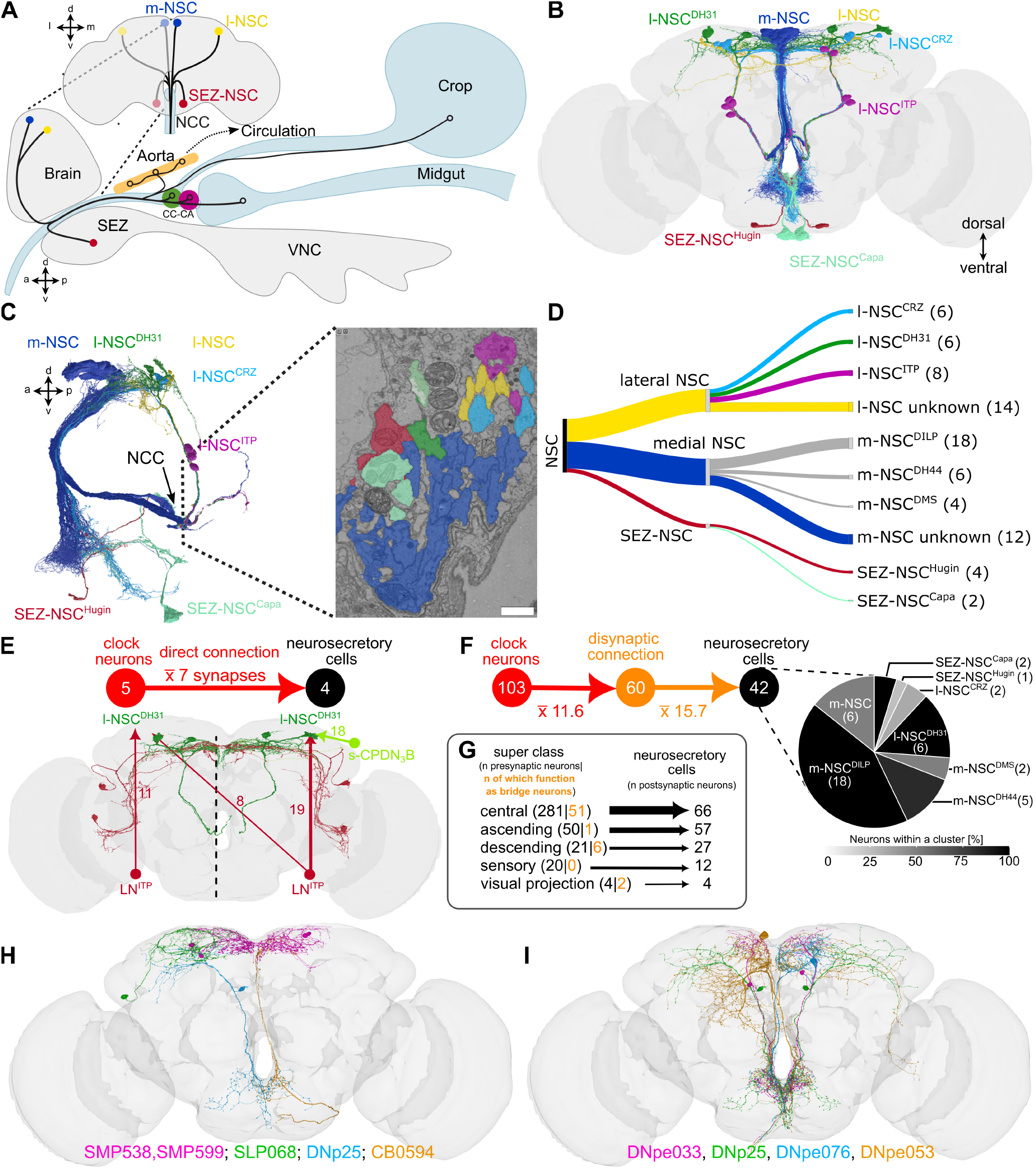
Identification of *Drosophila* neurosecretory cells and their clock inputs. **(A)** Schematic drawing of the different types of neurosecretory cells (NSC) and their projections to different release sites within the fly. **(B)** Reconstructions of the 80 NSC within the connectome. **(C)** All NSC projections exit the brain via NCC. Electron micrograph showcasing a cross section of the NCC. Scale bar = 750nm **(D)** Identification of 80 NSC and their classification based on location and neuropeptide expression. Refer to Table 2 for further details. **(E)** Direct inputs from clock neurons to NSC. LN^ITP^ (both hemispheres) and one s-CPDN_3_B (right hemisphere only) are the only clock neurons directly connected to NSC (l-NSC^DH31^). Numbers above the arrows represent the average number of synapses per connection. **F)** Disynaptic connections increase the connectivity between clock neurons and NSC. Numbers in circles indicate the number of neurons. The color gradient represents the percentage of neurons forming the connections relative to all cells of that corresponding group. **(G)** Total inputs to NSC grouped by the neuronal super classes annotated in the FlyWire connectome. Orange numbers indicate the subsets which mediate disynaptic connections between clock neurons and NSC. **(H)** Reconstructions of non-clock neurons that mediate strong disynaptic connections (>49 synapses with NSC) between the clock network and NSC. **(I)** Descending neurons that mediate disynaptic connections between clock neurons and NSC. Abbreviations: CC, corpora cardiaca; CA, corpora allata; NCC, nervii corpora cardiaca; VNC, ventral nerve cord; SEZ, subesophageal zone.

While all adult SEZ-NSC and some l-NSC can easily be classified based on their morphology and location (Figure 5B-C), this approach is not feasible for m-NSC since they are clustered together in the superior medial protocerebrum and appear similar based on gross morphology. Therefore, we asked whether cosine similarity-based clustering, such as the one used previously for clock neurons, can be used to distinguish and identify different m-NSC clusters, as well as the l-NSC^CRZ^ cluster. As expected, SEZ-NSC^Hugin^, SEZ-NSC^Capa^, and l-NSC^DH31^ form three separate clusters (Figure S13A-C). Most l-NSC^ITP^ (Figure S13B) do not have any input synapses in our dataset and were thus excluded from this analysis. Notably, this analysis resulted in two clusters of m-NSC comprising 4 and 6 neurons each. Hence, these clusters likely represent m-NSC^DMS^ and m-NSC^DH44^, respectively (Figure S13A and D). We obtained two additional clusters of m-NSC comprising 18 and 12 neurons, with the latter having low similarity between the neurons. The cluster comprising 18 m-NSC represents IPCs (m-NSC^DILP^). Whether all cells in this cluster express DILP2,3 and 5 remains unknown; however, DILP2 is expressed in more than 14 neurons in adults (Ohhara *et al*., 2018).

Interestingly, we could only reliably identify 6 out of the expected 14 l-NSC^CRZ^ (Figure S13B and Table 2). These 6 neurons cluster into two separate clades (Figure S13A) as they represent a heterogenous population both anatomically and functionally (Oh *et al*., 2019, Zandawala *et al*., 2021). Our inability to identify the remaining 8 CRZ neurons inspired us to examine if these adult-specific CRZ neurons are indeed neurosecretory. Using *Gr64a-Gal4* to label the adult-specific CRZ neurons (Fujii *et al*., 2015), we showed that there are only 6 adult l-NSC^CRZ^ (Figure S14). Contrary to our expectation, the adult-specific CRZ neurons do not project via the NCC and are thus not endocrine. Hence, our clustering analysis accounts for all the NSC that persist into adulthood. Additionally, it uncovered 12 putative m-NSC and 14 putative l-NSC in the adult brain (Figure 5D, S13E, Supplementary Video 5). These neurons have smaller somata compared to other identified NSC (Figure S13) and have relatively fewer dense core vesicles than neurons such as l-NSC^ITP^ (data not shown). Hence, the type (neuropeptide, biogenic amine, or fast-acting neurotransmitter) and the identity of the signaling molecules within these neurons remain unknown.

Informed by these new insights on different subgroups of NSC, we first explored potential inputs from clock neurons. Intriguingly, we observed sparse direct inputs from clock neurons to most NSC (Figure 5E), despite clock neuron projections being closely associated with NSC dendrites in the superior medial and lateral protocerebrum. The only exceptions are l-NSC^DH31^ which receive inputs from LN^ITP^ and s-CPDN_3_B. (Figure 5F). This observation prompted us to examine other types of synaptic inputs to NSC to ensure that the lack of synaptic connectivity between the clock network and NSC was not due to false negatives. Consistent with the location of m-NSC dendrites in the tritocerebrum (King *et al*., 2017, Zandawala *et al*., 2018), a large portion of their inputs derive from central, ascending and sensory neurons (Figure 5G). Therefore, a lack of direct clock output to the NSC is genuine, and this connectivity is likely indirect or paracrine in nature. To explore the extent of indirect connections, we examined disynaptic connectivity between clock neurons and NSC. Approximately 43% of the clock neurons provide inputs to half of the NSC disynaptically. The interneurons which facilitate these connections mainly include central neurons (Figure 5G and H) and descending neurons (Figure 5I). Taken together, these prominent indirect connections between the clock network and NSC could form the basis of circadian regulation of systemic physiology.

### Identifying the molecular basis of paracrine clock output pathways

Given the large repertoire of neuropeptides previously shown to be expressed in clock neurons (Abruzzi *et al*., 2017, Ma *et al*., 2021, Ma *et al*., 2023), we predicted that peptidergic paracrine signaling is crucial in mediating intercellular coupling within the clock network as well as output to downstream neurons such as NSC. First, we determined if any additional neuropeptides are expressed in clock neurons. To address this, we used a publicly available single-cell transcriptome dataset of clock neurons (Ma *et al*., 2021) combined with immunohistochemical localization and T2A-Gal4 lines. Unsupervised clustering of all clock neuron transcriptomes using t-SNE analysis yields 32 independent clusters (data not shown), 16 of which have high expression of clock genes (*tim* and *Clk*) and can be reliably identified based on known markers (Figure 6A-B). Our analysis revealed that most clock clusters express at least one neuropeptide (Figure 6B). Consistent with previous studies, l-LN_v_ express high levels of *Pdf*, whereas s-LN_v_ express both *Pdf* and *short neuropeptide F* (*sNPF*) (Figure 6B). Similar coexpression of neuropeptides is also observed in other clusters including the “DN_1p_^CNMa & AstC^” cluster which coexpresses *CNMamide* (*CNMa*), *AstC*, and *Dh31* neuropeptides. In total, at least 12 neuropeptides are highly expressed in the clock network (Figure 6B-C). Importantly, this includes novel clock-related neuropeptides, namely DH44 and Proctolin (Proc). *Dh44* is expressed in several clock clusters including DN_1a_, DN_1p_^AstA^, DN_1p_^sNPF^, DN_3_^VGlut^, LPN and LN_d_^NPF^ (Figure 6B-C). We independently confirmed the presence of DH44 peptide in these clusters using a combination of DH44 antibody or *DH44-T2A-Gal4* (Figure 6D, S15, and S16). *Proc*, on the other hand, is strongly expressed in the DN_1p_^CNMa^ and weakly in DN_2_ clusters which was verified by driving GFP using a *Proc-T2A-LexA* driver (Figure 6E and S15). *Proc* expression in other clock clusters such as LN_d_^NPF^ and DN_3_^VGlut^, remains to be validated. Lastly, we also detected *AstC* expression in additional clock neurons. AstC immunoreactivity was previously localized in DN_1p_, DN_3_, and LPN (Diaz *et al*., 2019, Reinhard *et al*., 2022a), which is in agreement with *AstC* transcript expression in DN_1p_^Rh7^ and DN_1p_^CNMa^ & ^AstC^ clusters (Figure 6B-C). Here, we show that AstC is additionally expressed in DN_2_, which were labeled using an antibody against VRI (Figure 6B, 6F, S15, and S16). Our expression analyses revealed the comprehensive neuropeptide complement of clock neurons (Figure 6B-F, S15, and S16) and provides the basis to explore paracrine targets of clock neurons.

**Figure 6:**
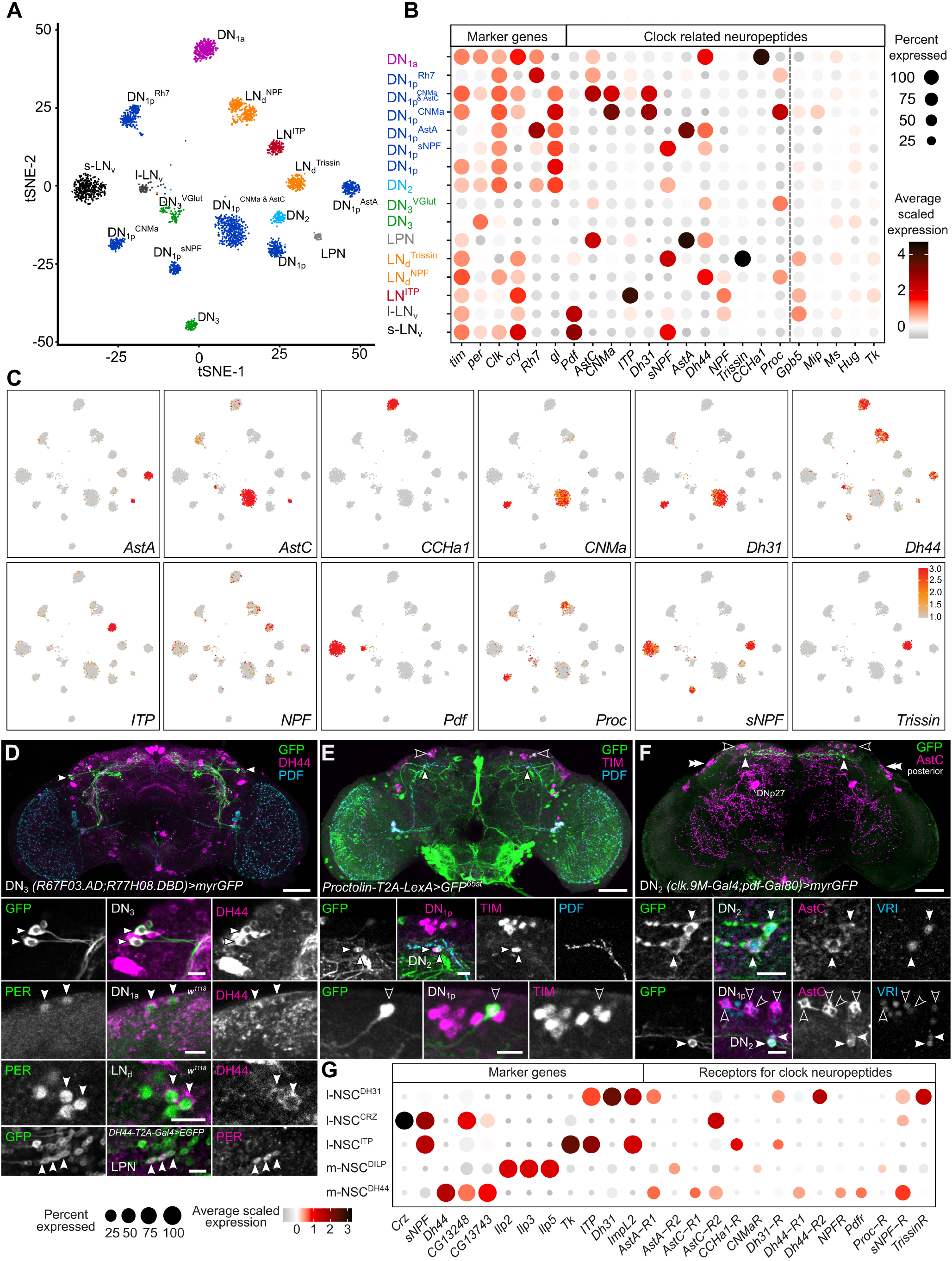
Circadian clock neuropeptidome. **(A)** Single-cell RNA sequencing of clock neurons reveals 16 distinct clock neuron clusters (shown in a t-SNE plot) that can be reliably identified based on known markers. **(B and C)** Clock neurons express at least 12 different neuropeptides. Based on average scaled expression (red = high and grey = low). **(D)** DH44 is a new clock-related neuropeptide which is expressed in APDN_3_, DN_1a_, LN_d_ and LPN (arrow-heads). **(E)** Proctolin is a new clock-related neuropeptide which is expressed in one DN_1p_ (open arrowhead) and two DN_2_ (filled arrowheads). **(F)** AstC is expressed in DN_2_ in addition to other clock neurons. Double arrowheads indicate DN_3_ labelled by anti-AstC, filled arrowhead indicate DN_2_, and open arrowheads indicate DN_1p_. Scale bars = 50μm for overview and 10μm for higher magnification images. Abbreviations: PER, Period; TIM, Timeless; VRI, Vrille; PDF, Pigment dispersing factor; DH44, Diuretic hormone 44; AstC, Allatostatin-C. **(G)** Single-cell transcriptomes of NSC express receptors for clock-related neuropeptides. Based on average scaled expression (red = high and grey = low).

As a first step in this direction and to validate our approach, we focused on select NSC which have been extensively characterized previously. We predicted that NSC are targeted by clock-related neuropeptides since they receive sparse monosynaptic inputs from clock neurons despite being closely associated with them anatomically. To investigate potential paracrine signaling between clock neurons and NSC, we again turned our attention to single-cell transcriptomics. We identified single-cell RNA transcriptomes of m-NSC^DH44^, m-NSC^DILP^, l-NSC^CRZ^, l-NSC^DH31^, and l-NSC^ITP^ (Davie *et al*., 2018) based on previously identified markers, and quantified the expression of clock peptide receptors (Figure 6G). We could not reliably mine m-NSC^DMS^ due to the lack of multiple molecular markers. We also disregarded SEZ-NSC^Capa^ and SEZ-NSC^Hugin^ from this analysis since they are much further away from clock neuron projections. Consistent with our prediction, our analysis indicated that multiple receptors for clock peptides are indeed expressed in NSC. Modulatory inputs to m-NSC^DILP^ have been examined extensively and our analysis is in agreement with previous expression and functional studies (Kapan *et al*., 2012, Nässel and Vanden Broeck, 2016, Nagy *et al*., 2019, Oh *et al*., 2019, Held *et al*., 2023). Taken together, our analysis uncovers the molecular substrates of paracrine signaling between clock neurons and NSC.

We next explored the magnitude of peptidergic paracrine signaling between clock neurons themselves. To predict putative paracrine connections, we extensively mapped the expression of clock neuropeptide receptors within the clock network. Consistent with promoter and genomic fragment-based *Pdfr-Gal4* expression (Im and Taghert, 2010), *Pdf receptor* (*Pdfr*) transcript is highly enriched in most clock clusters (Figure 7A-B), which we verified independently by expressing GFP using *Pdfr[RA]-T2A-Gal4* (Figure 7C and D). Other receptors are more sparsely expressed within the clock network (Figure 7A-B). However, most clock clusters express at least two receptors, with DN_1p_^sNPF^ expressing at least 7 receptors. We validated our single-cell transcriptome analysis by mapping the expression of select receptors using T2A-Gal4 knock-in lines. Our anatomical mapping of receptors (Figure 7D and S17) is largely in agreement with transcriptome data. In some cases, however, receptor mapping can provide additional insights. For instance, there are four transcript variants (A, B, C, and D) encoding the *Drosophila* Neuropeptide F receptor (NPFR). Using Gal4 lines specific for NPFR-A/C and NPFR-B/D isoforms, we showed that these isoforms are differentially expressed across the clock clusters, with A/C isoforms expressed more broadly than B/D isoforms (Figure 7D and S17).

**Figure 7:**
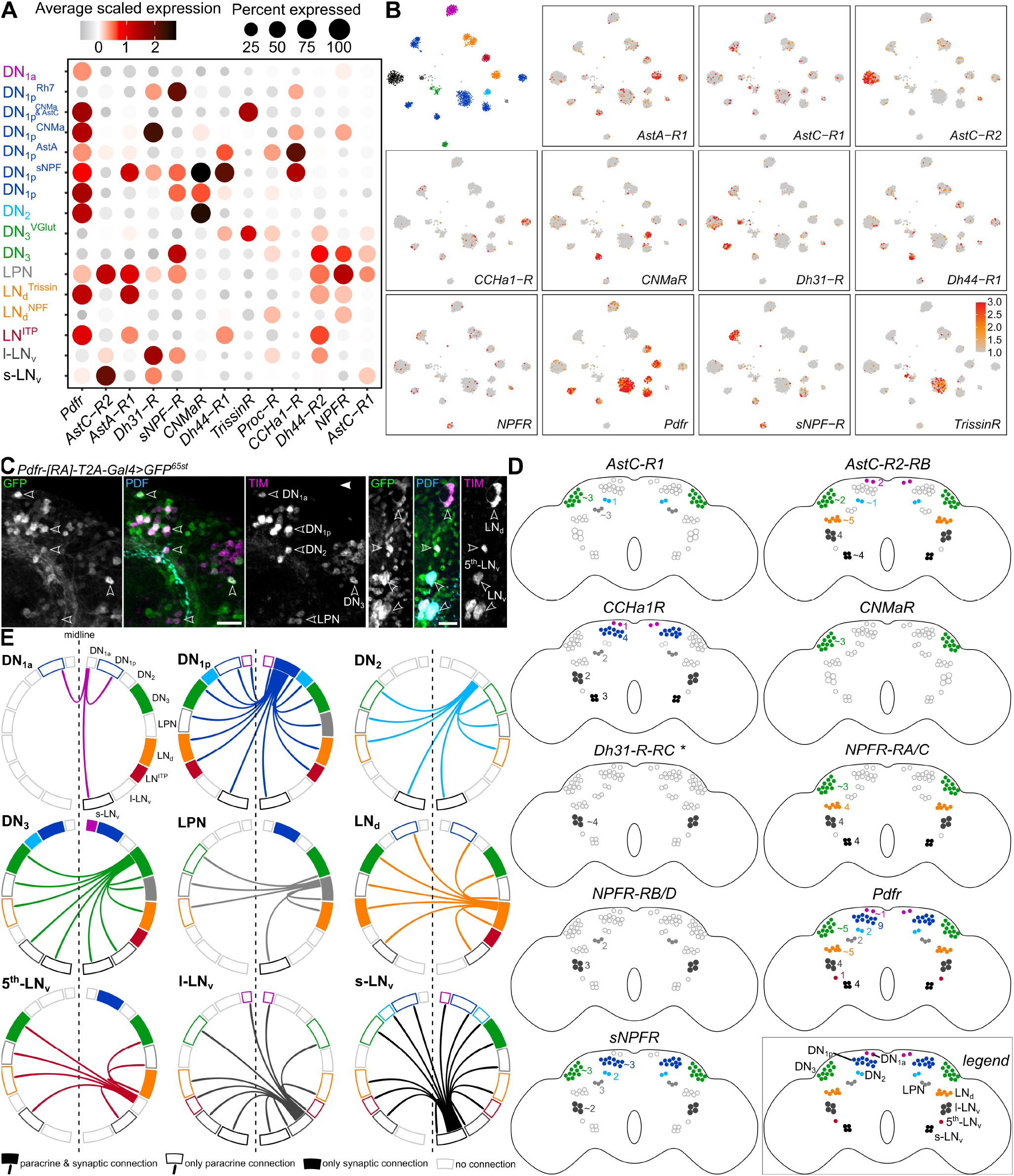
Paracrine signaling within the clock network. **(A)** Expression of clock-related neuropeptide receptors in different clock clusters identified using single-cell transcriptome analysis. **(B)** t-SNE plots showing the expression of receptors in different clock clusters. Clustering is based on Figure 6A. **(C)** *Pdfr[RA]-T2A-Gal4* drives GFP expression in all clock cell types. Arrowheads indicate different types of clock neurons that express PDFR[RA]. Scale bars = 20μm. Abbreviations: PDF, Pigment dispersing factor; TIM, Timeless. **(D)** Schematics depicting expression of receptors in different clock neurons. The schematics are based on GFP expression using T2A-Gal4 lines for different receptors (see Supplementary Figure 17 for confocal images). The numbers refer to neurons expressing that receptor in one hemisphere. **(E)** Chord diagrams showing synaptic (filled boxes) and putative peptidergic paracrine connections (arrows) between clock neurons. This figure is based on synaptic connectivity in Figure 1E, and peptide and receptor expression (following thresholding) mapping reported in Figures 6B-F, 7A-D and S15-17. See Figure S18 for a detailed explanation on filtering of putative paracrine connections.

Finally, we utilized our expression data of neuropeptides and their cognate receptors in clock neurons to delineate putative paracrine signaling pathways within the network. For this, we utilized an approach (Figure S18) similar to the one used recently to predict the *Caenorhabditis elegans* neuropeptide connectome (Ripoll-Sanchez *et al*., 2023). Briefly, we used expression data based on independent methods to conservatively localize the expression of neuropeptides and their receptors across all the clock clusters. We also utilized the connectome to factor in the distance between neurons of different clusters. This was done to ensure that the cells releasing the peptide and those expressing its receptor are not further apart than a cut-off of 14μm, which was set based on a previous paracrine connectivity study (Nagy *et al*., 2019). Lastly, we only considered strong peptide-receptor interactions by disregarding ligands with EC_50_ values for receptor activation higher than 500nM. This stringent *in silico* approach allowed us to predict paracrine connectivity within the clock network with high confidence. Taking s-LN_v_ as an example, this cluster expresses both PDF and sNPF. Following our expression thresholding, PDFR is expressed in most clock clusters, whereas sNPF receptor is expressed in DN_1p_, DN_3_, LPN and l-LN_v_ (Figure 7 and S18). Thus, in addition to providing synaptic inputs to DN_3_, s-LN_v_ can potentially provide paracrine inputs to most clock clusters across both hemispheres (Figure 7E). Connectivity from l-LN_v_ is also enhanced by paracrine signaling, although not to the same extent as it is for s-LN_v_. Expanding this analysis to other clock clusters allowed us to comprehensively identify putative peptidergic signaling pathways between clock clusters (Figure 7E). These pathways, however, can only be considered putative for three main reasons: (1) all clock neuropeptides are also expressed in other non-clock neurons (Nässel and Zandawala, 2019), suggesting likely inputs from neurons extrinsic to the clock network, (2) we don’t account for peptide efficacies and receptor affinities since these values were independently determined in distinct systems thus making comparisons difficult and (3) despite our best efforts to account for it, distance of peptide diffusion may vary. Nonetheless, the presence of receptors for clock peptides in other clock neurons provides the molecular basis for potential paracrine signaling between them. In summary, peptidergic signaling greatly enriches the connectivity between different subsets of clock neurons. Additional investigations are necessary to determine which of these putative connections are functional *in vivo*.

## Discussion

### Peptidergic signaling supplements synaptic connectivity within the circadian clock network

Using a multipronged approach centered around the FlyWire connectome (Dorkenwald *et al*., 2023, Schlegel *et al*., 2023), we describe the first whole-brain neural connectome of an animal circadian clock. Our analysis using the complete brain connectome eluded identification of one DN_2,_ and a couple of s-CPDN_3_. Nevertheless, given the fact that we identified almost all of the expected clock neurons as well as several additional DN_3_ (Table 1), our clock neuron synaptic wiring diagram is complete enough to be regarded as a connectome. Our clock connectome is also a significant upgrade (10-fold larger numerically) compared to the partial connectivity diagram based on the hemibrain connectome reported earlier (Shafer *et al*., 2022). The previous analysis was based on only 24 clock neurons and largely focused on LN clusters, while excluding l-LN_v_, and several DN_1p_, DN_2_, and DN_3_ clusters due to the incomplete nature of the dataset. However, as evident from our analysis here, the DN in fact represent an important hub in the clock network and display high synaptic connectivity. In particular, DN_1p_ play a large role in clock cluster interconnectivity and DN_3_ appear important for clock output pathways. Our analysis also sheds light on the precise number of DN_3_ in the clock network. Although approximately 80 DN_3_ were previously estimated in the entire clock network, there are in total about 170 DN_3_ based on our connectome and anatomical analyses. We anticipate that resources such as NeuronBridge (Meissner *et al*., 2023), which allow for comparisons between electron and light microscopy datasets, will facilitate the identification of Gal4 drivers that target the novel s-CPDN_3_ subtypes identified here. In addition, we also characterized the molecular basis for neuropeptide connectivity between the clock neurons, consequently highlighting putative peptidergic pathways within the clock network. Similar to vertebrates (Wen *et al*., 2020, Morris *et al*., 2021), the *Drosophila* clock network is highly peptidergic, with all clock neuron clusters expressing at least one neuropeptide. Notably, a majority of the clock clusters express two neuropeptides, and several express three, while the LPN express four neuropeptides. Similar neuropeptide coexpression is also evident in SCN neurons and thus appears to be a common feature of clock neurons (Wen *et al*., 2020). Surprisingly, there is little to no overlap in the neuropeptide complement of the *Drosophila* and vertebrate clock neurons (Figure 8). Orthologs of vertebrate clock neuropeptides including vasoactive intestinal peptide, arginine vasopressin, neuromedin S, cholecystokinin, gastrin-releasing peptide, and prokineticin 2 are either absent in the *Drosophila* genome or expressed outside the clock network. Hence, *Drosophila* and vertebrates have evolved to utilize different signaling molecules while still conserving the diversity of neuropeptide signaling within the clock networks. Remarkably, except for PDF, there appears to be little conservation in neuropeptide identities of clock neurons across different insects (Stengl and Arendt, 2016, Beer and Helfrich-Förster, 2020). This suggests that it is more important to conserve the mode of communication (paracrine signaling) rather than the messenger (specific neuropeptide).

**Figure 8:**
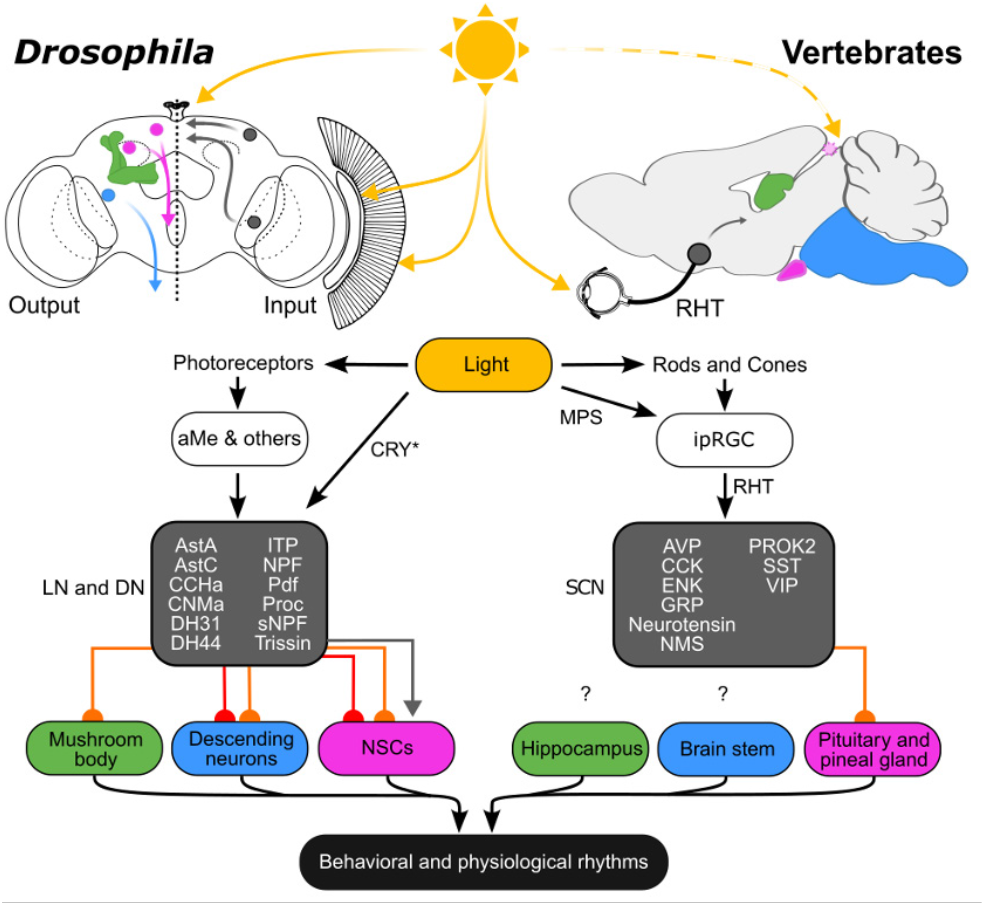
Parallels in *Drosophila* and vertebrate clock input and output pathways. Direct and indirect light input pathways to the *Drosophila* clock (comprised of LN and DN) and vertebrate suprachiasmatic nucleus (SCN). Note that the pineal gland receives light input only in non-mammalian vertebrates. *Drosophila* and vertebrate clocks utilize different neuropeptides (grey box). Output from the clock to downstream targets is either synaptic (direct in red and indirect in orange) or paracrine (grey arrow). Abbreviations: CRY, Cryptochrome; MPS, Melanopsin, RHT, Retinohypothalamic tract; ipRGC, Intrinsically-photosensitive retinal ganglion cells; aMe, Accessory medulla neurons; NSC, neurosecretory cell; AVP, arginine vasopressin; CCK, cholecystokinin; ENK, met-enkephalin; GRP, gastrin-releasing peptide; NMS, neuromedin S; PROK2, prokineticin 2; SST, somatostatin; VIP, vasoactive intestinal peptide.

### Contralateral connectivity within the network prevents decoupling of clock neurons across the hemispheres

Analysis of interconnectivity within the clock network revealed extensive contralateral synaptic connectivity between the clock neurons, which is largely mediated by DN_1p_A and to a lesser extent by s-CPDN_3_C, s-CPDN_3_D, l-CPDN_3_, and LN^ITP^. Furthermore, paracrine peptidergic signaling amongst clock neurons has the potential to further strengthen this contralateral connectivity. Such a strong bilateral coupling of clock neurons prevents the internal desynchronization of clock neuron oscillations between the two hemispheres – a phenomenon that can happen in other insects and even in mammals, but so far has not been observed in fruit flies (Helfrich-Förster, 2004). In addition, consistent with previous analysis using the hemibrain connectome (Shafer *et al*., 2022), we did not observe any synaptic connectivity between the s-LN_v_ and the LN_d_ neurons, which control morning and evening activities, respectively. In line with this observation, the phase relationship between morning and evening oscillators is plastic, consequently facilitating seasonal adaptations as in mammals (Yoshii *et al*., 2012). Our data may also explain how the morning and evening oscillators in flies internally desynchronize under certain conditions (Helfrich-Förster, 2014). One such relevant condition is increased PDF signaling during long days (Hidalgo *et al*., 2023), which was shown to delay the evening oscillators (Liang *et al*., 2016, Liang *et al*., 2017, Vaze and Helfrich-Förster, 2021) and may lead to internal desynchronization (Wulbeck *et al*., 2008). Here, we confirm the presence of PDFR in the evening neurons. Thus, enhanced PDF signaling could delay the evening neurons and bring them out of phase with the morning neurons, especially because the two sets of neurons are not connected via synapses. Taken together, the lack of interconnectivity between s-LN_v_ and LN_d_ could potentially be a factor promoting adaptation to different seasons and contexts.

### Light and other inputs to Drosophila and vertebrate circadian clocks

Our analyses reveal that extrinsic light input from the photoreceptor cells of the compound eyes, HB eyelets, and ocelli to the clock neurons is largely indirect, with the former two transmitting light inputs via aMe neurons. This situation may appear to be different from mammals where intrinsically photosensitive retinal ganglion cells in the retina project directly through the retinohypothalamic tract onto the SCN neurons (Figure 8) (Starnes and Jones, 2023). However, even the mammalian clock receives indirect photoreceptor inputs from rods and cones via bipolar cells and retinal ganglion cells. Furthermore, indirect light inputs may reach the SCN also via the intergeniculate leaflets of the thalamus (Kim and Harrington, 2008). Altogether, this suggests that the fly and mammalian system are not fundamentally different. One apparent difference is the presence of CRY, a cell-autonomous circadian photoreceptor, in subsets of *Drosophila* clock neurons that is sufficient for light entrainment in eyeless mutants (Helfrich-Förster *et al*., 2001). Mammals lack light-sensitive CRY. Instead, they possess light-sensitive melanopsin in retinal ganglion cells, and mice lacking rods and cones can still entrain to light/dark cycles due to melanopsin (Foster *et al*., 2020). Thus, flies and mice possess several redundant and partly parallel light-input pathways to entrain their clocks. The similarity is even higher when comparing flies with vertebrates in general. For instance, fish, birds, and reptiles possess additional photoreceptors in the pineal gland, which is reminiscent of the extraretinal HB eyelets or even the ocelli of flies (Figure 8). Most importantly, vertebrates and flies use their eyes for both vision and entraining their circadian clocks, tasks that require completely different properties of light inputs. Vision requires image formation and fast neurotransmission, whereas circadian entrainment is dependent on integrating light collection over a longer time that can be at a slower rate. The connectome reveals that the number of synapses as well as the axon thickness of the neurons mediating this connectivity are very different between the two types of photoreception. Hence, they are aptly suited to perform their required functions.

### Paracrine modulation of the neuroendocrine system by circadian clock

Since circadian control of organismal physiology is likely mediated via hormones, in parallel with the clock connectome, we also identified the cells that make up the neurosecretory center of an adult *Drosophila* brain. This neurosecretory center is comprised of 80 endocrine cells, located in distinct regions of the brain and having unique neuropeptide identities. Neurosecretory connectomes of larval *Drosophila* and the marine annelid, *Platynereis dumerilii* have been established previously (Williams *et al*., 2017, Huckesfeld *et al*., 2021). Unfortunately, these studies did not examine the connectivity between the circadian clock and the neuroendocrine centers. Thus, it remains to be seen whether the largely indirect and paracrine signaling between the adult *Drosophila* circadian network and NSC is a phenomenon conserved across other animals. However, given the relatively slow timescales at which the circadian output needs to be propagated to downstream neurons, neuropeptides seem suited for this role.

### Limitations of our approach

The novel synaptic and putative paracrine connections reported here can only be considered predictions until they are functionally verified. In the case of paracrine connectivity, functional connectivity experiments (in normal and peptide mutant flies) using electrophysiological methods or genetically encoded secondary messenger sensors (calcium and cAMP) are needed to confidently establish functional paracrine connectivity. Additionally, the synaptic connectivity reported here is likely an underestimation due to several factors: 1) we did not explore connectivity via gap junctions, 2) approximately 30% of synapses are missing for photoreceptors (Dorkenwald *et al*., 2023) and 3) we generally used a connectivity threshold of >4 synapses. Preliminary expression analysis using the single-cell transcriptomes of clock neurons suggests that gap junction genes are enriched in the clock network (data not shown) and they can influence activity-rest rhythms (Ramakrishnan and Sheeba, 2021). It remains to be seen which clock neurons are additionally coupled via gap junctions and how this electrical connectivity complements synaptic and peptidergic connectivity detailed here. Moreover, as discussed earlier, fewer than 5 synapses could also represent functional connections which were largely disregarded in our analyses. While the FlyWire and hemibrain connectomes exhibit a high degree of stereotypy, they are both based on an adult female brain. The lack of a male brain connectome currently prevents any comparisons on sex-specific differences within the circadian network and its output pathways which could influence sexually dimorphic behaviors and physiology. Further, the connectome provides a singular snapshot of connectivity which could change depending on the time of day, the age of the animal as well as its internal state.

## Conclusion

In conclusion, our circadian clock connectome, the first of its magnitude, is a significant milestone in chronobiology. Given the high conservation of circadian network motifs between *Drosophila* and vertebrates, this connectome provides the framework to systematically investigate circadian dysregulation which is linked to various health issues in humans including sleep, metabolic, and mood disorders. Moreover, it will also facilitate the development and experimental validation of novel hypotheses on clock function.

## Acknowledgments

We would like to thank Selina Hilpert and Barbara Mühlbauer for technical assistance, Maria Steigmeier for preliminary investigations on PDFR expression, and Tatsuya Yokosako for generating the LN^ITP^ split-Gal4 line. We are also grateful to Drs. Fumika Hamada for *Clk*.*9M-Gal4;pdf-Gal80*, Shu Kondo for T2A-Gal4 lines, Paul E. Hardin for VRI antibody, Ralf Stanewsky for PER antibody, Heinrich Dircksen for ITP antibody, Susan Morton for RFP antibody, and Jan Veenstra for AstC, CRZ and DH44 antibodies. We thank the Princeton FlyWire team and members of the Murthy and Seung labs, as well as members of the Allen Institute for Brain Science, for the development and maintenance of FlyWire (supported by BRAIN Initiative grants MH117815 and NS126935 to Murthy and Seung). We also acknowledge members of the Princeton FlyWire team and the FlyWire consortium, especially Drs. Gregory Jefferis, Sven Dorkenwald, and Philipp Schlegel for troubleshooting and guidance. Special thanks to Austin T Burke and Mareike Selcho from the FlyWire consortium for contributing >10% effort towards editing and proofreading at least 10% of clock neurons. We are also thankful to Drs. Theresa McKim and Dick Nässel for helpful feedback during the preparation of this manuscript, as well as the Division of Instrumental Analysis, Okayama University for the laser scanning confocal microscopes (FV1200 and FV3000). M.Z. was supported by funding from the University of Würzburg, Deutsche Forschungsgemeinschaft (DFG; ZA1296/1-1), and NV INBRE. D.R. (DFG; RI 2411/1-1) and C.H.F. (DFG; FO 207/16-1) were supported by DFG. T.Y. was supported by JSPS (KAKENHI 19H03265). A.S. and M.S. were supported by the OU fellowship (JST SPRING, Grant Number JPMJSP2126). We also acknowledge funding from the DFG for the Leica TCS SP8 microscope (251610680, INST 93/809-1 FUGG). This manuscript was typeset with the bioRxiv word template by @Chrelli: www.github.com/chrelli/bioRxiv-word-template.

## Author contributions

N.R., C.H.F., T.Y., and M.Z. conceived the study. N.R., D.R., C.H.F., T.Y., and M.Z. supervised the project. N.R., A.F., G.M., E.D., A.S., G.M., M.S., and M.Z. performed the experimental work and analyzed the data. N.R. and M.Z. performed computational analyses. M.Z. wrote the manuscript with input from C.H.F and N.R. All authors read, provided feedback, and approved the final manuscript.

## Materials and Methods

### Fly strains

*Drosophila melanogaster* strains used in this study are listed in Supplementary Table 1. T2A-Gal4 and T2A-LexA knock-in lines generated previously (Deng *et al*., 2019, Kondo *et al*., 2020) were obtained from the Bloomington *Drosophila* Stock Center (BDSC) and Dr. Shu Kondo. Flies were maintained under LD12:12. Flies for peptide and receptor mapping were reared at 25°C on *Drosophila* medium containing 0.7% agar, 8.0% glucose, 3.3% yeast, 4.0% cornmeal, 2.5% wheat embryo, and 0.25% propionic acid. Flies for *trans-* Tango and Gal4 verification were raised at 18°C and 25°C, respectively, on standard medium containing 8.0% malt extract, 8.0% corn flour, 2.2% sugar beet molasses, 1.8% yeast, 1.0% soy flour, 0.8% agar and 0.3% hydroxybenzoic acid.

### Immunohistochemistry and confocal imaging

#### Neuropeptide and receptor mapping

Immunostainings were performed as described previously (Yoshii *et al*., 2015). Briefly, male flies were entrained in LD12:12 at 25 °C for at least 3 days. Whole flies sampled at Zeitgeber time (ZT) 20 were fixed in 4% paraformaldehyde in phosphate-buffered saline (PBS) with 0.1% Triton X-100 (PBS-T) for 2.5 h at room temperature (RT). Fixed flies were washed three times with PBS, before dissecting their brains. Samples were washed with PBS-T three times. The samples were blocked in PBS-T containing 5% normal donkey serum for 1 hour at RT and subsequently incubated in primary antibodies at 4°C for 48 h. Following six washes with PBS-T, the brains were incubated in secondary antibodies at RT for 3 h. Lastly, the samples were washed six times in PBS-T and mounted in Vectashield mounting medium (Vector Laboratories, Burlingame, CA, USA). At least five brains for each strain were used for immunostaining to characterize the clock cells stained by anti-GFP antibodies (Gal4 or LexA expression) in the first experiment and the clock cells were briefly characterized with PDP1 and PDF antibodies. In the second experiment, we conducted the same immunostaining with anti-TIM antibodies, but only for positive strains, to confirm the prior results. Images were taken from at least three different brains using laser scanning confocal microscopes (Olympus FV1200 and FV3000, Olympus, Tokyo, Japan).

#### Mapping DH44 and AstC expression in clock neurons, verification of clock neuron Gal4 lines, and trans-Tango analysis

Flies for PER staining were fixed at ZT23 (1 hour before lights-on) since PER levels are maximal at this ZT (Zerr *et al*., 1990). Flies for VRI staining were fixed at ZT 20. Immunohistochemistry was performed as described previously (Reinhard *et al*., 2022a, Reinhard *et al*., 2022b). Images were scanned with a Leica TCS SP8 confocal microscope equipped with a photon multiplier tube and hybrid detector. A white light laser (Leica Microsystems, Wetzlar, Germany) was used for excitation. We used a 20-fold glycerol immersion objective (0.73 NA, HC PL APO, Leica Microsystems, Wetzlar Germany) for wholemount scans and obtained confocal stacks with 2048 × 1024 pixels with a maximal voxel size of 0.3 × 0.3 × 2 μm and an optical section thickness of 3.12 μm. For noise reduction, we used a frame average of 3. Images were analyzed using Fiji. All the primary and secondary antibodies are listed in Supplementary Table 2.

#### Multi-color flip-out

Multi-color flip-out (MCFO) analysis was performed to identify the morphology of the 8 so far uncharacterized DN_1p_ (Nern *et al*., 2015). *Clk4*.*1M-Gal4* flies were crossed to MCFO7 flies. The F1 generation was kept at 25°C and 1-7 days old male and female flies were used for staining. Immunohistochemistry was performed as described above. All the primary and secondary antibodies are listed in Supplementary Table 2.

#### DN_3_ quantification

To quantify the number of DN_3_ in the adult brain, *tim-(UAS)-Gal4>JFRC81-GFP* male and female flies were entrained at 25°C for 3 days and then fixed at ZT1. Brains were stained for GFP, PER, and VRI as described above. Images were acquired using a Leica TCS SP8 confocal microscope as described above. A 63-fold glycerol immersion objective (1.3 NA, HC PL APO, Leica Microsystems, Wetzlar Germany) was used to acquire detailed images of the DN_3_ cluster. Images were recorded at 1024 × 1024 pixels with a maximum voxel size of 0.18 × 0.18 × 0.33 μm and an optical section thickness of 0.99 μm. Cells were counted using the Multi-Point tool in Fiji. 5 brains each for males and females were used for quantification.

### Connectome datasets and neuron identification

For all analyses, we used the v783 snapshot of the FlyWire whole brain connectome and its annotations (Dorkenwald *et al*., 2023, Schlegel *et al*., 2023). We also used the adult hemibrain connectome (v1.2.1) for comparisons (Scheffer et al. 2020). To identify all NSC in the FlyWire connectome, we first identified NSC subsets that have a characteristic morphology and location e.g. l-NSC^ITP^. We then identified the nerve bundle (nervii corpora cardiaca, NCC) through which l-NSC^ITP^ axons exit the brain. This nerve bundle contains all the axons for NSC in the *Drosophila* brain. We manually assessed other axons in NCC to identify the remaining NSC in the connectome. Independently, we also examined the cross-section of the soma of all putative NSC for the presence of dense core vesicles, which suggests that they contain neuropeptides/neuromodulators. The pipeline for identifying clock neurons is summarized in Figure S1. Briefly, neurons were first identified based on morphological and/or connectivity features described previously (Lamaze *et al*., 2018, Schubert *et al*., 2018, Reinhard *et al*., 2022a, Reinhard *et al*., 2022b, Sun *et al*., 2022), or based on NBLAST similarity to identified neurons in the hemibrain (Schlegel *et al*., 2023). Morphology clustering (v630) for 124,988 FlyWire neurons (Schlegel *et al*., 2023) was then used to identify neurons that were morphologically similar to the clock neurons identified above. Several clock neuron types (e.g. APDN_3_, s-LN_v_, LN_d_, etc.) formed their own morphology cluster, suggesting that additional neurons similar to them do not exist in the dataset. If a morphology cluster comprised more neurons besides the clock neurons identified earlier, all the additional neurons were considered putative clock neurons (Figure S1), which were subsequently filtered based on different features. First, the cell body position was determined by mapping the coordinates of the nuclei (Mu *et al*., 2021) to the root IDs of the neurons. The average distance between the cell body position of neurons in one clock cluster was determined per hemisphere and candidate neurons laying less than twice the average distance away were retained for manual comparison of morphological features and connectivity described in previous studies. In addition, hemilineage and cell type information was used to determine whether the candidate neurons could be additional clock neurons. If new clock neurons were identified, the whole procedure was repeated. By definition, a cell type is a uniquely identifiable neuron in the dataset (Schlegel *et al*., 2023). Based on our analysis, if a certain cell type was part of a clock cluster, all other neurons of that cell type were also part of this clock cluster. Thus, the likelihood of other cells that have a similar morphology to the clock neurons identified here is extremely low. FlyWire cell IDs of identified clock neurons and NSC are provided in Supplementary Tables 3 and 4, respectively.

### Paracrine connectivity prediction

For paracrine connectivity analysis, peptide and receptor expression was first determined based on single-cell RNA sequencing analyses (Ma *et al*., 2021). A gene was considered to be expressed in a given cluster if 1) it was expressed in more than 49% of the cells within that cluster and 2) the expression level was above the set threshold. A threshold of 0.208 average scaled expression was used for peptides and 0.067 for receptors to align single-cell RNA sequencing data with previously established expression data. Further, T2A-Gal4 expression patterns for peptides and receptors were analyzed in conjunction with antibody staining. This information, together with published antibody staining and live imaging data for receptors (Supplementary Table 6), was used to generate expression matrices based on the different methods (Figure S18A). To reduce the number of false positives, we only considered the expression of peptides when shown by two independent methods. A receptor was deemed present if its expression was demonstrated using two independent methods or based on live imaging data (Figure S18A). Next, we calculated the distance between neurons of different clock clusters based on their skeletons (v783_L2). To calculate a threshold for the distance that a peptide can diffuse, we used the closest distance (14μm) between the s-LN_v_ and the NSC^DILP^ since paracrine PDF signaling between these neurons has been demonstrated previously (Nagy *et al*., 2019). If two clock clusters included neurons that were located closer than this distance, we assumed that peptidergic paracrine signaling was possible between these clusters (Figure S18B). Additionally, some *Drosophila* receptors are activated by multiple ligands. Hence, based on the previously reported EC_50_ values for receptor activation (Supplementary Table 6), we set a stringent threshold of 500nM to determine if a peptide can activate a receptor. Lastly, to identify putative paracrine connections, we combined the distance matrix with the peptide-receptor expression matrix (Figure S18C).

### Data visualization

Data was visualized using ggplot2 (v 3.4.2, Wickham, 2016) and circlize (v 0.4.15) for R (v 4.2.2) in RStudio (2022.12.0) (Gu *et al*., 2014). Reconstructions were downloaded using the navis library (v 1.0.4, https://github.com/navis-org) and cloud-volume library (v 8.10.0, https://github.com/seung-lab/cloud-volume) for python (v 3.8.5), and visualized using blender (v 3.01, Community, B. O. 2018). For visualizing a large number of neurons, Codex (doi: 10.13140/RG.2.2.35928.67844) and FlyWire neuroglancer were used (Dorkenwald *et al*., 2022).

### Connectivity analyses

Connectivity data was analyzed using the natverse libraries (v 0.2.4) for R in RStudio (Bates *et al*., 2020). Synaptic connectivity similarity for clock neurons was analyzed based on all their input and output synapses. In the case of NSC, connectivity similarity was based only on their input synapses as output synapses were largely absent. Cosine similarity analyses were conducted with coconatfly (v 0.1.1.9; https://github.com/flyconnectome/coconatfly) for R. Filtered synapses were retrieved in Python using the navis and pandas (v 1.1.3, (McKinney, 2010)) libraries. Unless stated otherwise, a connection with more than 4 synapses was considered significant.

### Single-cell transcriptome analyses

Expression of neuropeptides and their cognate receptors in clock neurons was mined using the single-cell RNA sequencing dataset and analysis pipeline established earlier (Ma *et al*., 2021). NSC transcriptomes were identified from the brain transcriptomes generated previously (Davie *et al*., 2018). The parameters used to identify the different cell types were based on previous studies and provided below (Kahsai *et al*., 2010, Kapan *et al*., 2012, Miyamoto and Amrein, 2014, Cannell *et al*., 2016, Yang *et al*., 2018, Nässel and Zandawala, 2019, Oh *et al*., 2019, Gera *et al*., 2024).

DH31 (6 cells): ITP > 2 & Dh31 > 4 & amon > 0 & Phm > 0

DH44 (12 cells): Dh44 > 1 & CG13248 > 1 & CG13743 > 0 & Lkr > 0

CRZ (4 cells): Crz > 3 & sNPF > 3 & Dh44 == 0 & ITP == 0 & ChAT == 0 & Gr64a == 0

ITP (7 cells): Tk > 1 & sNPF > 1 & ITP > 1 & ImpL2 > 1 & Crz = 0

IPC (19 cells): Ilp2 > 3 & Ilp3 > 3 & Ilp5 > 3

Expression was scaled based on all genes in the dataset. All analyses were performed in R-Studio (v2022.02.0) using the Seurat package (v4.1.1 (Hao *et al*., 2021)).

## Data Availability

Connectivity analysis can be performed using the cell IDs provided at https://codex.flywire.ai/.

## Code Availability

Code used to visualize the data will be made available upon request.

