## Supplementary File for "Synaptic connectome of the *Drosophila* circadian clock"

bioRxiv

### Supplementary material

**Supplementary Table 1:** Fly strains used in this study

| Fly strain | BDRC Stock number / Reference |
|---|---|
| *QUAS-nlsRFP; UAS-trans-Tango* | (Snell *et al.*, 2022) |
| *w; R43D05AD/Cyo; R93B11 DBD/TM6B (DN$_{1a}$)* | (Sekiguchi *et al.*, 2020) |
| *yw; + ; Clk4.1M-Gal4 (DN$_{1p}$)* | (Zhang *et al.*, 2010) |
| *w; R43D05AD/Cyo; VT003234DBD/TM6B (DN$_2$)* | (Sekiguchi *et al.*, 2020) |
| *w; R67F03AD/Cyo; R77H08DBD/TM6B (DN$_3$)* | 70786, 69630 (Meiselman *et al.*, 2022) |
| *w; R11B03AD/Cyo; R65D05DBD/TM6B (LPN)* | (Sekiguchi *et al.*, 2020) |
| *w; R22E04AD ; R18F07DBD (LN$^{ITP}$)* | 49309, 70047 (This study) |
| *yw; PDF-Gal4 (LN$_v^{PDF}$)* | (Park *et al.*, 2000) |
| *w$^{1118}$* | 5905 |
| *w; +; 10xUAS-myr::GFP (myrGFP)* | (Pfeiffer *et al.*, 2010) |
| *UAS-GFP S65T* | 1522 |
| *13xLexAop2-6xGFP* | 52266 |
| *JFRC81-10xUAS-IVS-Syn21-GFP-p10 (EGFP)* | (Pfeiffer *et al.*, 2012) |
| *MCFO7* | 64091 (Nern *et al.*, 2015) |
| *Clk.9M-Gal4; pdf-Gal80* | (Kaneko *et al.*, 2012) |
| *tim-(UAS)-Gal4* | 80941 (Blau and Young, 1999) |
| *Gr64a-Gal4* | 57661 |
| *CCHa1-LexA* | 84361 |
| *Proc-LexA* | 84432 |
| *AstA-Gal4* | 84593 |
| *AstC-Gal4* | 84595 |
| *CNMa-Gal4* | 84619 |
| *Dh31-RA/C-Gal4* | 84623 |
| *Dh44-Gal4* | 84627 |
| *sNPF-Gal4* | 84706 |
| *AstC-R1-Gal4* | 84596 |
| *CNMaR-Gal4* | 84620 |
| *Dh31R-RA/B/C-Gal4* | 84625 |
| *Dh31R-RC-Gal4* | 84626 |
| *NPFR-RA/C-Gal4* | 84672 |
| *NPFR-RB/D-Gal4* | 84673 |
| *PDFR-RA-Gal4* | 84684 |
| *PDFR-RA-Gal4* | (Kondo *et al.*, 2020) |
| *sNPFR-Gal4* | 84691 |
| *AstC-R2-RB-Gal4* | (Kondo *et al.*, 2020) |
| *CCHa1R-Gal4* | (Kondo *et al.*, 2020) |

**Supplementary Table 2:** Antibodies used for immunohistochemistry

| Antibody | Dilution | Source / Reference |
|---|---|---|
| chicken anti-GFP | 1:1000 | Rockland, Limerick, PA, USA |
| chicken anti-GFP | 1:1000 | Abcam, RRID: AB_300798 |
| guinea pig anti-RFP | 1:5000 | Gift from Dr. Susan Morton |
| mouse C7 anti-PDF | 1:1000 | Developmental Studies Hybridoma Bank; (Cyran *et al.*, 2005) |
| rabbit anti-ITP | 1:5000 | (Hermann-Luibl *et al.*, 2014) |
| guinea pig anti-ITP | 1:1000 | (Manoli *et al.*, 2023) |
| rabbit anti-DH44 | 1:1000 | (Cabrero *et al.*, 2002) |
| rabbit anti-PDP1 | 1:9000 | (Cyran *et al.*, 2003) |
| rat anti-TIM | 1:3000 | (Yoshii *et al.*, 2008) |
| mouse nc82 anti-Bruchpilot | 1:50 | (Wagh *et al.*, 2006) RRID: AB_2314866 |
| mouse M2 anti-FLAG-tag | 1:500 | Sigma-Aldrich #F1804 |
| human anti V5-tag [SV5-P-K] | 1:500 | Abcam #ab206562 |
| rabbit anti-HA-tag | 1:800 | Sigma-Aldrich #H6908 |
| guinea pig anti-VRI | 1:2000 | (Glossop *et al.*, 2003) |
| goat dN-19 anti-PER | 1:200 | Santa Cruz Biotechnology, Cat# sc-15720; (Shiga and Numata, 2009); RRID: AB_654018 |
| rabbit anti-PER | 1:1000 | (Stanewsky *et al.*, 1997) RRID: AB_2315105 |
| rabbit anti-AstC | 1:250 | (Park *et al.*, 2008) RRID: AB_2569126 |
| rabbit anti-CRZ | 1:1000 | (Veenstra and Davis, 1993) |
| goat anti-chicken Alexa Fluor® 488 | 1:500 to 1:200 | Life technologies, Carlsbad, CA, USA; Thermo Fisher Scientific |
| goat anti-mouse Alexa Fluor® 647 | 1:500 | Life technologies, Carlsbad, CA, USA |
| goat anti-rabbit Cy3 | 1:500 | Millipore, Billerica, MA, USA |
| goat anti-rat Cy3 | 1:500 | Millipore, Billerica, MA, USA |
| goat anti-guinea pig Alexa Fluor® 647 | 1:400 | Thermo Fisher Scientific |
| goat anti-rabbit Alexa Fluor® 635 | 1:400 | Thermo Fisher Scientific |
| goat anti-mouse Alexa Fluor® 555 | 1:400 | Thermo Fisher Scientific |
| goat anti-human F(ab')2 Alexa Fluor® 488 | 1:200 | Thermo Fisher Scientific |
| donkey anti-guinea pig Alexa Fluor ® 555 | 1:400 | Thermo Fisher Scientific |
| donkey anti-rabbit Alexa Fluor ® 555 | 1:400 | Thermo Fisher Scientific |
| donkey anti-goat Alexa Fluor ® 488 | 1:400 | Thermo Fisher Scientific |
| donkey anti-rabbit Alexa Fluor ® 647 | 1:400 | Thermo Fisher Scientific |
| donkey anti-mouse Alexa Fluor ® 555 | 1:200 | Thermo Fisher Scientific |

**Supplementary Table 3:** FlyWire cell IDs of identified clock neurons
See separate file

**Supplementary Table 4:** FlyWire cell IDs of identified neurosecretory cells (NSC)
See separate file

**Supplementary Table 5:** Hemibrain cell IDs of identified clock neurons
See separate file

**Supplementary Table 6:** Previously published antibody staining, live imaging and ligand specificity data for neuropeptide receptors
See separate file

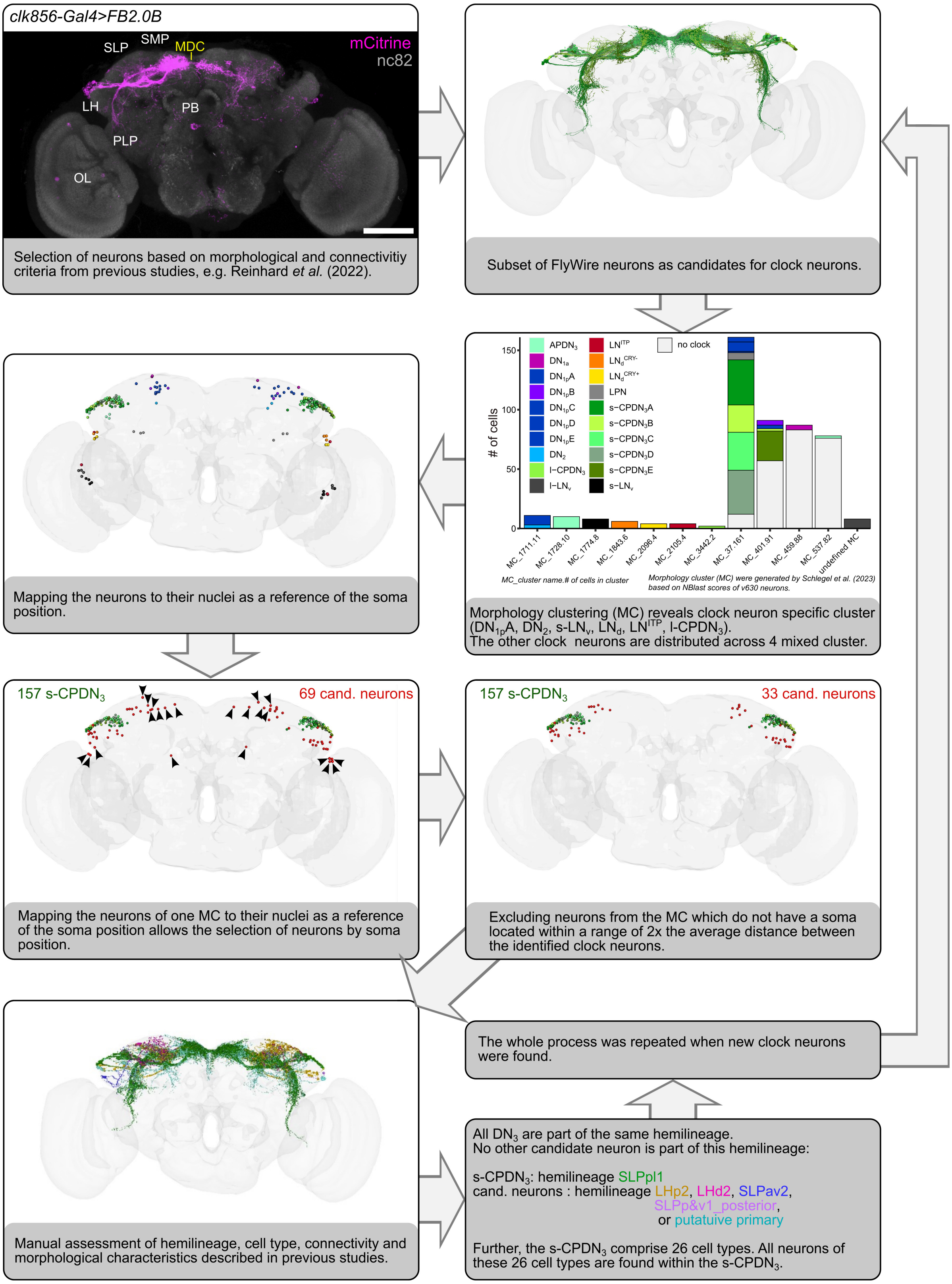


**Figure S1: Pipeline for identifying clock neurons in the FlyWire dataset.** Neurons were identified based on morphological or connectivity features described previously (Lamaze *et al.*, 2018, Schubert *et al.*, 2018, Reinhard *et al.*, 2022a, Reinhard *et al.*, 2022b, Sun *et al.*, 2022) or based on NBLAST similarity to identified clock neurons in the hemibrain (Schlegel *et al.*, 2023). Morphology clustering (v630, (Schlegel *et al.*, 2023)) was then used to identify neurons with similar morphology. Several clock neurons formed clock neuron-specific morphology clusters (e.g. APDN$_3$, s-LN$_v$ and LN$^{ITP}$ amongst others). If clock neurons did not form a unique morphology cluster, all neurons of the corresponding morphology cluster were considered as possible candidates. The cell body position was determined by mapping the coordinates of the nuclei (Mu *et al.*, 2021) to the root ids of the neurons. The average distance between the cell body position of neurons in one clock cluster was determined per hemisphere and candidate neurons laying within twice the average distance (neurons laying further apart are marked by arrowheads) were manual assessed based on morphological features and connectivity described in previous studies. In addition, hemilineage and cell type information was used to determine whether the candidate neurons could be additional clock neurons. If new clock neurons were identified, the whole procedure was repeated. All identified clock neurons include all neurons of the different cell types found within these clock neurons. By definition, a cell type is a uniquely identifiable neuron in the dataset (Schlegel *et al.*, 2023). All numbers refer to neurons across both hemispheres.

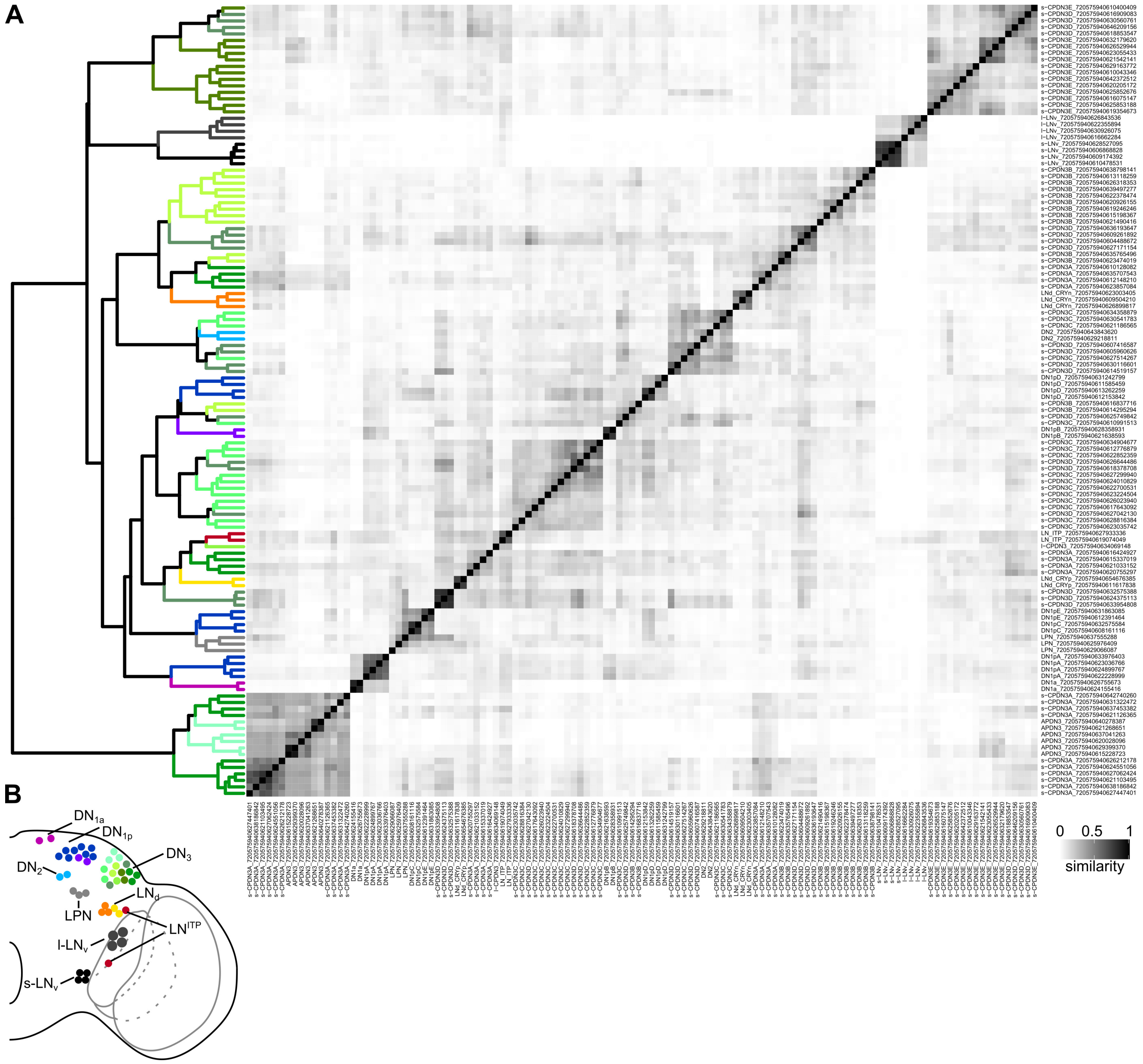


**Figure S2: Clustering of clock neurons based on their total inputs and outputs. (A)** Cosine similarity matrix of clock neurons of the right hemisphere based on their total inputs and outputs from other neurons. For the similarity analysis, no threshold for the number of synapses was used. The darker the color, the higher the similarity between neurons. Neurons within the clades are colored based on the schematic in **(B).** With the exception of $DN_3$ and $DN_{1p}$, all neurons from one clock neuron group cluster together. Note that the $DN_3$ do form $DN_3$-specific clusters but often also cluster with other clock neuron groups.

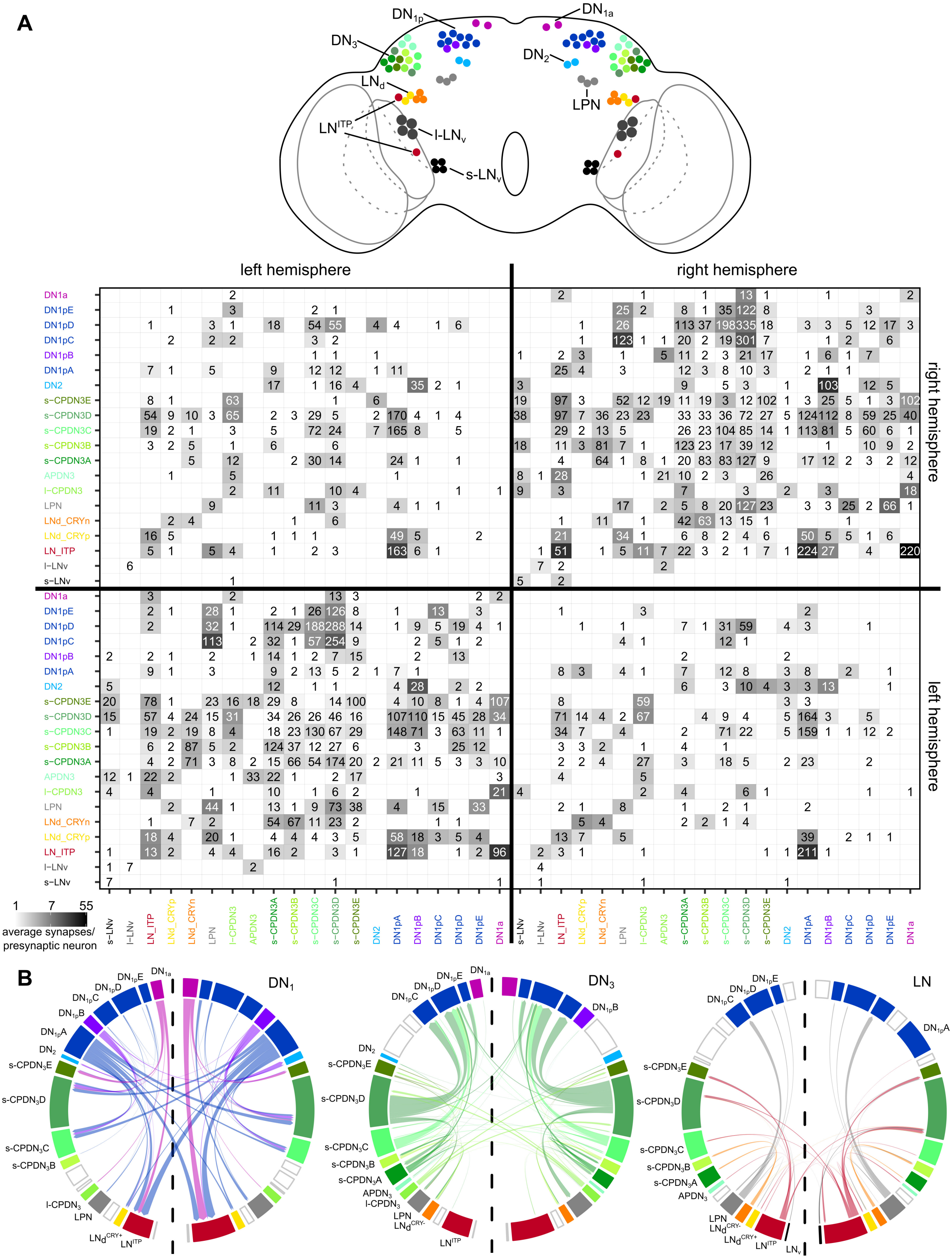


**Figure S3: Connectivity between different clock clusters. (A)** Connectivity matrix highlighting the interconnectivity between different clock cell types across the two hemispheres. Locations of different clock cells are depicted in the schematic. The numbers within the matrix indicate the number of synapses. No synapse threshold was applied. Grey shading reflects the average strength of the connections. **(B)** Clock interconnectivity as shown in Figure 1 separated by $DN_1$, $DN_3$, and LN. Only connections greater than 4 synapses were considered.

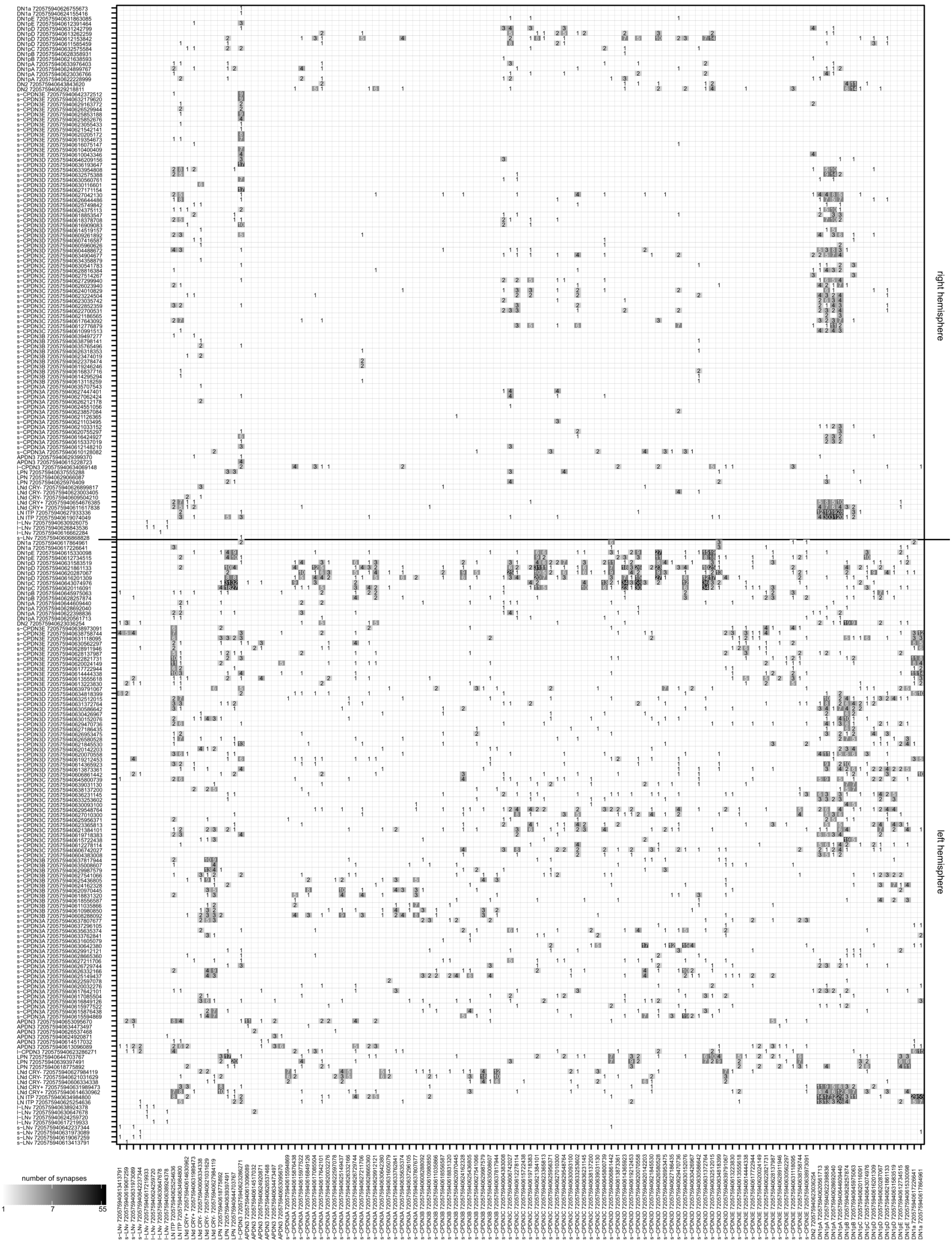


**Figure S4: Connectivity from individual clock cells in the left hemisphere to all clock cells.** Connectivity matrix highlighting the interconnectivity between individual clock cells. Grey shading and the numbers within the matrix indicate the number of synapses per connection. Black indicates strong connectivity and white indicates low or no connectivity strength. No synapse threshold was applied.

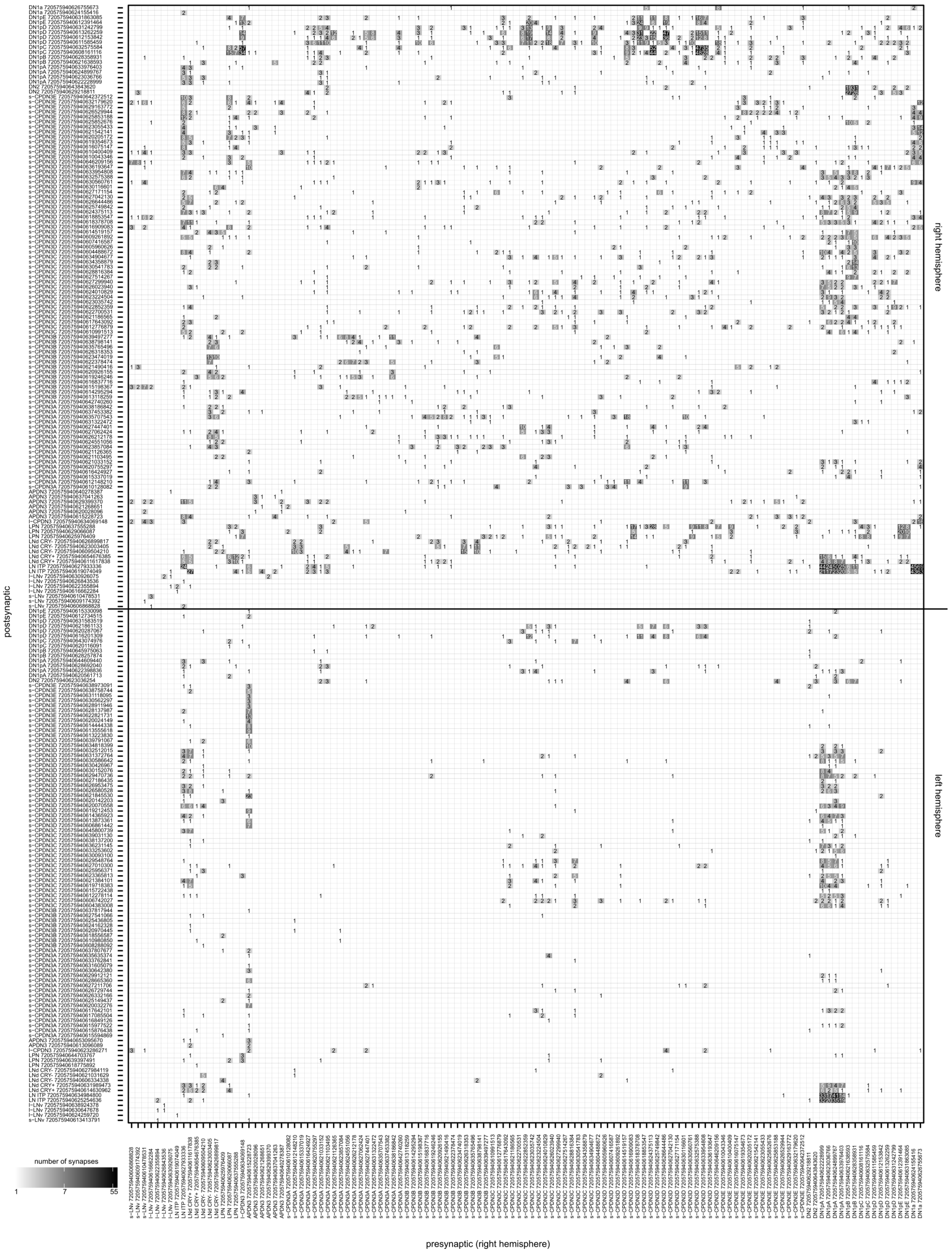


**Figure S5: Connectivity from individual clock cells in the right hemisphere to all clock cells.** Connectivity matrix highlighting the interconnectivity between individual clock cells. Grey shading and the numbers within the matrix indicate the number of synapses per connection. Black indicates strong connectivity and white indicates low or no connectivity strength. No synapse threshold was applied.

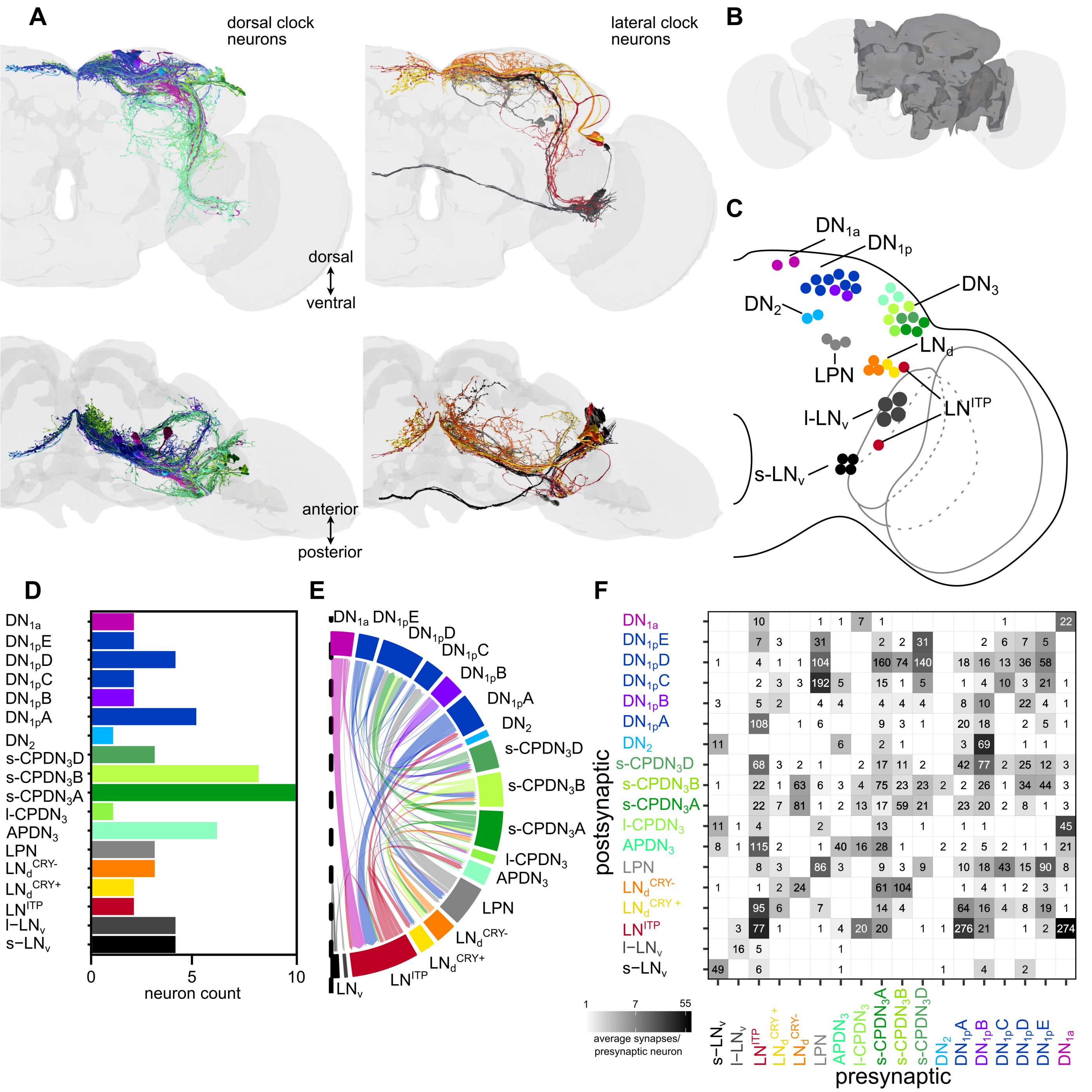


**Figure S6: *Drosophila* hemibrain circadian clock network. (A)** Reconstructions of identified clock neurons in the **(B)** *Drosophila* hemibrain connectome**. (C)** Schematic showing the locations of different clock neuron cell types. **(D)** Numbers of the different clock cell types that were identified in the hemibrain connectome. **(E)** Synaptic interconnectivity between different cell types within the clock network. The direction of the arrow indicates the flow of information. Only connections with >9 synapses were taken into account. The dashed line indicates the brain midline. **(F)** Connectivity matrix highlighting the interconnectivity between different clock cell types. The numbers within the matrix indicate the number of synapses. Grey shading reflects the average strength of this connectivity. No synapse threshold was applied.

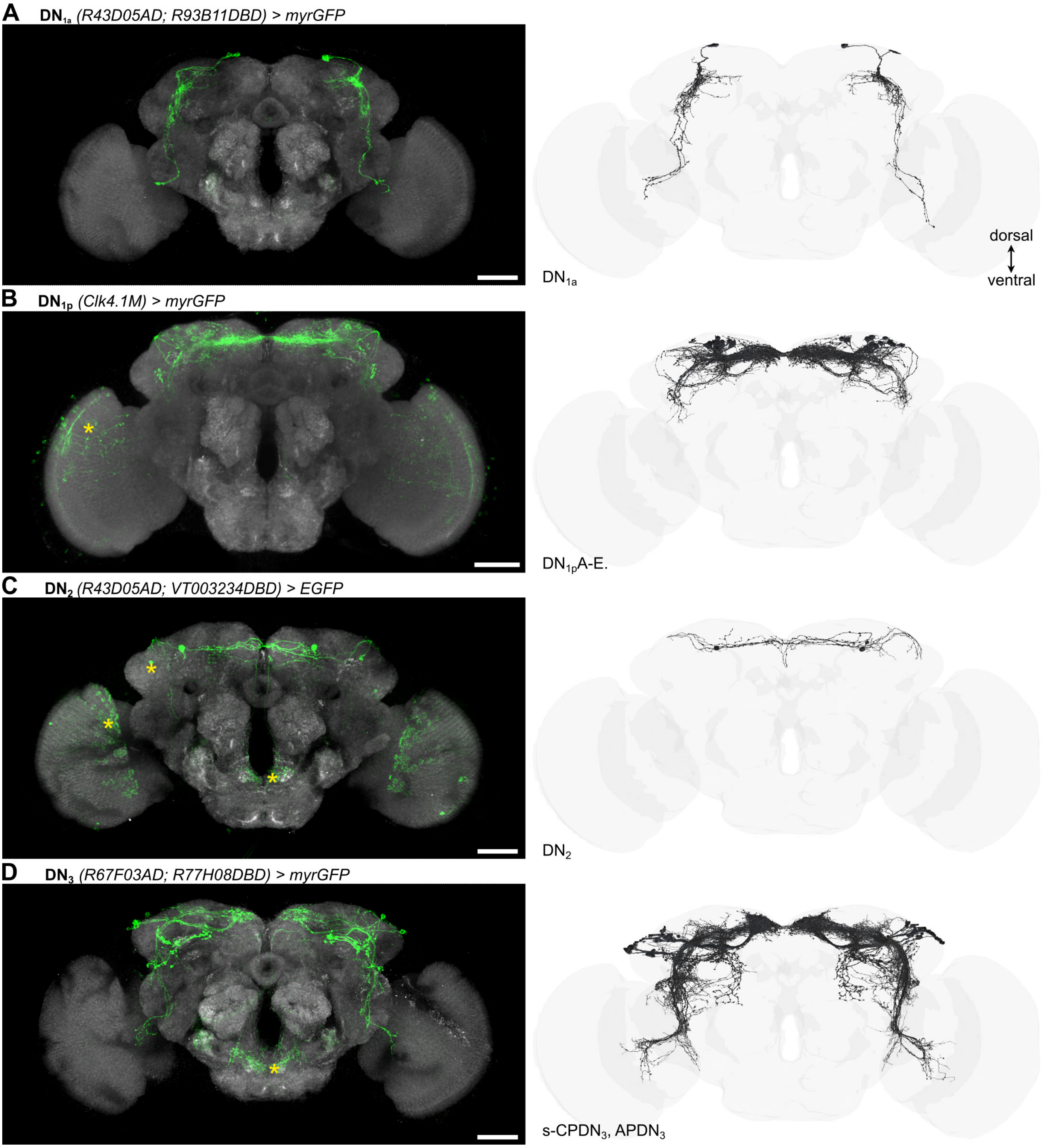


**Figure S7: Expression patterns of Gal4 lines targeting different populations of DN.** Confocal stacks showing GFP expression in **(A)** $DN_{1a}$, **(B)** $DN_{1p}$, **(C)** $DN_2$, and **(D)** $DN_3$ that are included in the different Gal4 lines. Reconstructions of corresponding neurons in the FlyWire connectome are provided on the right. Neurons not part of the clock network are marked by an asterisk. Scale bars = 50µm.

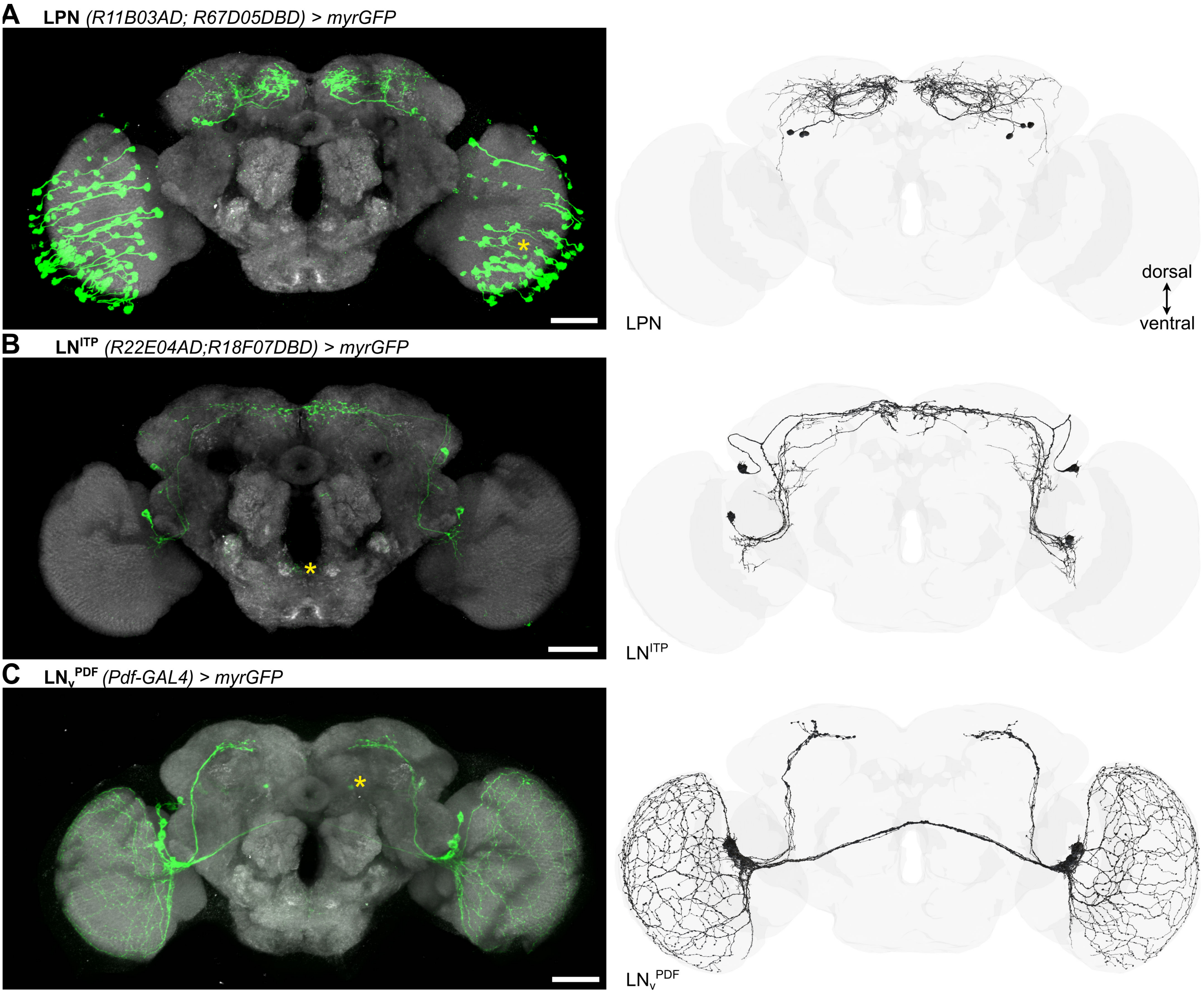


**Figure S8**: **Expression patterns of Gal4 lines targeting different populations of LN.** Confocal stacks showing GFP expression in **(A)** LPN, **(B)** $LN^{ITP}$, and **(C)** $LN_v^{PDF}$ that are included in the different Gal4 lines. Reconstructions of corresponding neurons in the FlyWire connectome are provided on the right. Neurons not part of the clock network are marked by an asterisk. Scale bars = 50μm.

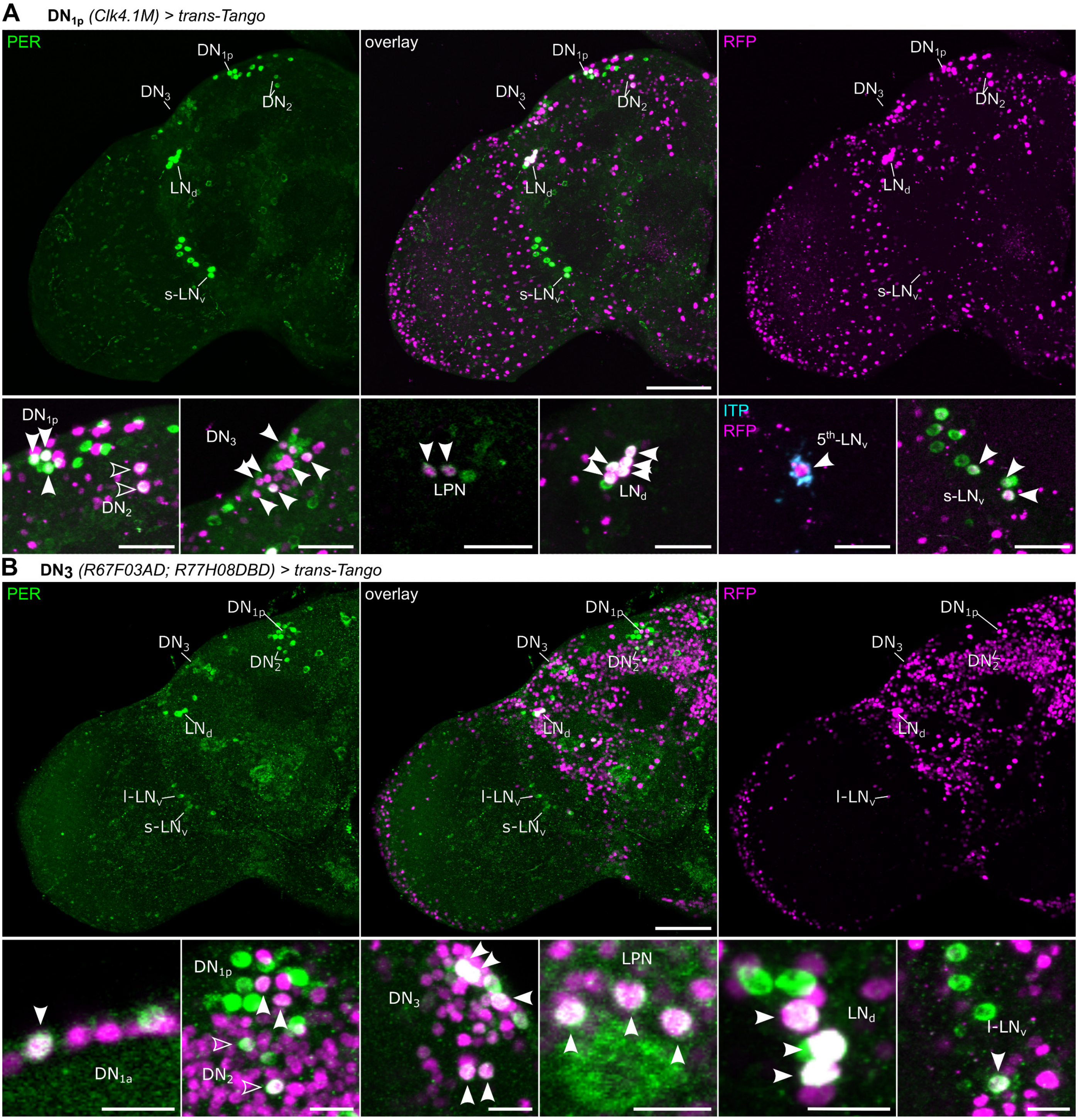


**Figure S9: Validating connectivity within the clock network using *trans*-Tango. (A)** Driving *trans*-Tango using *Clk4.1M-Gal4* revealed $DN_{1p}$, $DN_2$, $DN_3$, LPN, $LN_d$, $5^{th}$-$LN_v$, and s-$LN_v$ as post-synaptic partners of $DN_{1p}$. **(B)** Expressing *trans*-Tango in a subset of the $DN_3$ (including s-CP$DN_3$ and AP$DN_3$) generated a post-synaptic signal in all dorsal clock neuron clusters, as well as LPN, $LN_d$, and l-$LN_v$. Scale bars = 50µm for overview and 20µm for higher magnification images. These two examples highlight a general agreement in synaptic connectivity observed with the connectomes and *trans*-Tango. Abbreviations: PER, Period; ITP, Ion transport peptide; RFP, red fluorescent protein.

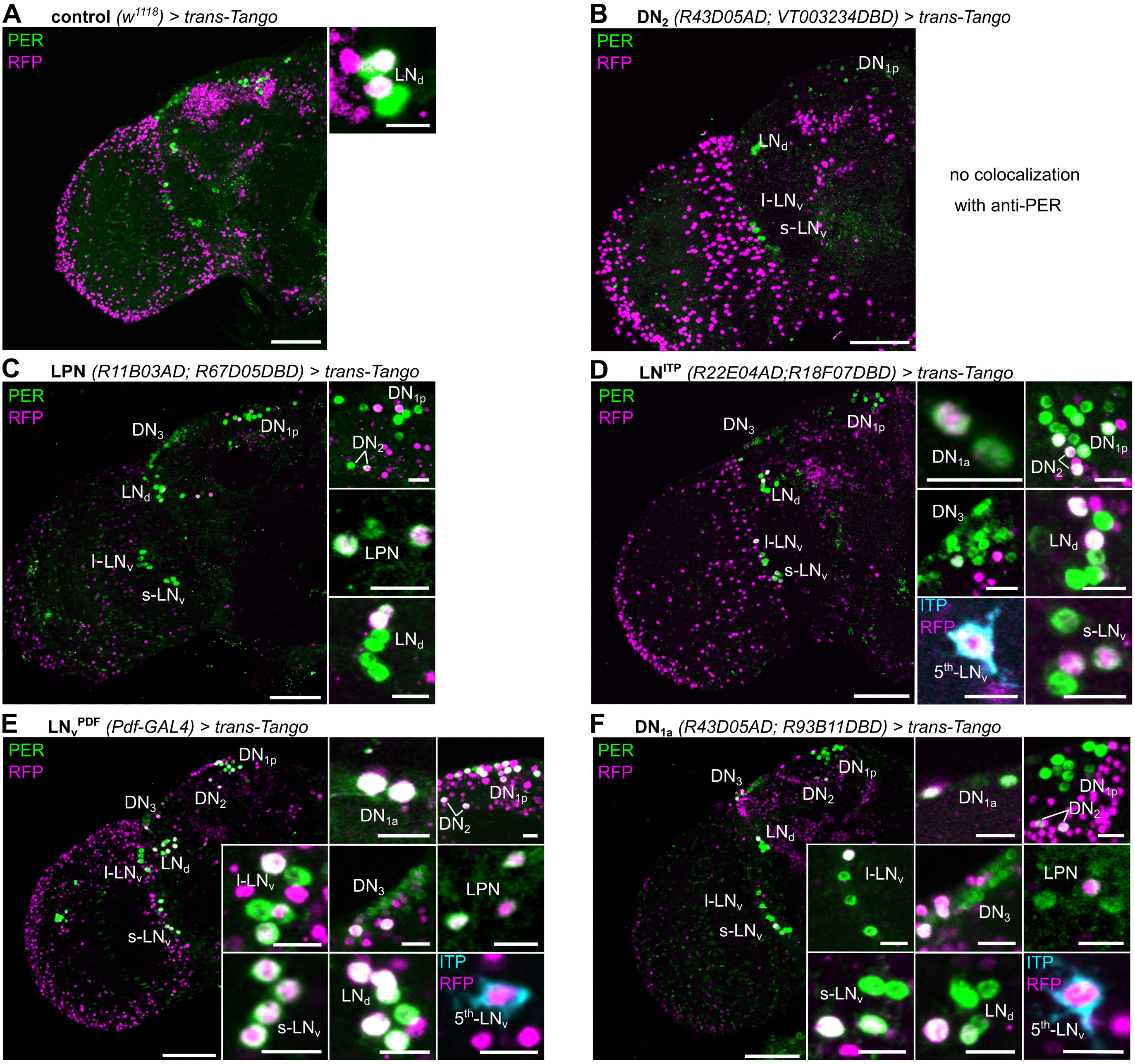


**Figure S10**: **Validating connectivity within the clock network using *trans*-Tango. (A)** Control for *trans*-Tango occasionally generated a false post-synaptic signal in two $LN_d$. Expressing *trans*-Tango in **(B)** $DN_2$ does not generate a post-synaptic signal in any clock neurons but **(C)** LPN, **(D)** $LN^{ITP}$, **(E)** $LN_V^{PDF,}$ and **(F)** $DN_{1a}$ are all presynaptic to other clock neurons. Scale bars = 50µm for overview and 20µm for higher magnification images. Abbreviations: PER, Period; ITP, Ion Transport Peptide; RFP, Red Fluorescent Protein.

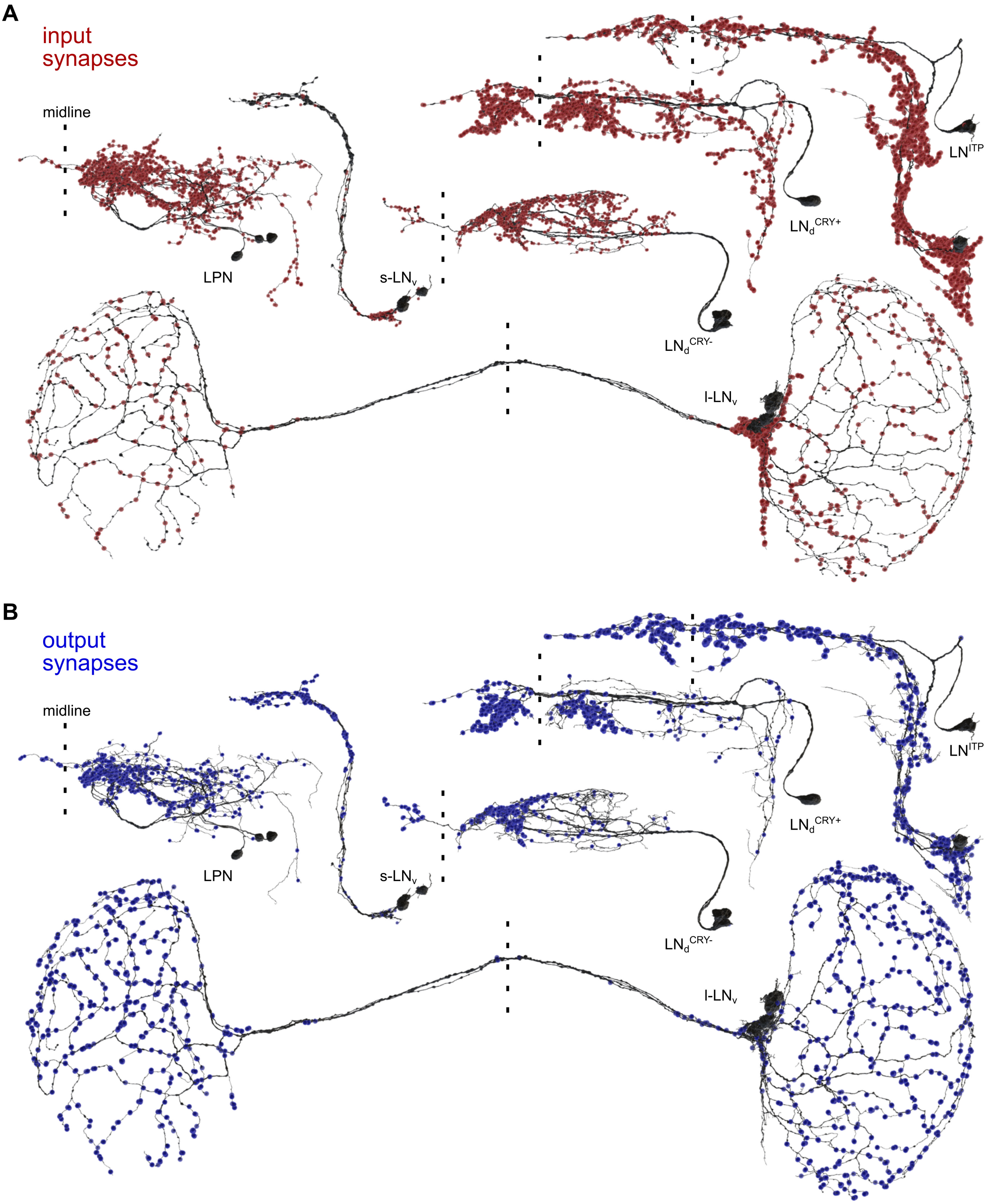


**Figure S11: Synapse locations of lateral clock neurons. (A)** Postsynaptic (red) and **(B)** presynaptic (blue) sites of different groups of lateral clock neurons in the right hemisphere. The dashed line indicates the brain midline. Distinguishable dendrites are seen seldomly.

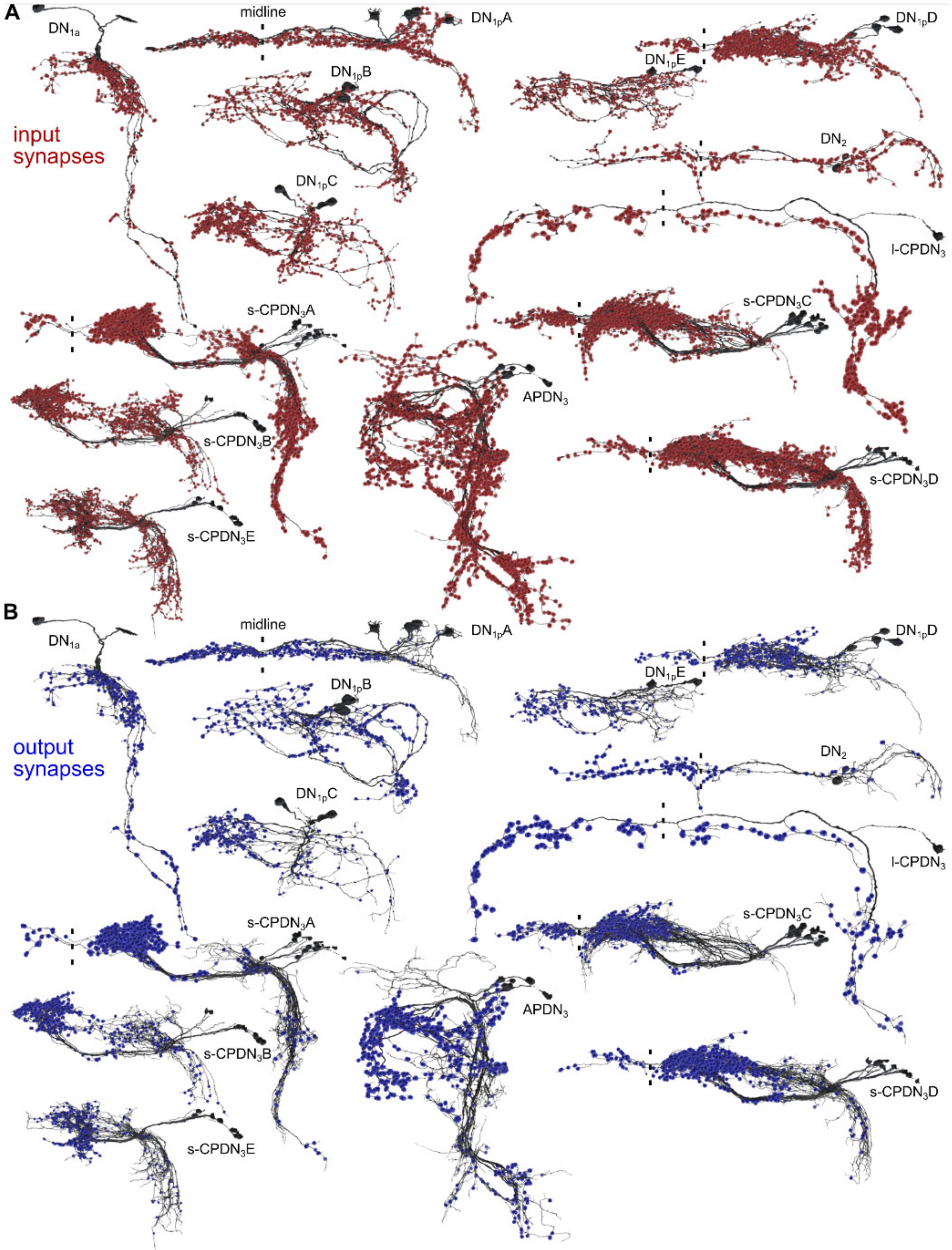


**Figure S12: Synapse locations of dorsal clock neurons. (A)** Postsynaptic (red) and **(B)** presynaptic (blue) sites of different groups of dorsal clock neurons in the right hemisphere. The dashed line indicates the brain midline. Distinguishable dendrites are seen seldomly.

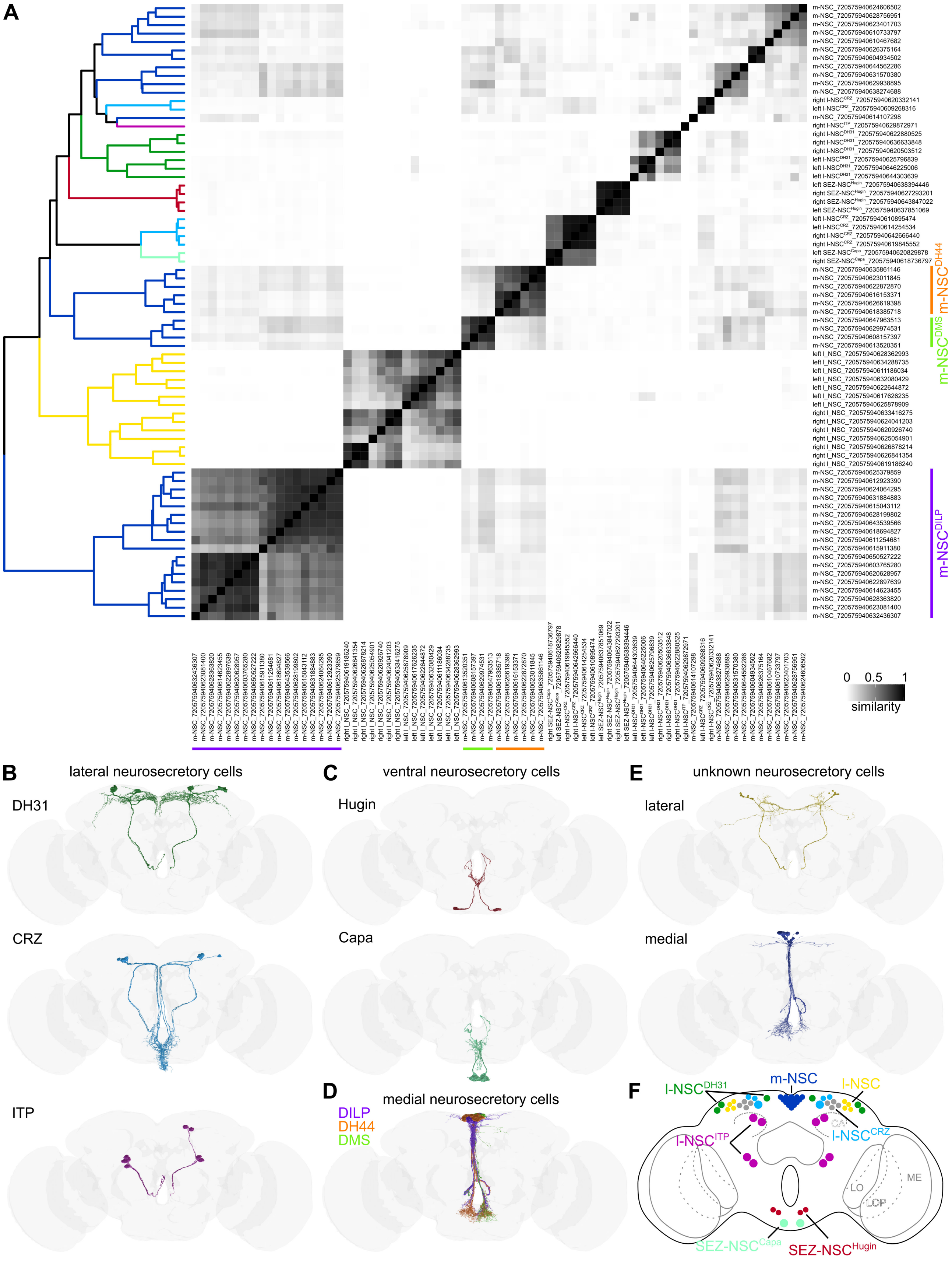


**Figure S13**: **Clustering of neurosecretory cells based on total inputs. (A)** Cosine similarity matrix of all neurosecretory cells (NSC) based on their total inputs. The darker the color, the higher the similarity between neurons. Neurons within the clades are colored based on the schematic in **(F).** Putative m-NSC$^{DILP}$, m-NSC$^{DH44,}$ and m-NSC$^{DMS}$ clusters resolved by this analysis are marked. Reconstructions of **(B)** DH31, CRZ, and ITP lateral NSC, **(C)** Hugin and Capa subesophageal NSC, **(D)** DILP, DH44, and DMS medial NSC, and **(E)** other lateral and medial NSC with unidentified neurohormone. **(F)** Schematic showing the locations of different NSC types.

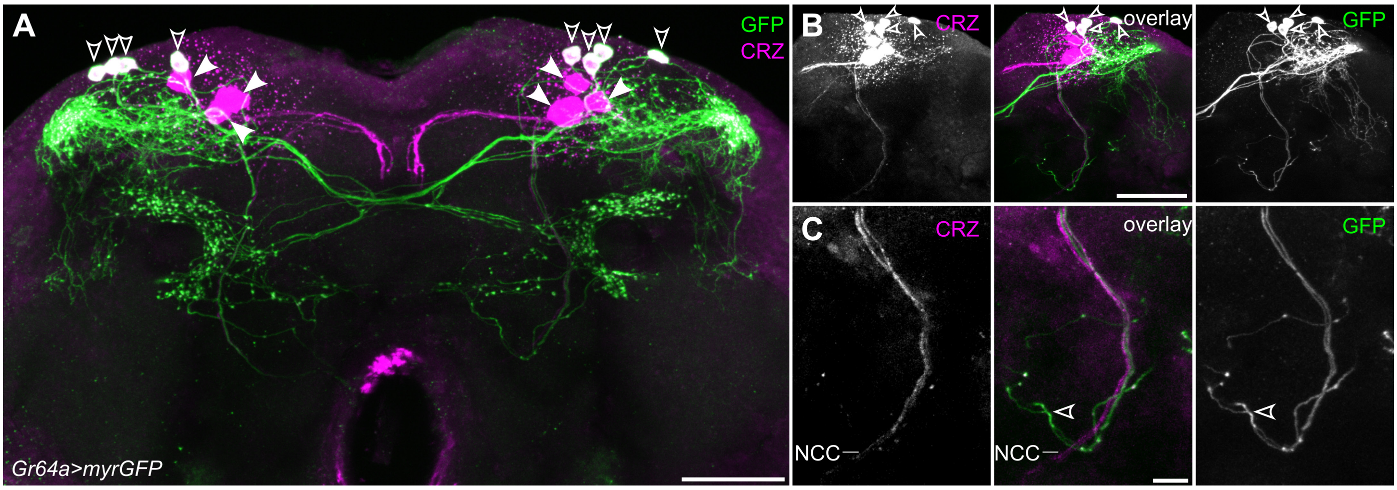


**Figure S14: Anatomical characterization of Corazonin (CRZ) neurosecretory cells. (A)** CRZ is expressed in 7 pairs of neurons in adult flies, 4 of which co-express *Gr64a* (empty arrowheads) **(B)**. These smaller *Gr64a*-expressing CRZ neurons form dense arborizations in the lateral horn. They project contralaterally but do not send projections with the nervii corpora cardiaca (NCC) **(C)** and are thus not considered neurosecretory. Scale bars = 50μm for overview and 20μm for higher magnification images.

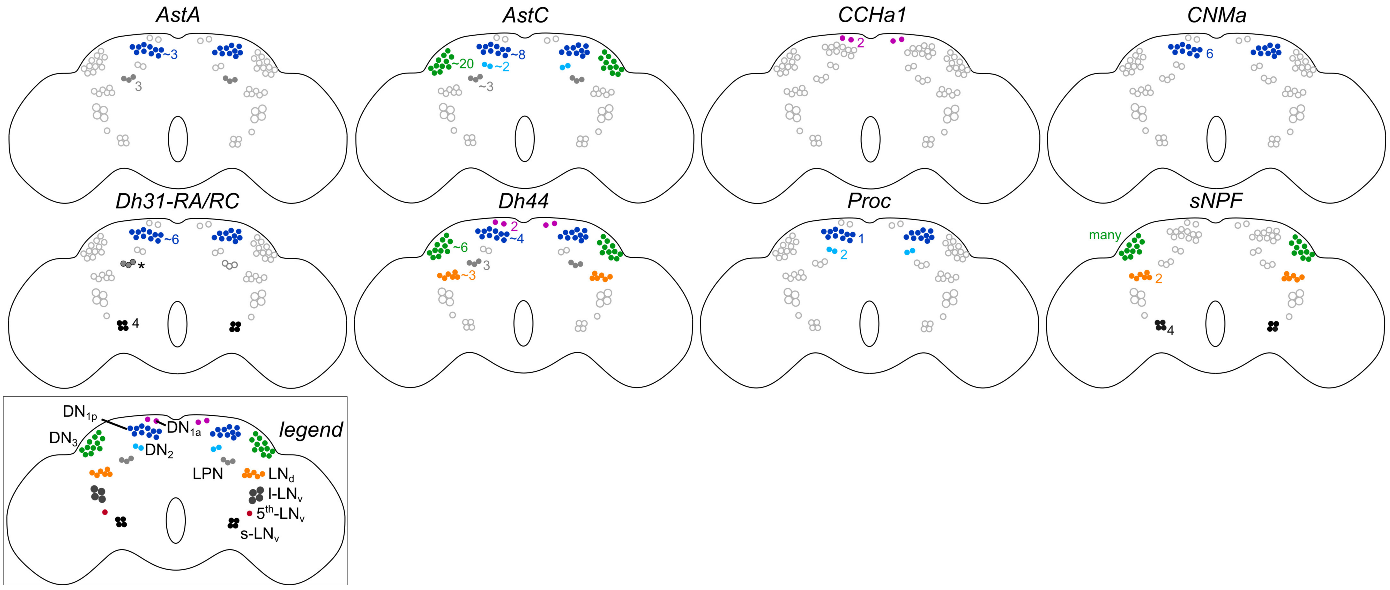


**Figure S15: Neuropeptide expression in clock neurons.** Schematics showing the clock neurons that express different neuropeptides. Schematics are based on expression mapping reported in Figure S16. The numbers of clock cells of a given type that express the neuropeptide are also included where appropriate. DH31 expression schematic is based on GFP expression in Figure S16 and antibody staining reported earlier (marked by an asterisk) (Reinhard *et al.*, 2022a). Numbers refer to expression in one hemisphere.

**Figure S16: Neuropeptide expression in clock neurons.** Confocal stacks showing expression of GFP (specifically in clock neurons) driven by different neuropeptide-T2A-Gal4 lines. Arrowheads indicate GFP-expressing clock neurons. For all Gal4 lines, panel (**A**) shows a brain overview, and subsequent panels show detailed images of clock neurons. Scale bars = 100µm for overview and 20µm for detail images. Abbreviations: TIM, Timeless; PDF, Pigment dispersing factor; PER, Period.

See separate file

**Figure S17: Neuropeptide receptor expression in clock neurons.** Confocal stacks showing expression of GFP (specifically in clock neurons) driven by different neuropeptide receptor-T2A-Gal4 lines. Arrowheads indicate GFP-expressing clock neurons. For all Gal4 lines, panel **(A)** shows a brain overview, and subsequent panels show detailed images of clock neurons. Scale bars = 100µm for overview and 20µm for detail images. Abbreviations: TIM, Timeless; PDF, Pigment dispersing factor.

See separate file

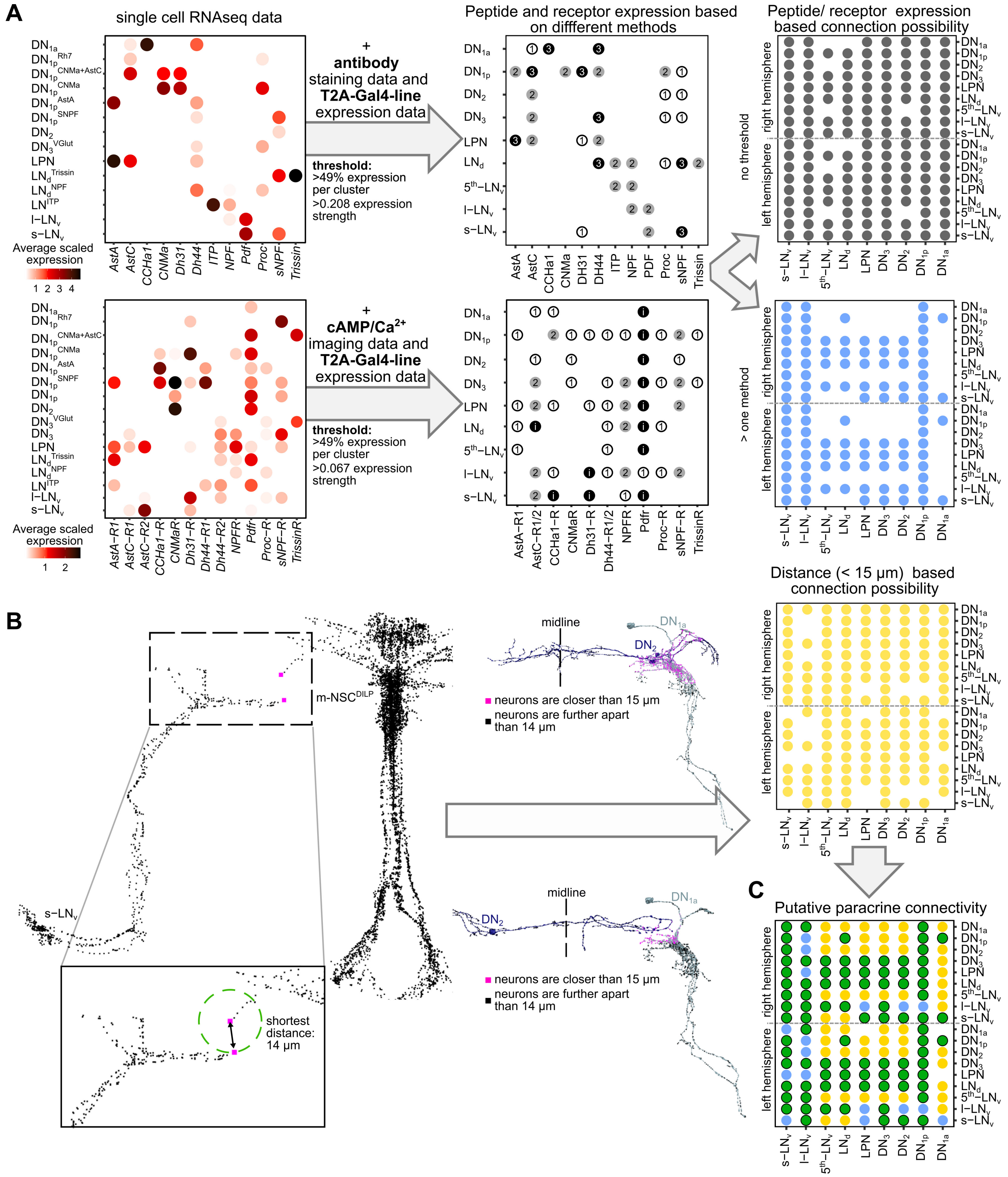


**Figure S18: Workflow to determine putative paracrine connectivity within the clock network. (A)** Expression patterns reported here and in previous studies (see Supplementary Table 6) were used to determine a threshold for the single-cell RNA sequencing data (presented in Figures 6 and 7). 0.208 averaged scaled expression for neuropeptides and 0.067 for receptors was used as a threshold. Additionally, expression in less than 50% of the cells in a given cluster was considered absent. This RNA expression was combined with T2A-Gal4 expression data (Figure S15-17, Figure 7) and antibody or imaging data (i) (see Supplementary Table 6), to generate a peptide/receptor expression map which also takes into account the number of methods that report positive expression. The peptide and receptor expression maps were used to compute a connectivity map. In the absence of any thresholding, all clock clusters communicate with almost all the other clusters via paracrine signaling. To reduce the number of false positives, we only considered expression of peptides and receptors when shown by two independent methods. This filtering reduces the number of putative paracrine connections to some extent (blue). **(B)** The shortest distance between s-LN$_v$ and m-NSC$^{DILP}$ was used as the distance threshold. As shown previously, PDF from the s-LN$_v$ can signal to the m-NSC$^{DILP}$ (Nagy *et al.*, 2019). For all neuron groups, the distance of the neurons to other neuron groups was calculated. If the distance between them was below the threshold of 15µm, these neurons were considered to be in the range for paracrine signaling (yellow). **(C)** Combining the expression and distance matrices allows the calculation of a hypothetical connection matrix for paracrine connectivity (green).

**Video S1:** Reconstructions of single clock neurons from the FlyWire dataset displayed by cluster.

**Video S2:** Reconstructions of all clock neurons from the FlyWire dataset.

**Video S3:** All $DN_3$ are labeled by *tim-(UAS)-Gal4* and could be counted using antibody staining against Period (PER) and Vrille (VRI).

**Video S4:** Reconstructions of all neurosecretory cells from the FlyWire dataset.

**Video S5:** Reconstructions of single neurosecretory cells from the FlyWire dataset displayed by cluster.
