## Supplementary Figure 16 for "Synaptic connectome of the *Drosophila* circadian clock"

**Figure S16: Neuropeptide expression in clock neurons.** Confocal stacks showing expression of GFP (specifically in clock neurons) driven by different neuropeptide-T2A-Gal4 lines. Arrow heads indicate GFP-expressing clock neurons. For all Gal4 lines, panel A shows brain overview and subsequent panels show detail images of clock neurons. Scale bars = 100 µm for overview and 20 µm for detail images. Abbreviations: TIM, Timeless; PDF, Pigment dispersing factor; PER, Period.

#### *AstA-Gal4*

**$DN_{1p}$, LPN**

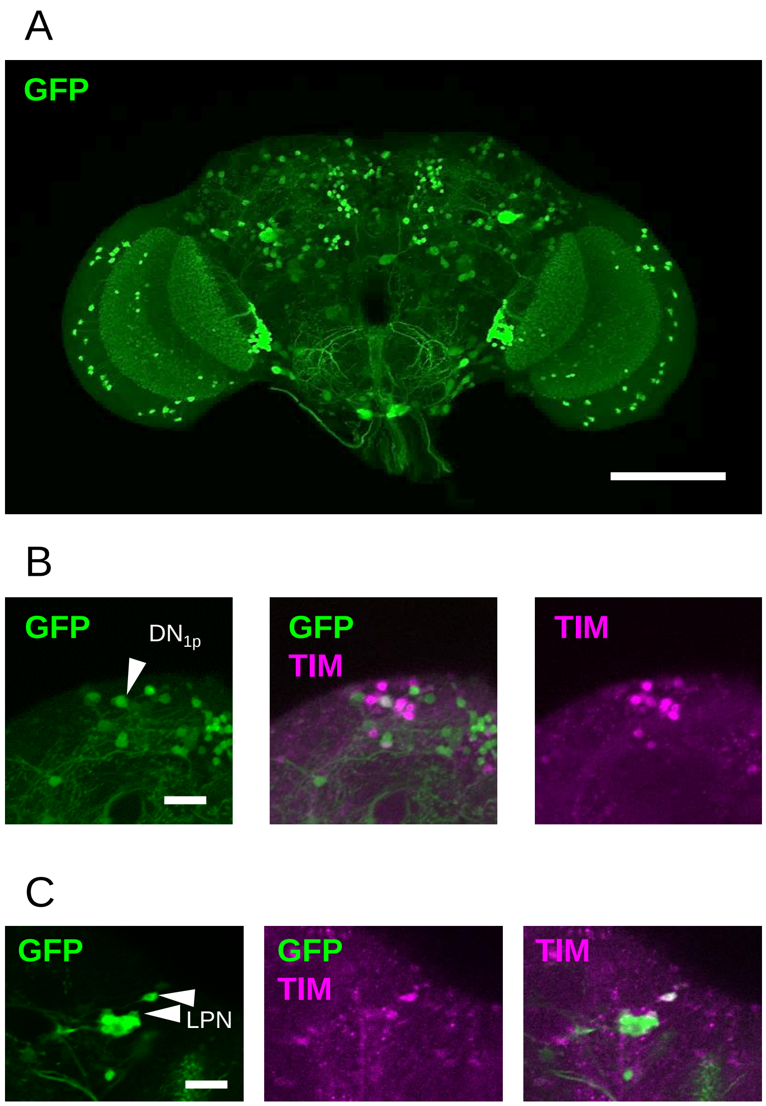

***AstC-Gal4***

**$DN_{1p}$, $DN_2$, $DN_3$, LPN**

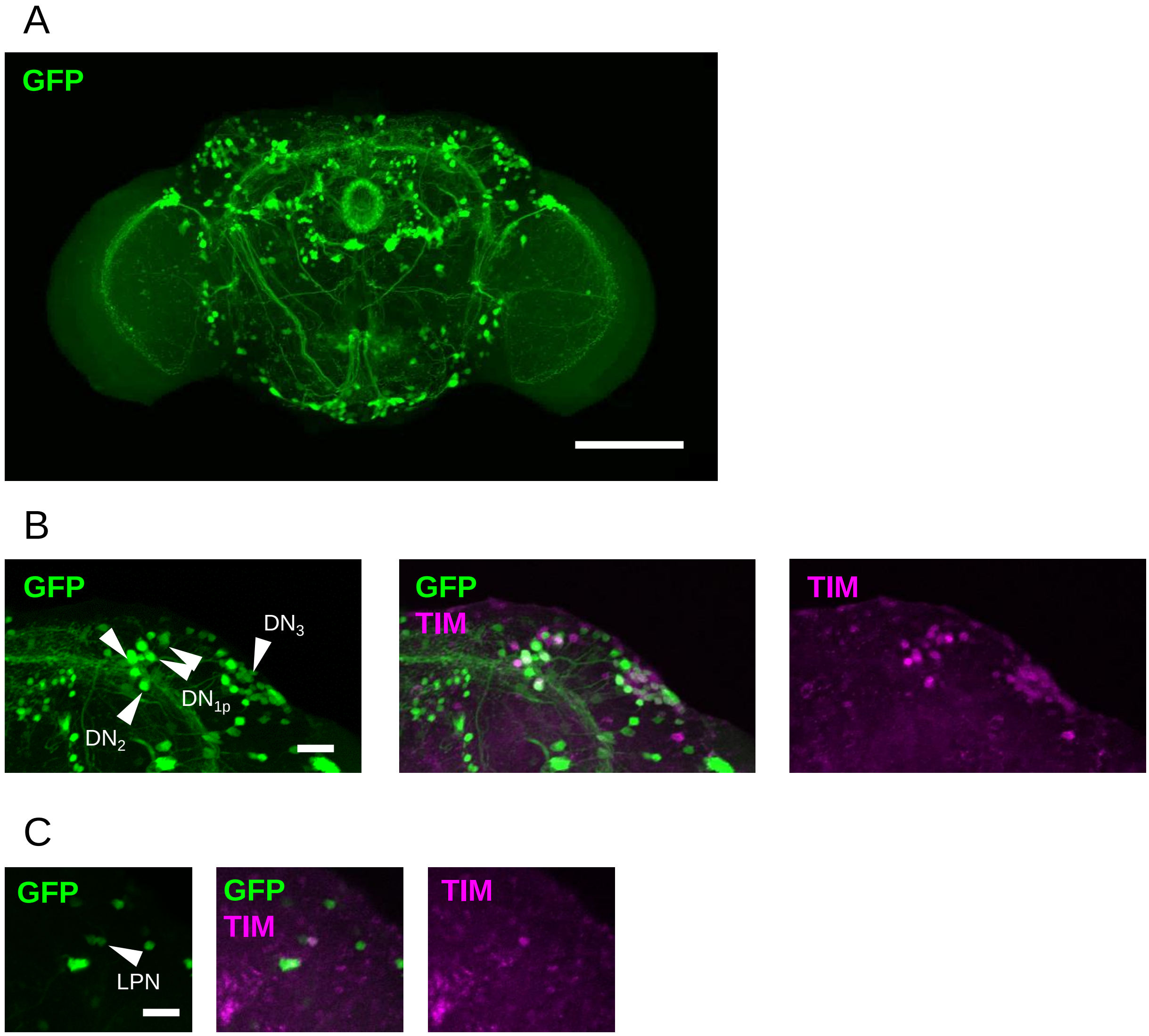

***CCHa1-LexA***

**$DN_{1a}$**

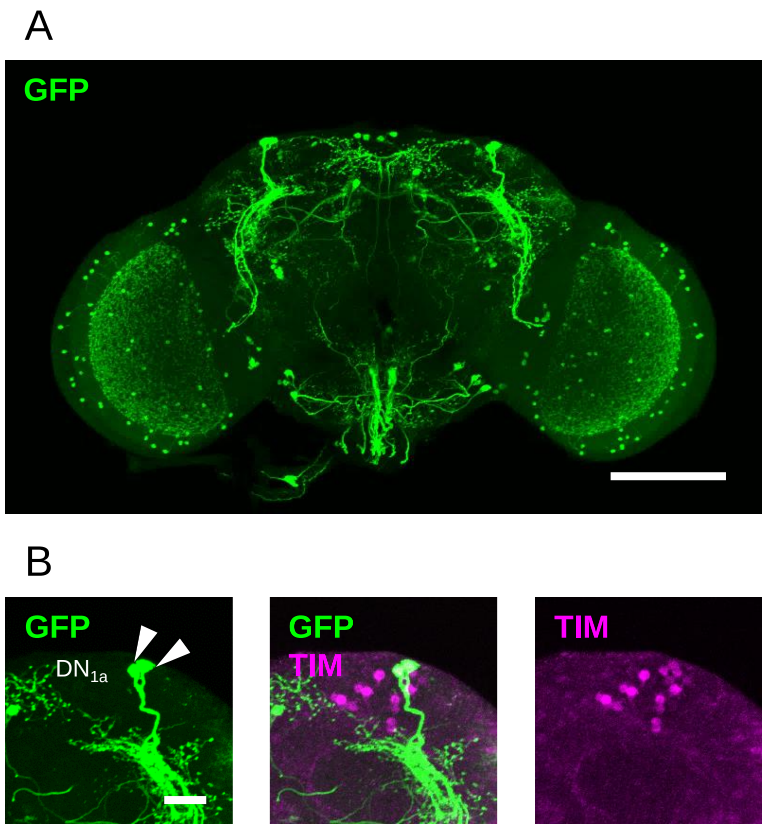

***CNMa-Gal4***

**$DN_{1p}$**

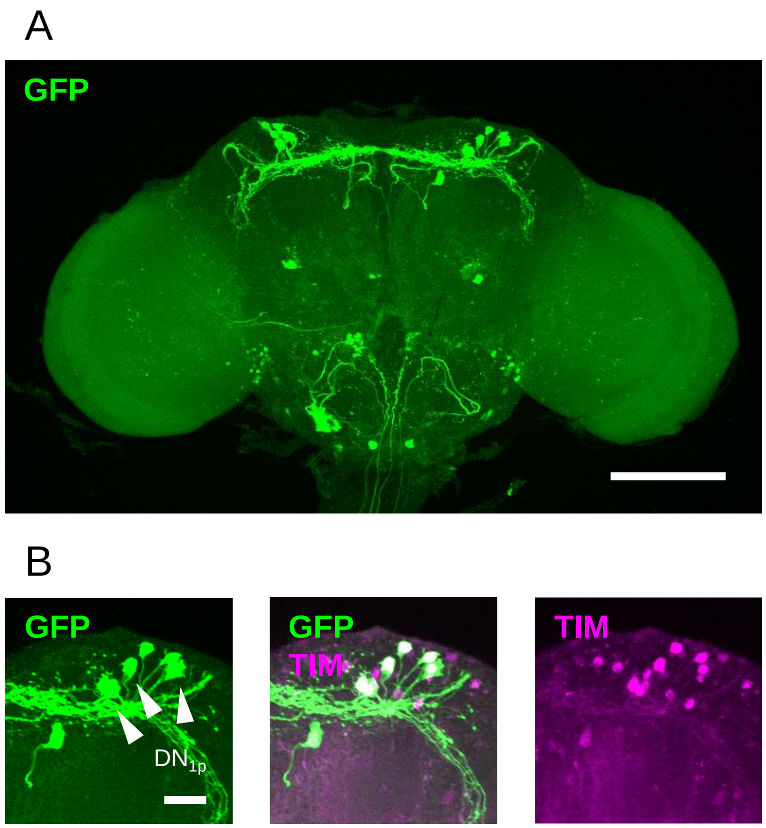

#### *Dh31-RA/C-Gal4*
## $DN_{1p}$, s-$LN_v$

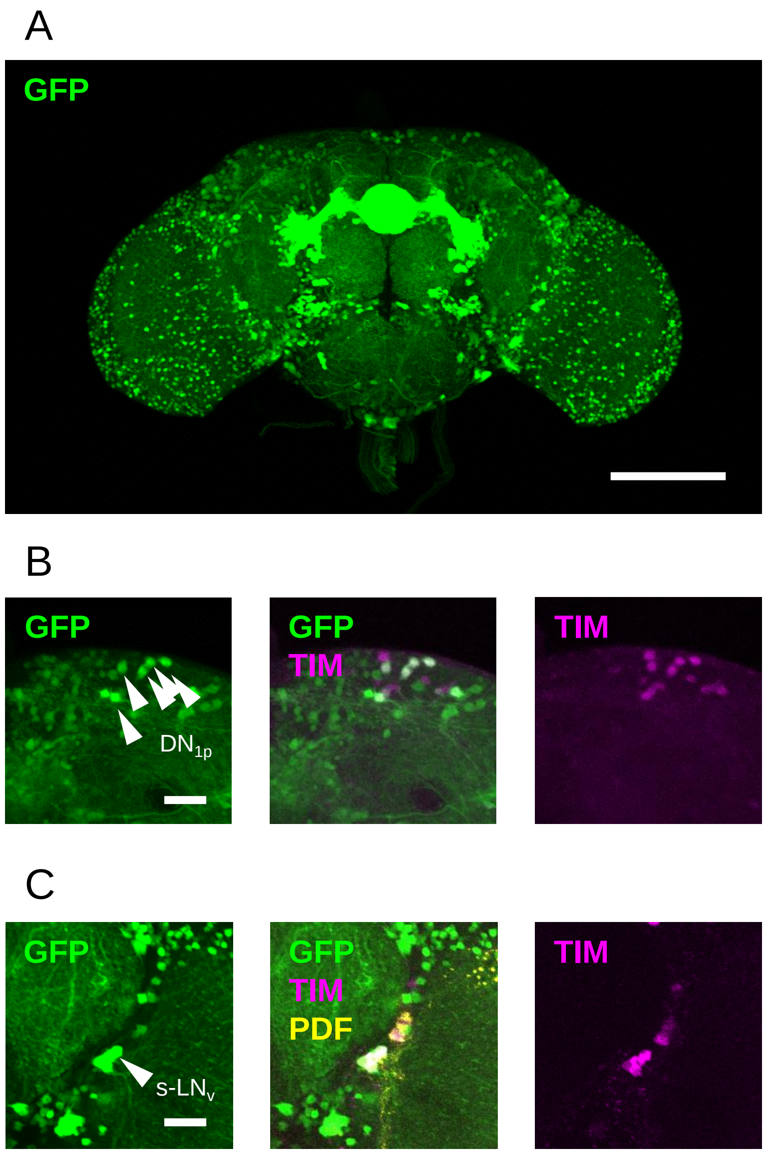

### *Dh44-Gal4*

## $DN_{1a}$, $DN_{1p}$, $DN_3$, $LN_d$

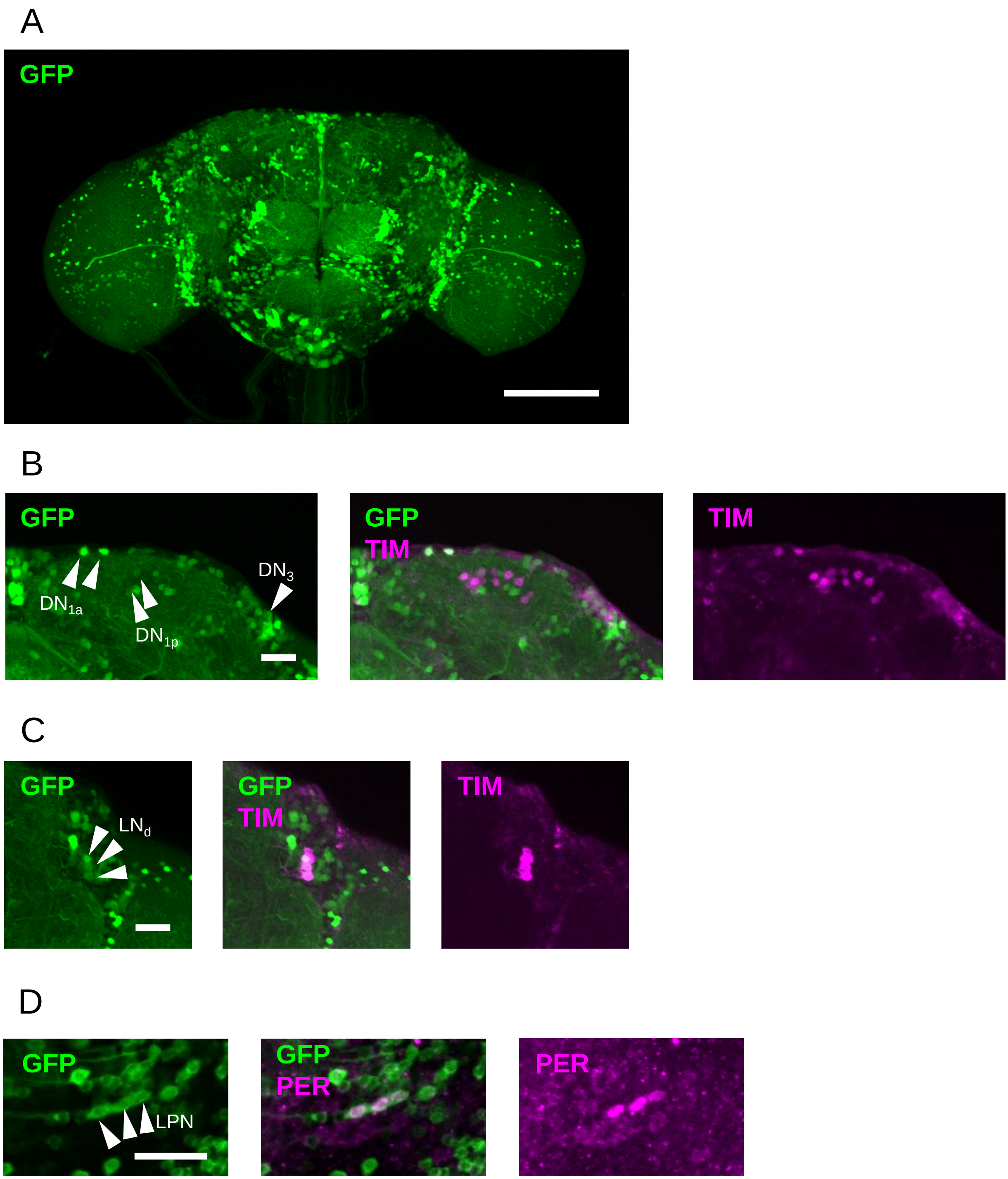

***Proc-LexA***

**$DN_{1p}$**

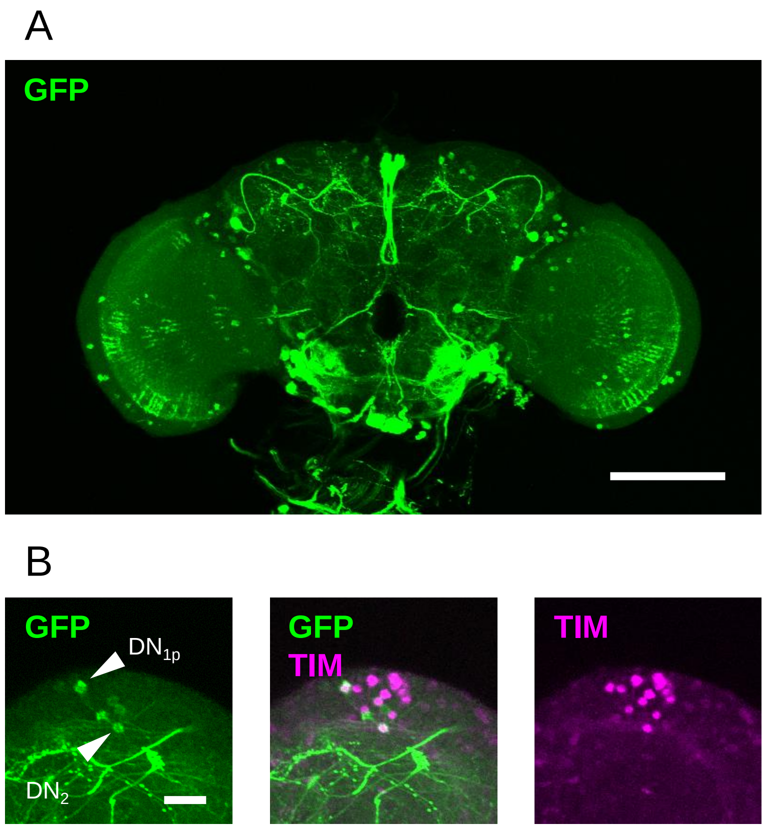

### *sNPF-Gal4*
## $DN_3$, $LN_d$, s-$LN_v$

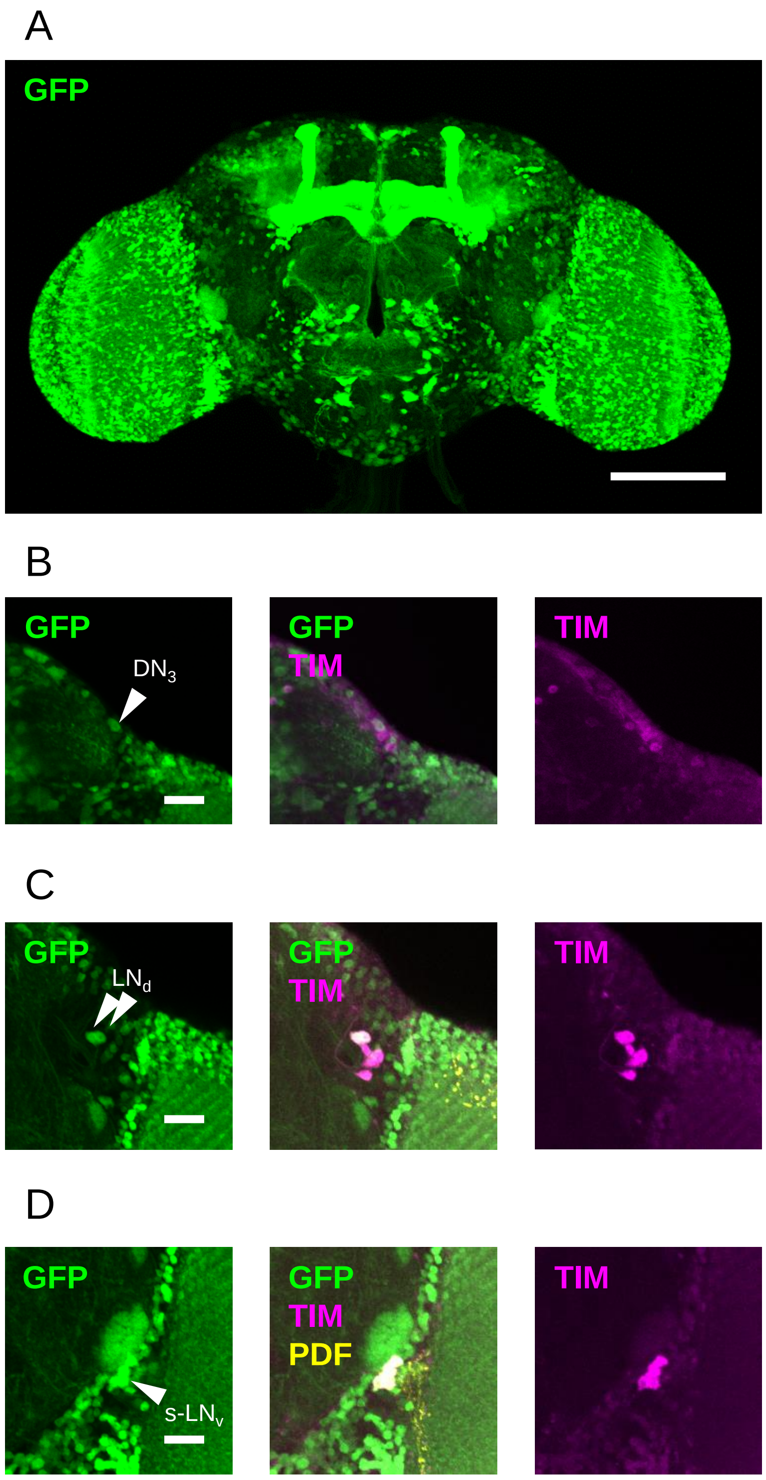
