## Supplementary Figure 17 for "Synaptic connectome of the *Drosophila* circadian clock"

**Figure S17: Neuropeptide receptor expression in clock neurons.** Confocal stacks showing expression of GFP (specifically in clock neurons) driven by different neuropeptide receptor-T2A-Gal4 lines. Arrow heads indicate GFP-expressing clock neurons. For all Gal4 lines, panel A shows brain overview and subsequent panels show detail images of clock neurons. Scale bars = 100 µm for overview and 20 µm for detail images. Abbreviations: TIM, Timeless; PDF, Pigment dispersing factor.

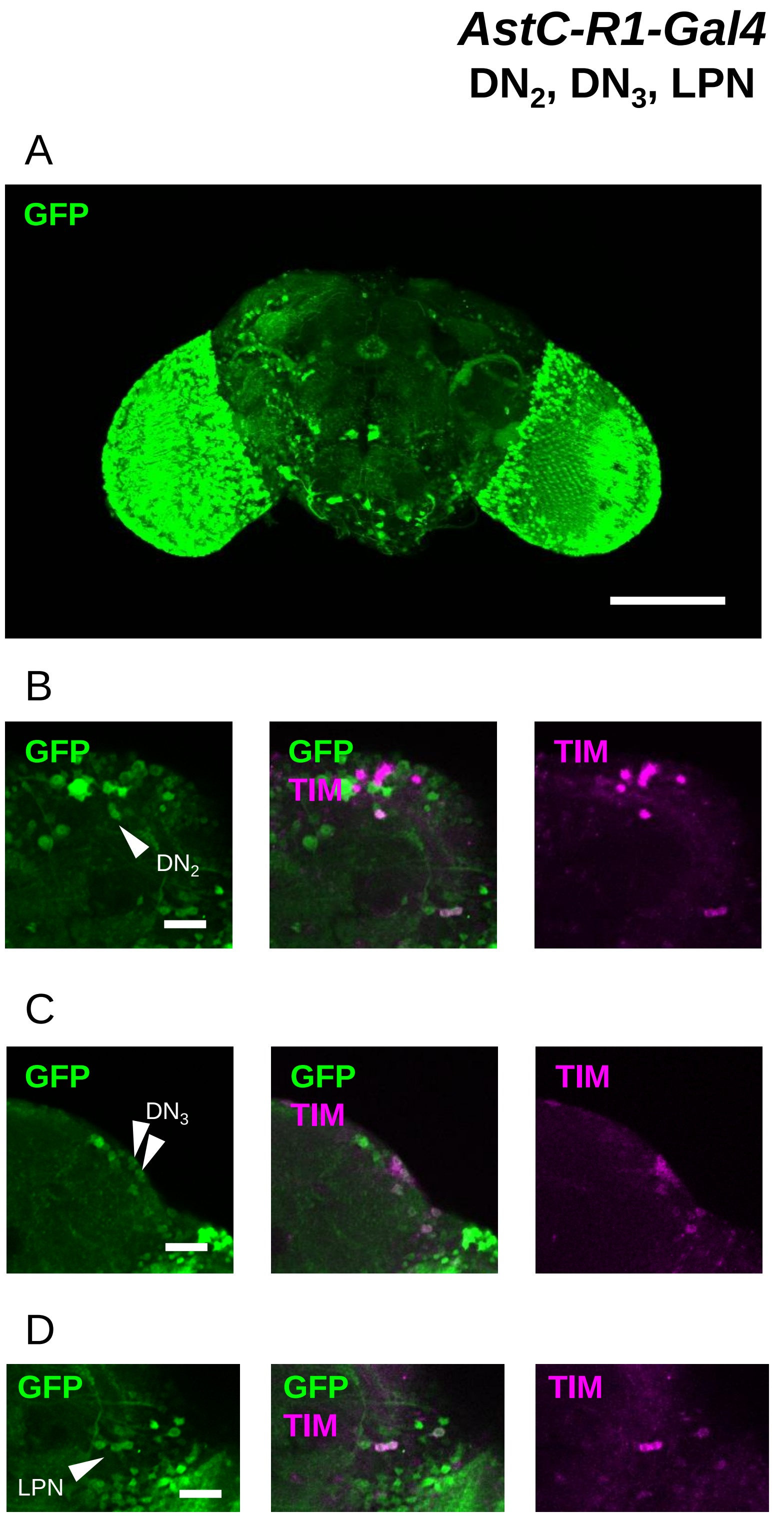

#### *AstC-R2-RB-Gal4*

## $DN_{1a}$, $DN_2$, $DN_3$, $LN_d$, s-$LN_v$, l-$LN_v$

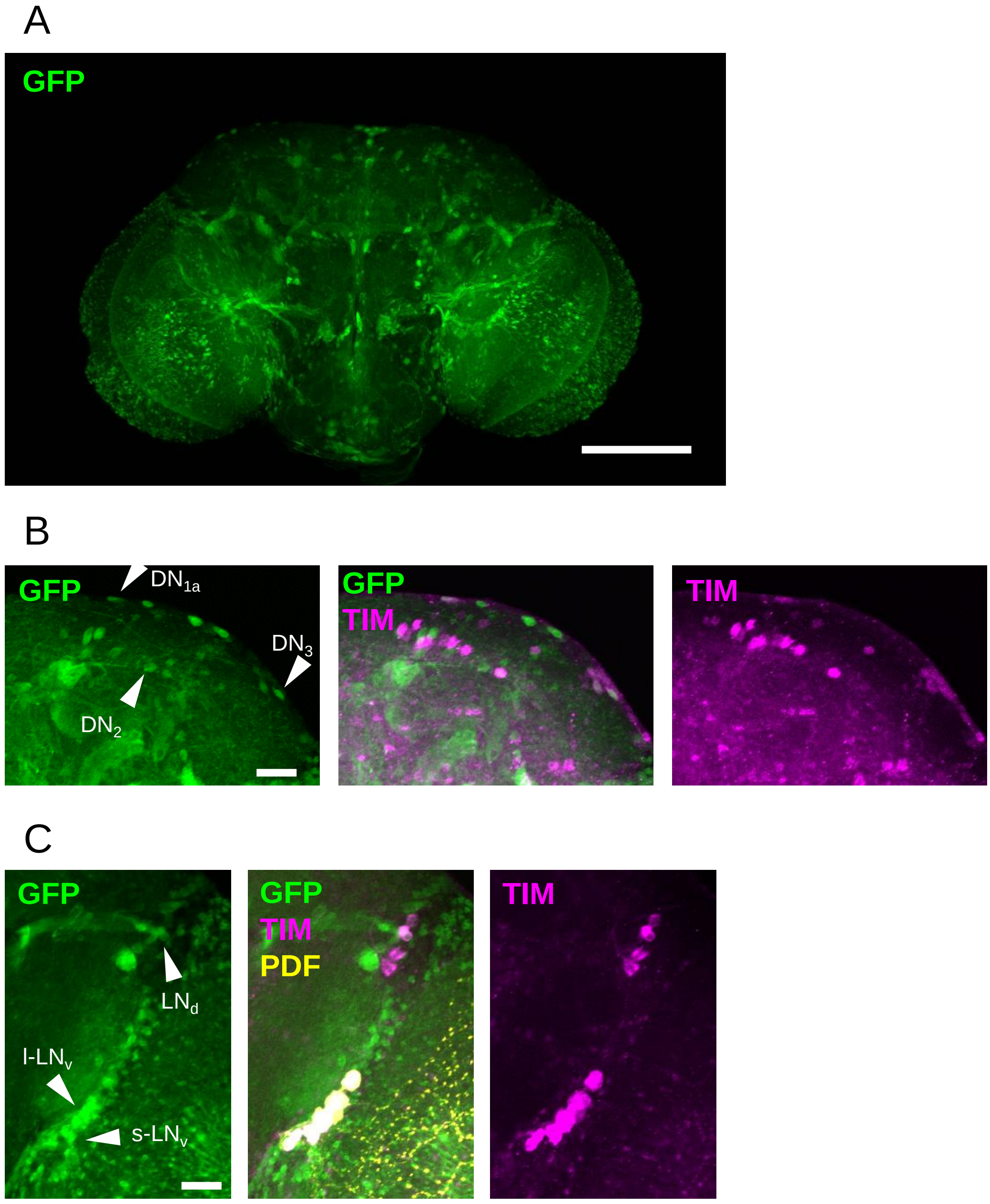

#### *CCHa1R-Gal4*

#### $DN_{1a}$, $DN_{1p}$, LPN, s-$LN_v$, l-$LN_v$

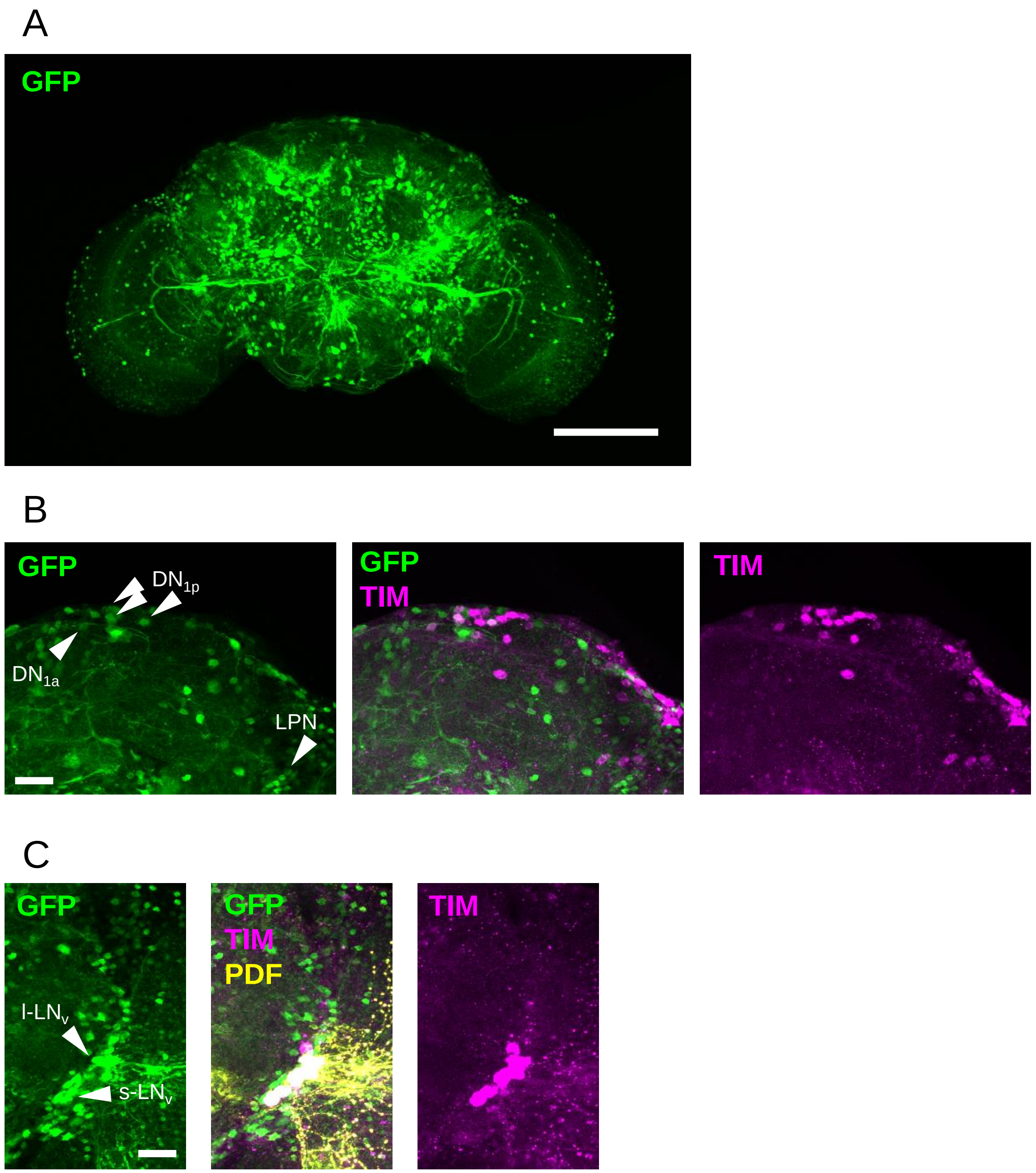

***CNMaR-Gal4***

**$DN_3$**

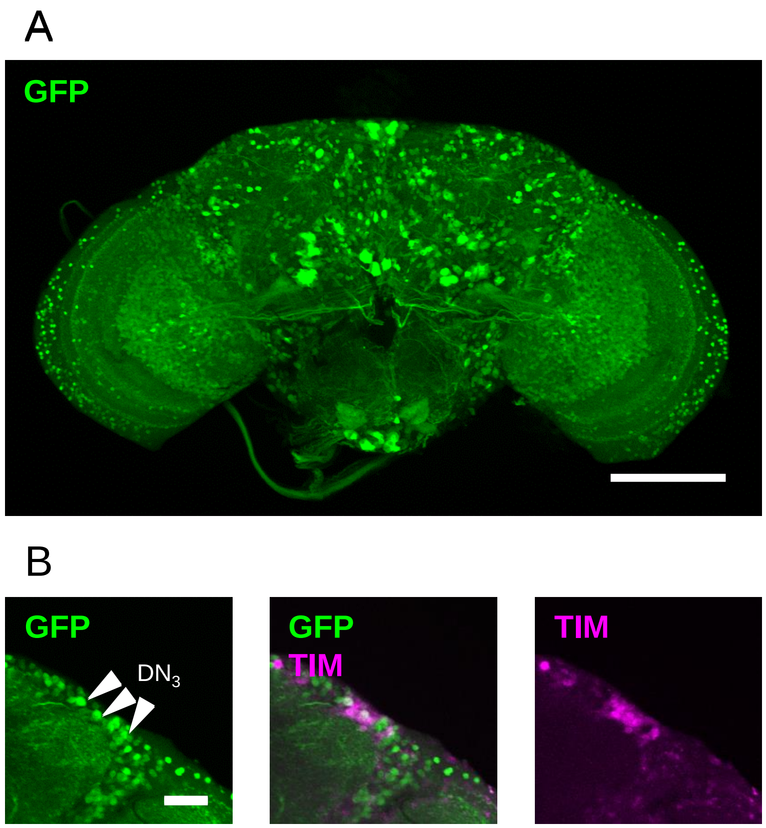

#### *Dh31R-RA/B/C-Gal4*

### l-$LN_v$

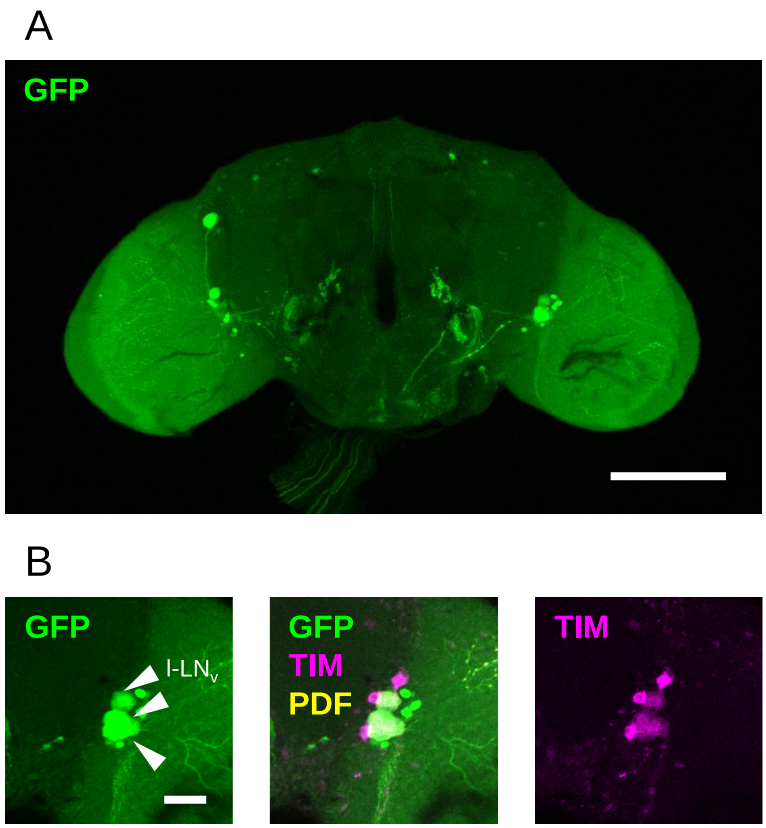

*Dh31R-RC-Gal4*

**l-LN$_v$**

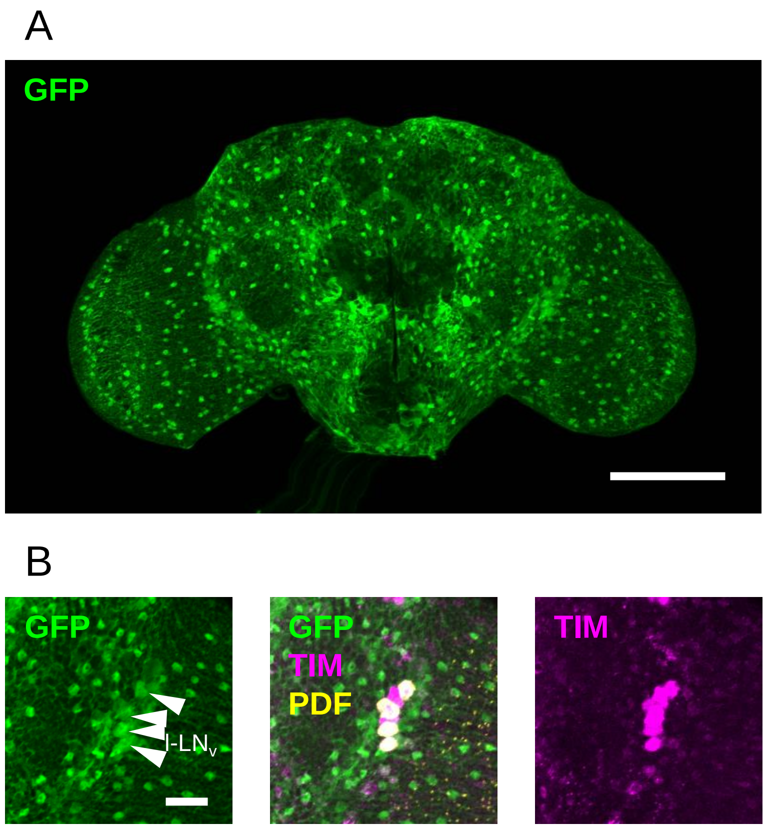

### *NPFR-RA/C-Gal4*

## $DN_3$, $LN_d$, s-$LN_v$, l-$LN_v$

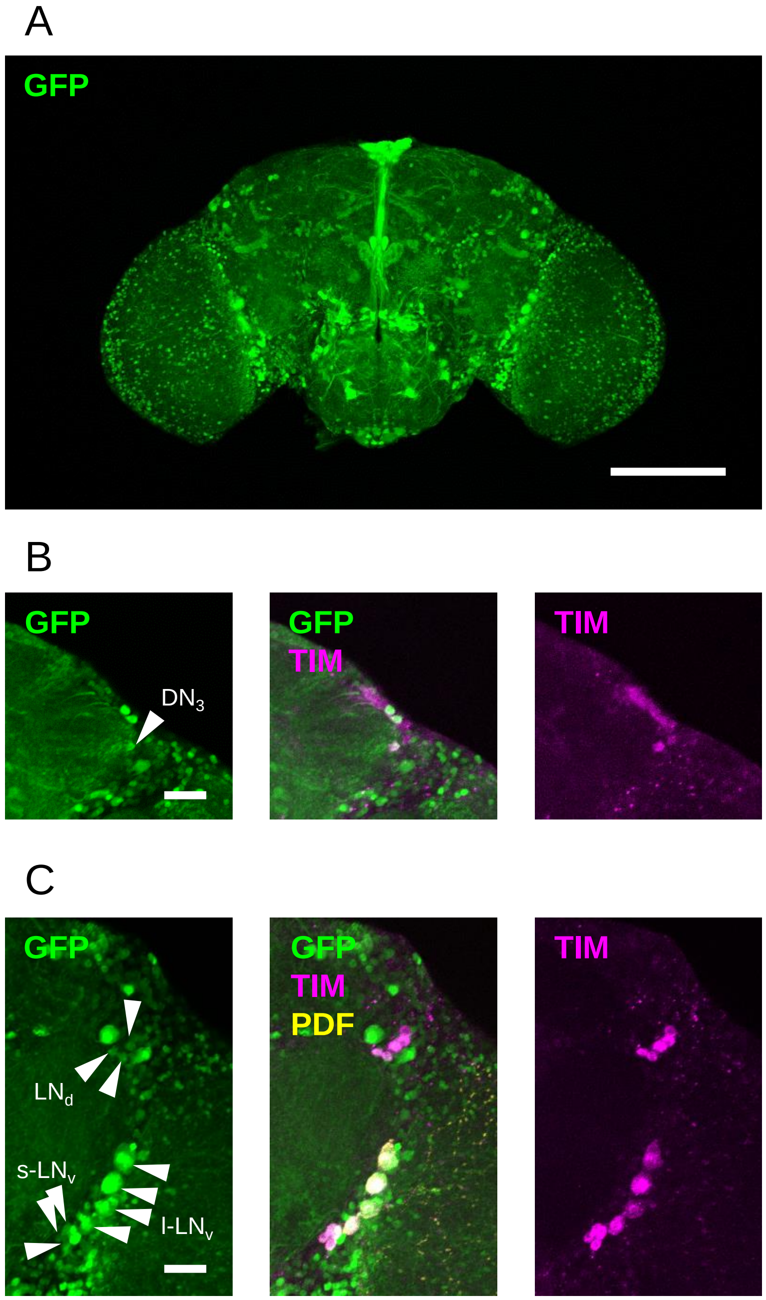

### *NPFR-RB/D-Gal4*

#### l-$LN_v$, LPN

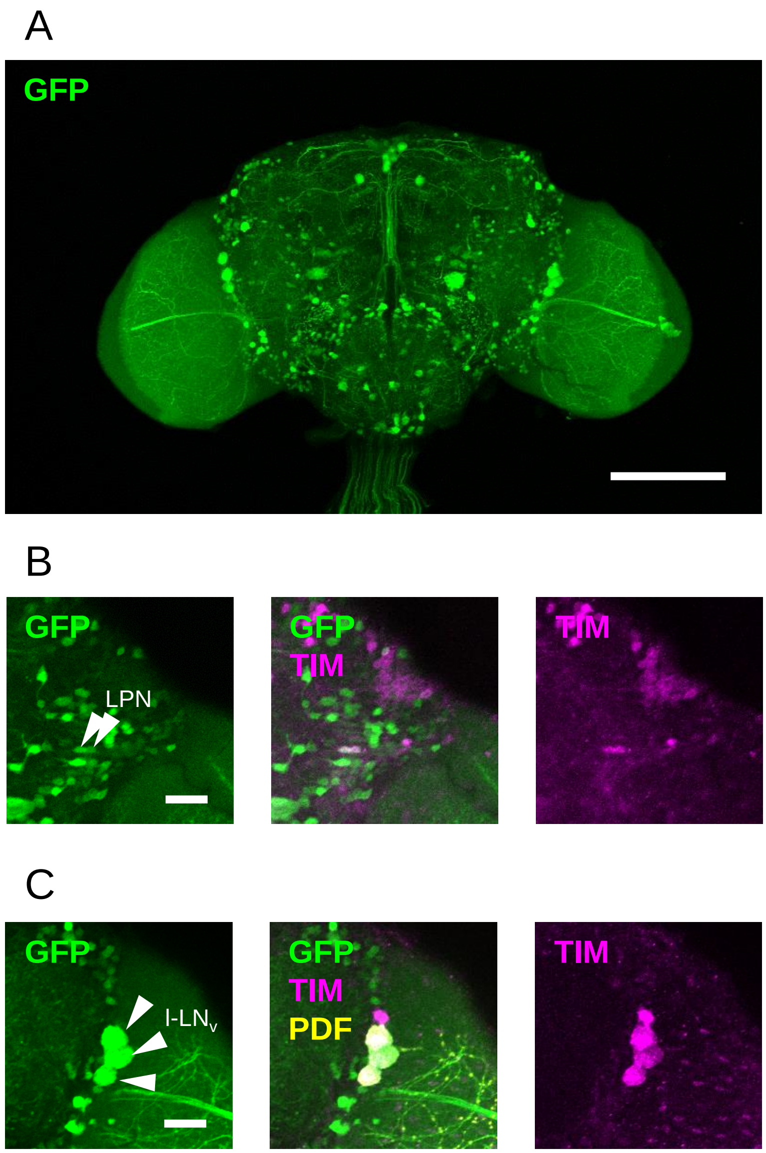

#### *Pdfr-RA-Gal4*

**$DN_{1a}$, $DN_{1p}$, $DN_2$, $DN_3$, LPN, $LN_d$, $5^{th}$-$LN_v$, l-$LN_v$, s-$LN_v$**

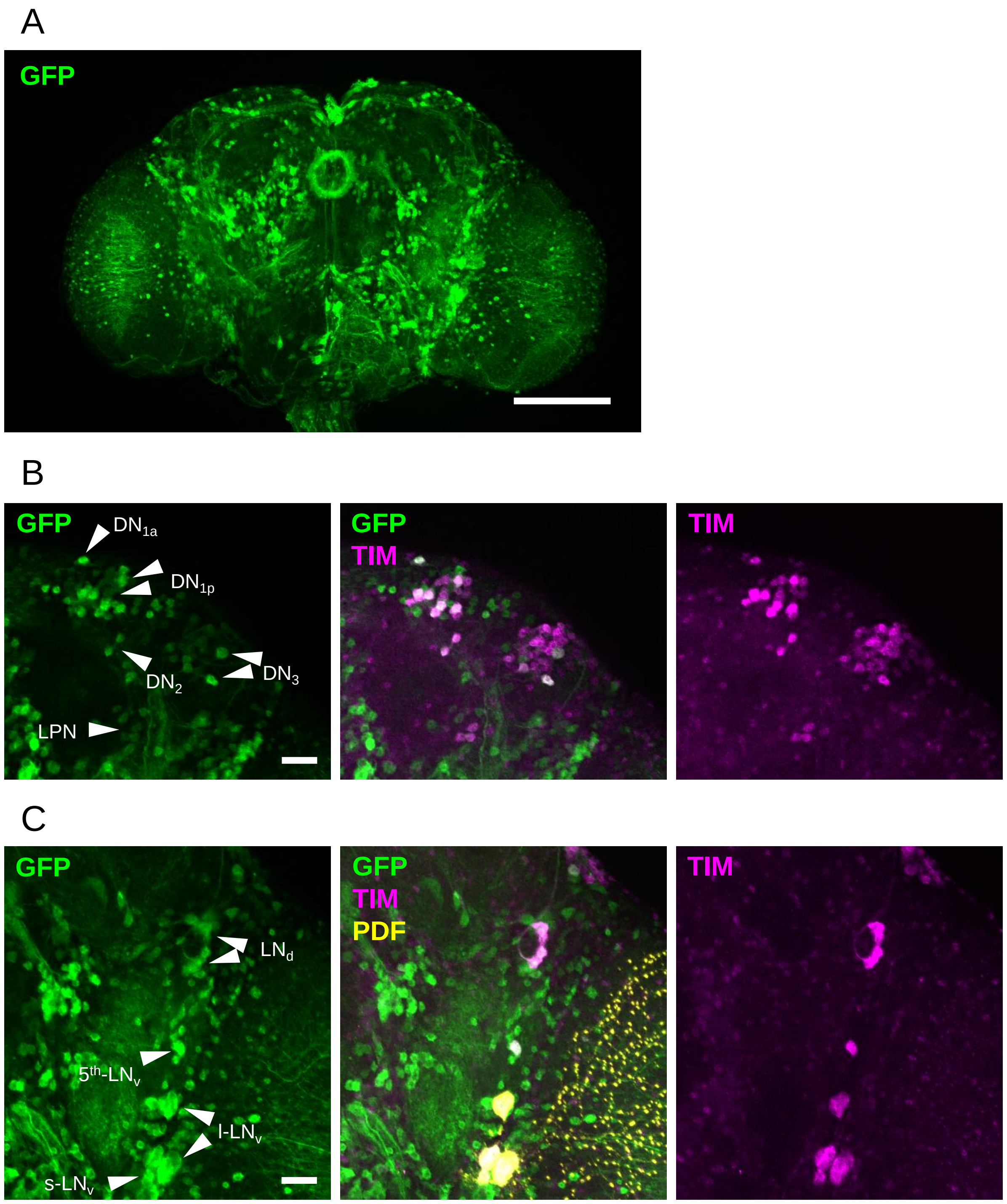

### *sNPFR-Gal4*

#### $DN_{1p}$ $DN_2$, $DN_3$, LPN, $l\text{-}LN_v$

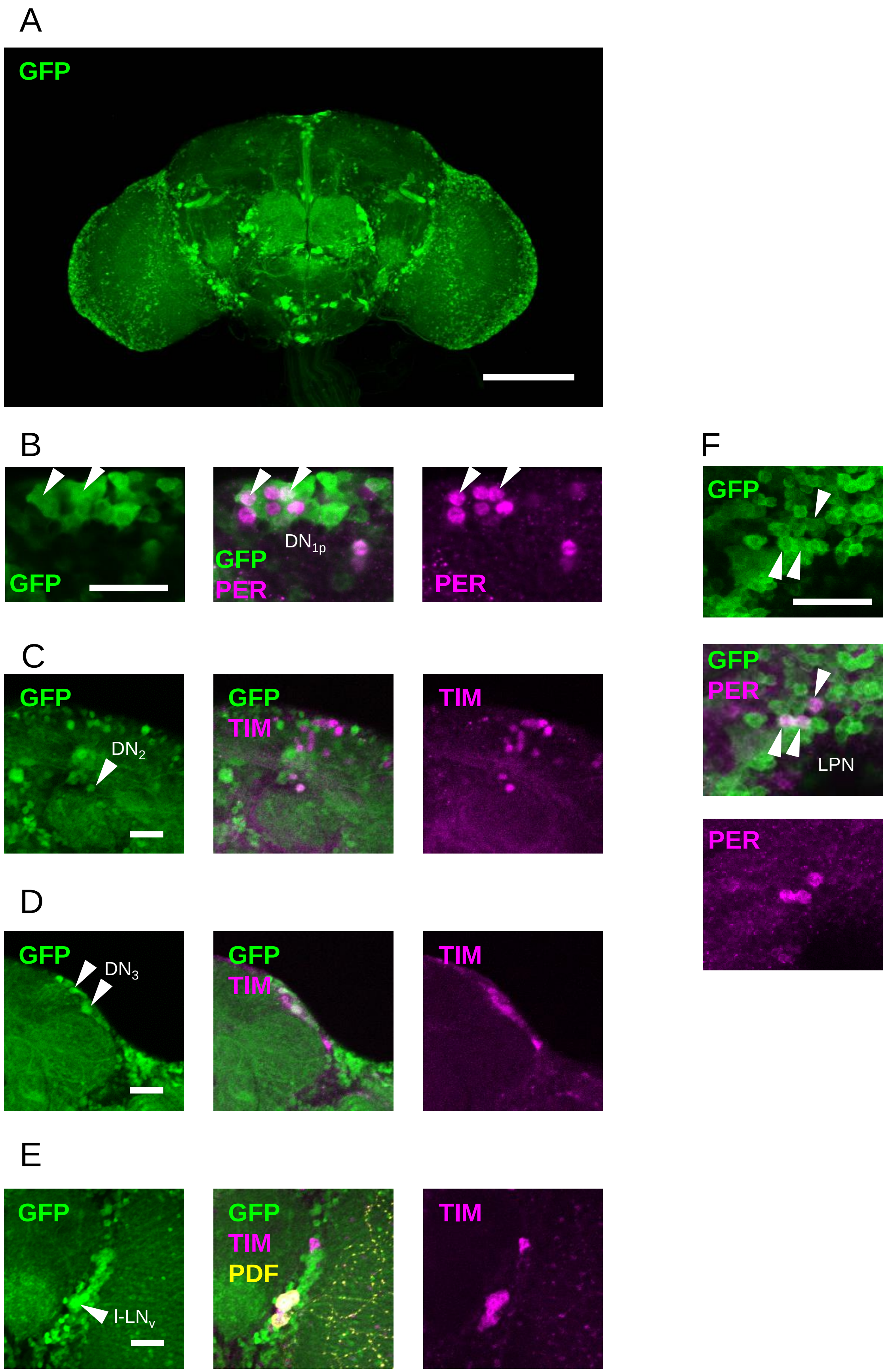
